# Functional diversity of soil microbial communities increases with ecosystem development

**DOI:** 10.1101/2025.02.05.636575

**Authors:** Tord Ranheim Sveen, Maria Viketoft, Jan Bengtsson, Justine Lejoly, Franz Buegger, Karin Pritsch, Joachim Strengbom, Joachim Fritscher, Falk Hildebrand, Ernest Osburn, Emilia Hannula, Mo Bahram

## Abstract

Land abandonment is the single largest process of land-use change in the Global North driving succession and afforestation at continental scales, but assessing its impacts on soil microbial communities remains a challenge. Here, we established a nationwide successional gradient of paired grassland and forest sites to track developments in microbial structure and functioning following land abandonment and gradually changing plant communities. We show that microbes generally respond through threshold dynamics, leading to increasing functional but decreasing taxonomic diversity. Succession also increased the specialization of microbial nutrient (C-N-P) cycling genes while decreasing genetic redundancy, highlighting a putative trade-off between two desirable ecosystem properties: functional diversity and functional redundancy. Increasing fungal functional diversity underpinned higher microbial C-cycling capacity, underscoring the causal link between functional traits and ecosystem processes. Changing litter quality similarly provided a mechanistic link between plant and microbial communities despite otherwise largely decoupled successional developments. Land abandonment is frequently touted as an opportunity to increase biodiversity and carbon storage. Our results show that deeper knowledge about the multifaceted development of soil microbial communities and its links to plant communities during succession may be needed to fully grasp the impacts of global land abandonment processes.

## INTRODUCTION

Land abandonment and afforestation is the single largest process of land-use change in the Global North^1^. Since the onset of industrialization in the mid-1800s, an estimated ∼500 Mha of agricultural land has been abandoned globally^2^, leading to profoundly altered landscapes, ecosystems, and biodiversity patterns^3^. After management ceases, fields are usually colonized by pioneer grasses and herbs, followed by shrubs, bushes and trees, until a stage of full afforestation is reached. For many organismal groups, these changes occur gradually, through directional and largely predictable turnover stages referred to as *succession*^4^, with net biodiversity effects differing depending on organism group and study region^5^. Most afforested sites in Europe are nonetheless ultimately incorporated into plantation-type forest managements^6^, leading to overall negative biodiversity outcomes for a wide range of biota^7–11^. Yet, we still know relatively little about the implications large-scale land abandonment and afforestation has on soil microbial communities.

Soil microbes represent the largest pool of terrestrial diversity^12^, which moreover links mechanistically to gradual changes in resource quality and soil properties during succession^13^. For instance, soil fungal communities typically respond to changing litter quality of plants (resource-driven succession) through increasing leaf-dry matter content (LDMC)^14^, whereas bacteria are more sensitive to abiotic properties like pH and soil organic matter quality (soil C:N)^15^. In contrast to the often predictable patterns observed during plant succession^16^, microbes often show unimodal (i.e. hump-shaped) or threshold responses of abrupt change during succession^17–19^. However, our ability to draw general patterns of microbial community changes during ecosystem development are severely hampered by limitations to the traditional space-for-time designs^20^, which typically fail to adequately reflect super-regional scales of land abandonment^21^. Ecological theory, moreover, posits that niche differentiation and specialization of the decomposer community increase with increasing litter complexity and heterogeneity^22^, and this is accompanied by “tighter” and more efficient nutrient cycling processes^23^. Yet, accumulating evidence shows high functional overlap of microbial genes related to nutrient cycling across differing successional stages^18,24^, suggesting that the overall capacity to perform nutrient cycling remains at similar levels. This aligns with the notion that microbial communities are underpinned by high functional redundancy, whereby multiple taxa share genetic capacities and are, therefore, robust to biodiversity loss^25^.

Here, we established a nationwide gradient spanning from managed and open grasslands to abandoned and increasingly afforested sites undergoing succession (Fig. 1a,c). Each grassland site was paired with an adjacent forest site representing the land-use change endpoint^26^ (Fig. 1b), typically a coniferous-dominated forest managed for wood production. We used amplicon (16S for bacteria, ITS for fungi) and metagenomic sequencing (Fig. 1d), to study how microbial taxonomic diversity and biogeochemical potential developed with abandonment and succession, including links to measured rates of carbon cycling. We hypothesized that increasing resource complexity would increase the functional diversity and specialization of soil microbes, but that high biogeochemical redundancy would maintain stable rates of carbon cycling throughout all successional stages. We, moreover, expected these community changes to occur predominantly as unimodal or threshold-like responses (Fig. 1e). x

**Fig. 1:**
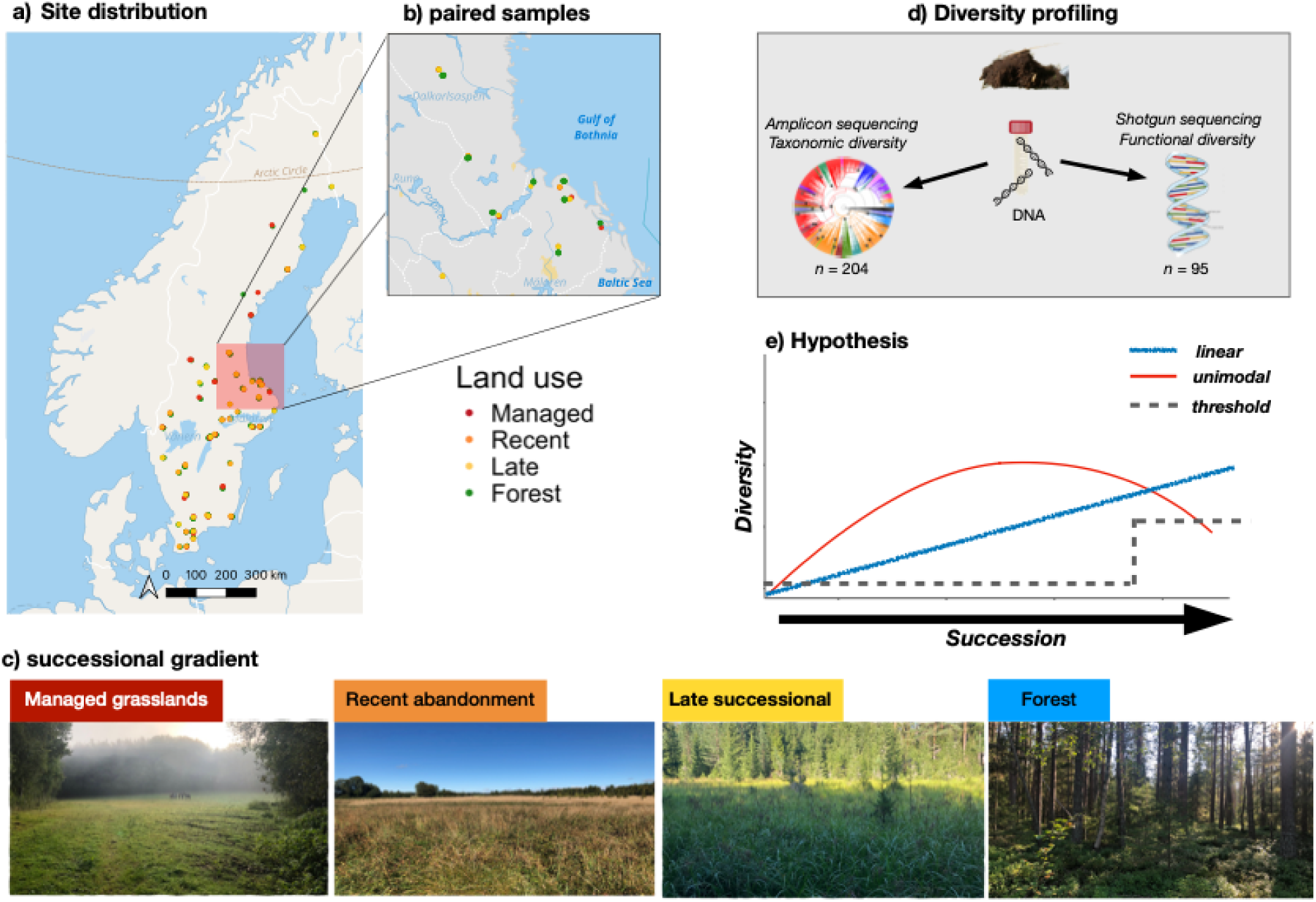
Study design. Distribution of (**a**) sites across a national successional gradient in Sweden, based on (**b**) a paired grassland and forest sites comprising (**c**) managed, recently abandoned, and late successional grasslands together with fully afforested reference sites representing the successional endpoint. Soil samples were gathered in the field and sequenced with (**d**) amplicon and shotgun metagenomics to yield taxonomic and functional microbial community profiles, with the hypothesis that these would change (**e**) linearly, unimodally, or with threshold responses to succession.

## RESULTS

### Threshold effects dominate impacts on soil microbial communities

Our successional gradient revealed a striking decrease of microbial (taxonomic) diversity and marked shifts in community composition with land-use change between grasslands and forests (Fig. 2). By contrast, grassland microbes were overall structurally and functionally similar, independent of the successional stage (i.e. managed, recently abandoned, and late-stage successional grasslands, Fig. 1c) and increasing levels of afforestation (Table S5–6). These patterns are consistent with threshold-like tipping points occurring between late-stage successional grasslands and fully afforested sites (Fig. 1e). Soil bacteria showed the steepest diversity decline (Fig. 2b), whereas fungal responses were more moderate and evened out during the later stage of succession (Fig. 2a). Community composition similarly differed strongly between grassland and forest sites (Fig. 2c-d). In addition, fungal community composition differed between managed and abandoned grasslands for fungal community (Permanova: *p* < 0.05 for all comparisons, Table S6), indicating a compositional shift after the cessation of grassland management. Effect size estimations of the differences between grasslands and forests were overall large for bacteria both in terms of diversity and community composition (partial ω^2^ = 0.46 and 0.19, respectively) and moderate for fungi (partial ω^2^ = 0.07 for alpha diversity and 0.05 for beta diversity, respectively). Notably, the thresholds coincided with a sharp decline in soil pH, increasing soil C:N ratio, and higher levels of leaf dry matter content (LDMC; Table S3). These factors were also consistently identified as the main predictors of microbial change in the variable selection analyses, although the relative importance of each factor varied with diversity aspect (alpha vs beta diversity) and microbial group (Fig. S1). Notably, including distance between paired grassland-forest sites to account for spatial autocorrelation and climate factors (temperature, precipitation) kept the observed patterns of threshold changes the same (Table S5).

**Fig. 2:**
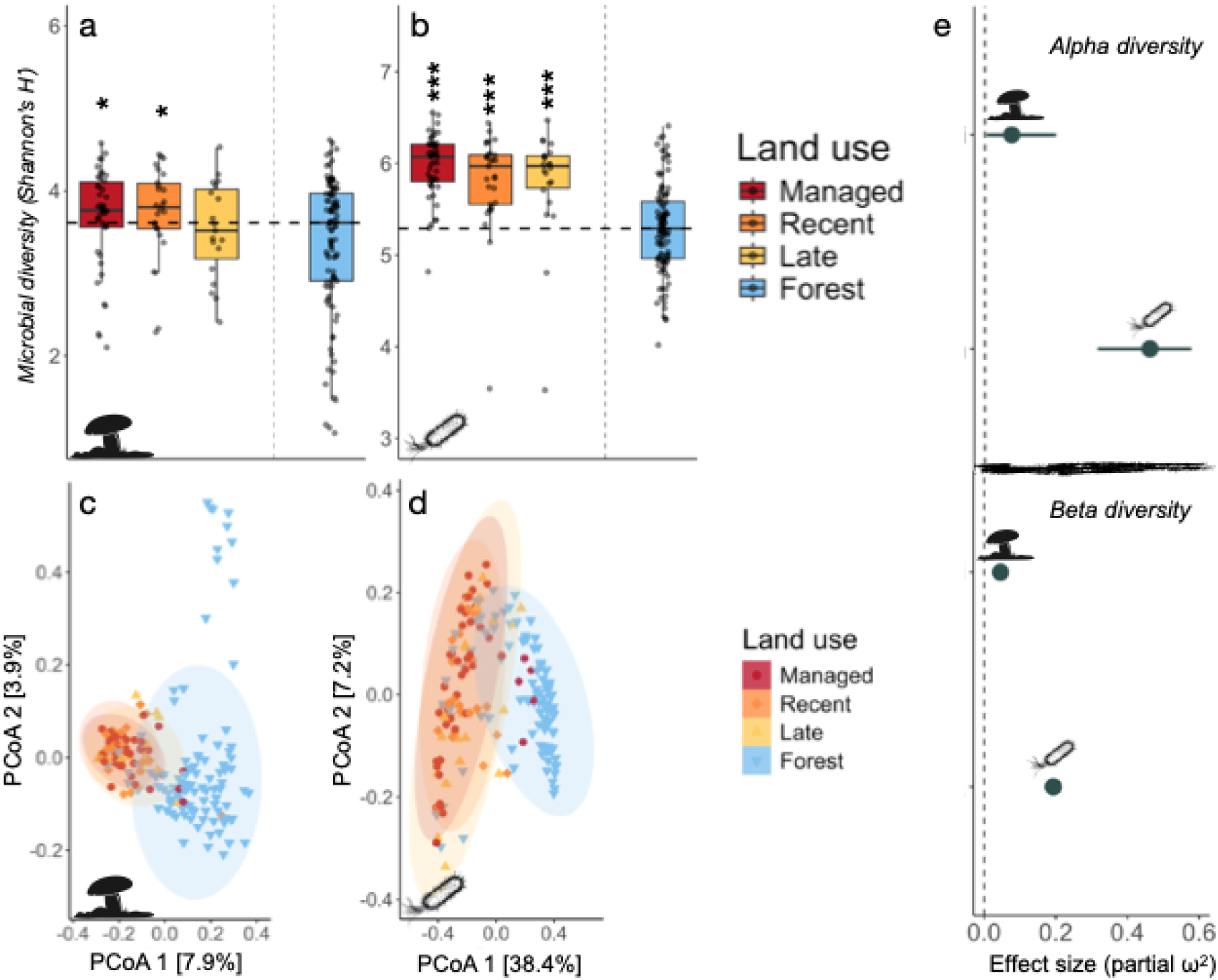
Threshold changes to microbial diversity between grasslands and forests. Taxonomic diversity (Shannon’s *H’*) of **a**) fungal, and **b)** bacterial communities across the land abandonment gradient. The lower and upper hinges of the boxplots represent the 25^th^ and 75^th^ percentiles, respectively, and the middle line is the median. The whiskers extend from the median by 1.5x the interquartile range. The dashed horizontal line indicates the median of the forest sites, with asterisks (*) denoting significant differences between paired grassland and forest sites based on mixed-effect linear models accounting for paired structure and spatial distance with the following significance levels: \**p* < 0.05, \*\**p* < 0.01, \*\*\**p* < 0.001. Full test details are listed in Supplementary tables S5–S6. Principle coordinate analyses (PCoA) of **c**) fungal, and **d**) bacterial community composition during succession based on bray-curtis distances. Results from pairwise permutational multivariate tests (perMANOVA) for difference in community composition between land-use stages are found in Supplementary tables S7. **e)** Differences between grasslands and forest alpha and beta diversity for bacterial and fungal communities based on estimated partial ω^2^ effect sizes ±95 CI.

### Functional diversity increases relative to taxonomic diversity during land abandonment

We next examined how the microbial genetic potential for biogeochemical cycling (i.e. C, N, P-related genes) changed with ecosystem development. Also, here, we found no differences between grasslands across differing stages of management and succession (*p* > 0.05 for all pairwise comparisons; Table S8). Conversely, threshold effects were notable in the response of fungal C-cycling genes, which increased in forests compared to grasslands (Fig. 3a; partial ω^2^ = 0.50). By contrast, the diversity of bacterial C-cycling and all microbial P-cycling genes remained at similar levels across all land uses (Fig. 3b-d). Bacterial N-cycling genetic diversity decreased in forests compared to grasslands (Fig. 3e; partial ω^2^ = 0.54). When juxtaposing the results obtained from the taxonomic and functional profiling (Figs. 2-3), a pattern of increasing functional relative to taxonomic diversity merged, indicative of changing microbial redundancy and functioning.

**Fig. 3:**
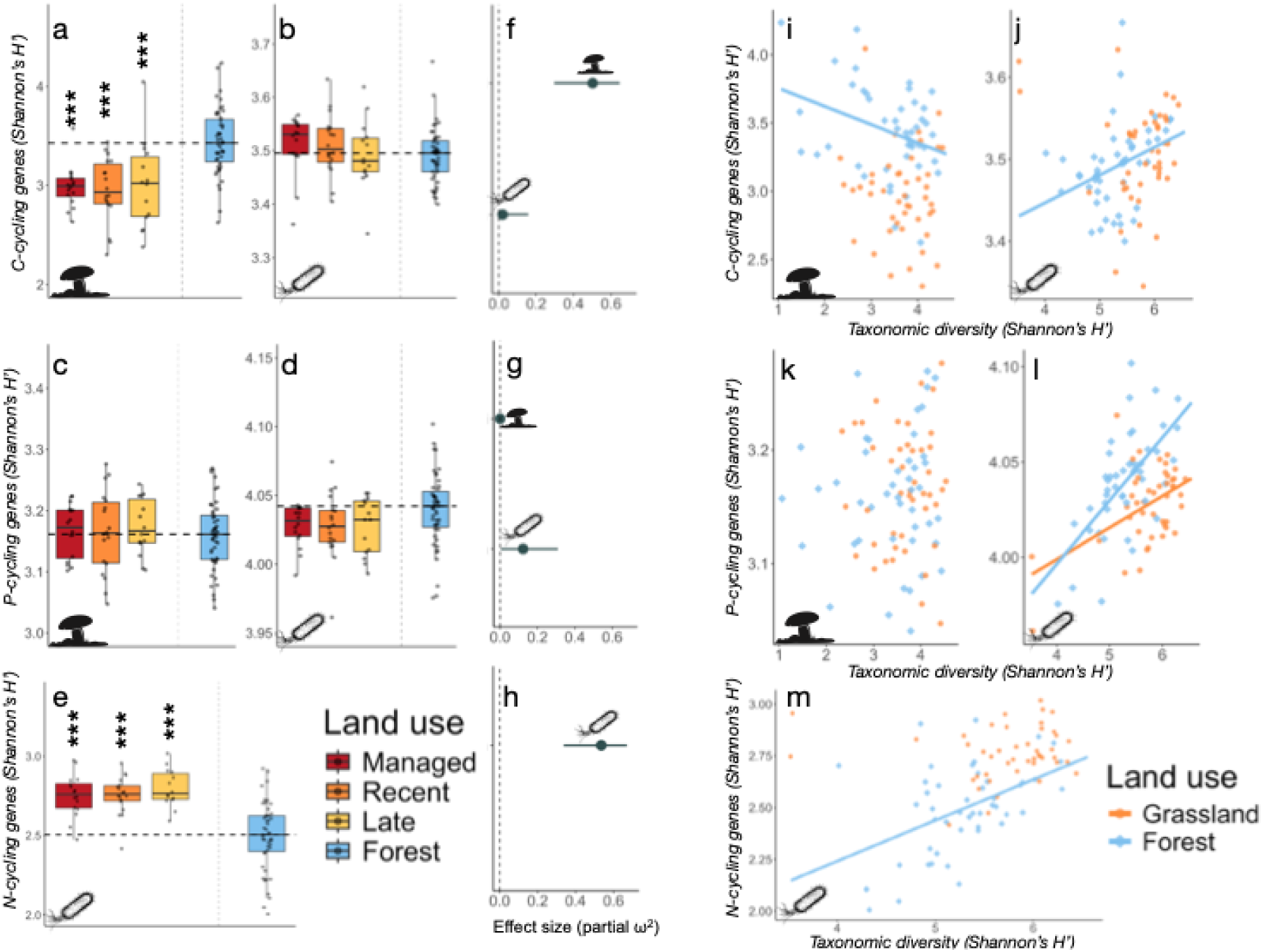
Genetic functional redundancy decreases with change from grasslands to forests. Shannon diversity of genes related to **a-b**) C-cycling, **c-d**) P-cycling and **e)** N-cycling across the land abandonment gradient, along with their respective effect size estimates based on estimated partial ω^2^ effect sizes ±95 CI. (**f-h**). The lower and upper hinges of the boxplots represent the 25^th^ and 75^th^ percentiles, respectively, and the middle line is the median. The whiskers extend from the median by 1.5x the interquartile range. The dashed horizontal line indicates the median of the forest sites, with asterisks (*) denoting significant differences between paired grassland and forest sites based on mixed-effect linear models accounting for paired structure and spatial distance with the following significance levels: \**p* < 0.05, \*\**p* < 0.01, \*\*\**p* < 0.001. Full test details are listed in Supplementary tables S8–S9. Scatter plots (**i-m**) showing the relationship between taxonomic and genetic diversity for fungal and bacterial communities. Solid lines indicate significant (*p* < 0.05) fit based on ordinary least-square regression. Full regression statistics are listed in Supplementary table S10.

### Loss of biogeochemical redundancy with grassland afforestation

The absence of direct relationships between taxonomic and functional diversity is considered a strong indicator of functionally redundancy, as taxa can be replaced without losing genetic potential^27,28^. To assess this important aspect of ecosystem stability^29^ more closely, we first combined all grassland sites into one single grassland category as the differences between successional stages were minimal (Figs. 2-3). Next, we used ordinary least-square regression (OLS) to examine the strength and direction of the relationship between taxonomic diversity (predictor) and functional diversity (response variable). For fungi, the marked increase of C-cycling genes in the forest sites occurred despite a general loss of taxonomic diversity (Figs. 3a vs 2a), producing a negative relation (OLS: estimate = −0.13, *p* = 0.022; Fig. 3i) which was not significant in grasslands (OLS: estimate = −0.04, *p* = 0.613). Comparing the slopes of both ecosystems showed that these differed significantly (Anova: F_1,86_ = 42.6, *p* < 0.001), indicating that the coupling of taxonomic diversity and genetic C-cycling potential undergoes substantial alteration as grasslands transition to forests. Similar results were obtained when assessing bacterial C-cycling and N-cycling genetic diversity (Fig. 3j,m). Both these genetic pools were moreover significantly related to taxonomic diversity in forests but not in grassland soils (OLS, *p* < 0.05 for both comparisons; Table S10). Importantly, the average genome size (AGS) of bacterial communities was included as a fixed effect in all models assessing bacteria but did not alter the direction or significance of the taxonomic-functional diversity relationships (Table S8). For P-cycling, diversity coupling was significant in both ecosystem types for bacteria (Fig. 3l), albeit stronger in forests than in grasslands (Anova, F_1,87_ = 34.2, *p* < 0.001). Interestingly, no corresponding coupling was found for fungi (Fig. 3k), indicating that functional redundancy may be relatively high when it comes to fungal P-cycling independent of ecosystem development. Soil pH and carbon quality were the main factor related to genetic C-cycling diversity for both fungi and bacteria, whereas P-cycling diversity was related to the total and mineral N content for fungi and bacteria, respectively (Fig. S2).

### Widespread specialization of genetic nutrient cycling potential during ecosystem development

To examine whether the functional specialization of microbial communities increases after land abandonment, we quantified the average niche overlap of C-N-P-related genes among successional stages (see methods). Results revealed a gradual specialization across the entire soil microbial communities during ecosystem development (i.e. both fungi and bacteria) for C- and P-cycling, as evidenced by lower levels of genetic overlap going from managed grasslands to forests (Fig. 4a-d). In contrast, overlap of N-cycling genes, calculated for bacteria only due to poor genetic annotation of fungal N-cycling genes, showed a weakly unimodal pattern peaking in early successional grasslands (Fig. S3). When partitioning the genetic overlap across differing substrates of increasing carbohydrate complexity (C-cycling; Fig. 4f-g) and pathways (N-P-cycling; Figs. S3–4), we also found that specialization increased across the full spectrum of simple and complex substrates. Notably, P-cycling specialization increased gradually for fungi, whereas the overlap between many of the P-cycling pathways

**Fig. 4:**
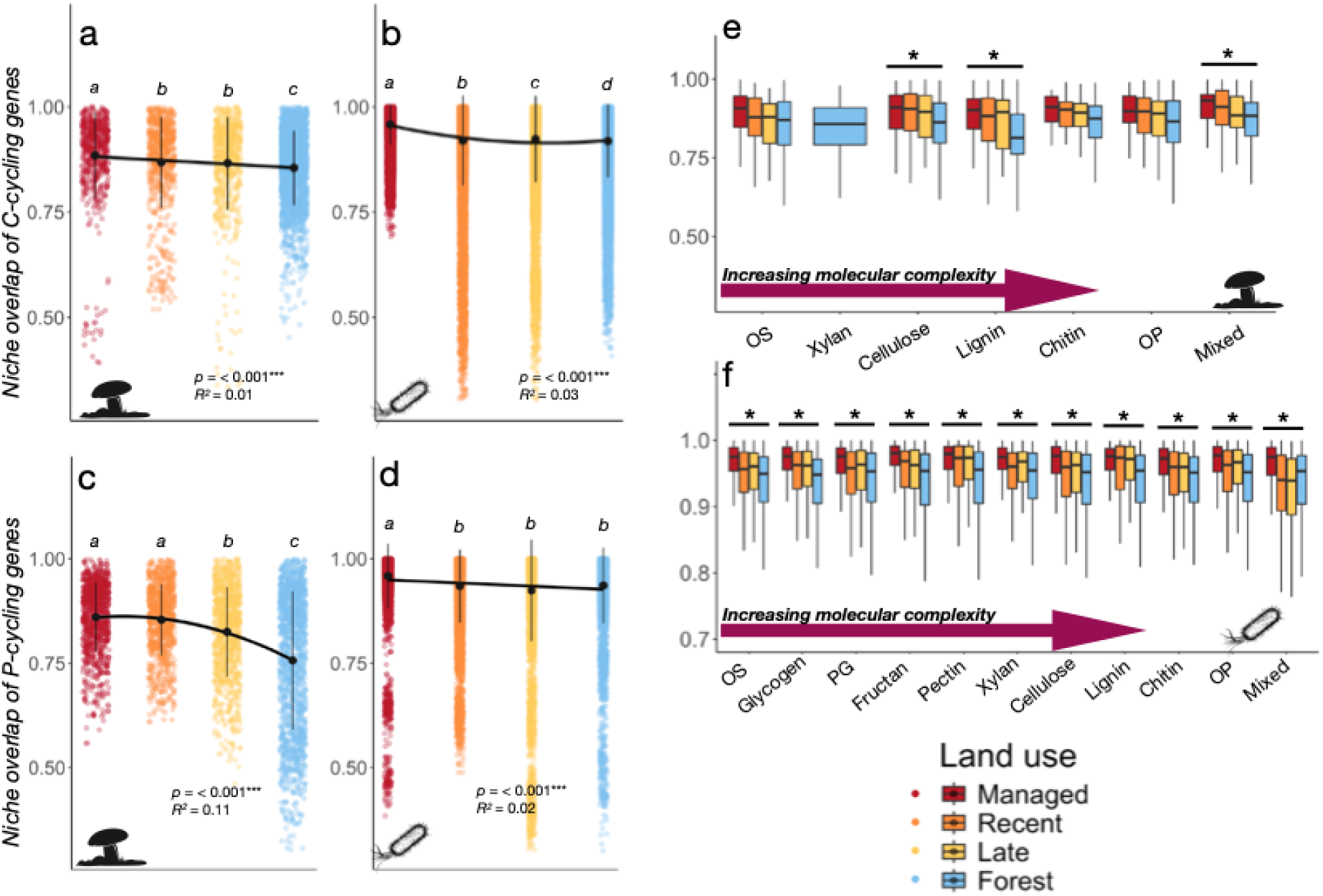
Community-wide genetic specialization of microbial nutrient cycling increases with succession. Niche overlap of carbon cycling genes for **a**) fungal and **b**) bacterial communities across the land abandonment gradient. Panel **c-d**) show corresponding niche overlap of P-cycling genes. An additional figure showing bacterial N-cycling genes is found in the Supplementary Fig. 3. Black points show mean genetic overlap ± s.d. (vertical lines) for each land-use stage. Horizontal lines connecting land-use stages indicate significant (*p* < 0.05) ordinary least-square or second-order polynomial regression fits (Table S12). Letters indicate significant differences (*p* < 0.05) between land use-stages based on pairwise Wilcoxon Rank-Sum Tests (Table S13). In panels **e-f**) the C-cycling have been partitioned according to their C substrate class and displayed for **f)** fungi and **g)** bacteria across a gradient of increasing molecular complexity. Stars (*) above these show substrates where differences between one or more land-use stages are significant (*p* < 0.05) based on pairwise Wilcoxon tests, with full test results found in Table S14. Corresponding boxplots for P-cycling pathways are found in Supplementary Data Fig. 4. OS = Oligosaccharides; PG = Peptidoglycan; OP = Other polysaccharides; Mixed = Mix of oligo- and polysachhcarides decreased sharply between managed and abandoned grasslands for bacteria (Fig. S4).

### A diversity-redundancy framework for soil microbial communities during succession

As our results strongly indicated that a land-use change from grasslands to forests entail a shift from communities characterized by functional redundancy to increasing functional specialization, we formalized and tested this trade-off using the framework developed in refs^30,31^. Briefly, by combining functional and taxonomic dimensions of a community, three interrelated properties describing taxonomic diversity, functional uniqueness (hereafter used synonymously with specialization) and functional redundancy can be derived and plotted along an axis of redundancy and specialization (Fig. 5a). Yet, as these analyses require traits-by-taxa matrices, we first predicted bacterial metagenomes from amplicon sequences (*n* = 204) using *picrust2*^32^, and then inferred C-N-P-cycling genes based on KEGG orthology (see methods). With the predicted metagenomes, we assessed redundancy and specialization under the hypothesis that grasslands and forest sites would cluster differentially along the vertical axis separating these diversity dimensions. As expected, functional redundancy related to C-cycling decreased sharply between grasslands and forests, whereas functional specialization instead increased (Fig. S5), consistent with a threshold response to afforestation occurring late in the successional gradient. The quantity (Total C) and quality (C:N) of organic carbon were the main variables driving redundancy and specialization (Fig. S6a), differing from the factors affecting bacterial alpha diversity and community composition (Fig. S1). Clustering of grassland and forest sites occurred mainly along the redundancy-specialization axis, in line with our hypothesis (Fig. 5c; Fig. S7), and these results were further corroborated by compositional differences between the two land-use types (Table S17). Notably, N- and P-cycling showed the same patterns of a redundancy-specialization trade-off (Fig. S7), whereas no differences were found between any of the differing grassland successional stages. However, the main factors underlying this trade-off varied, with N-cycling more related to soil pH in addition to soil carbon quality (Fig. S5b) whereas P-cycling was related to the successional stage of the site and plant leaf traits (Fig. S5c).

**Fig. 5:**
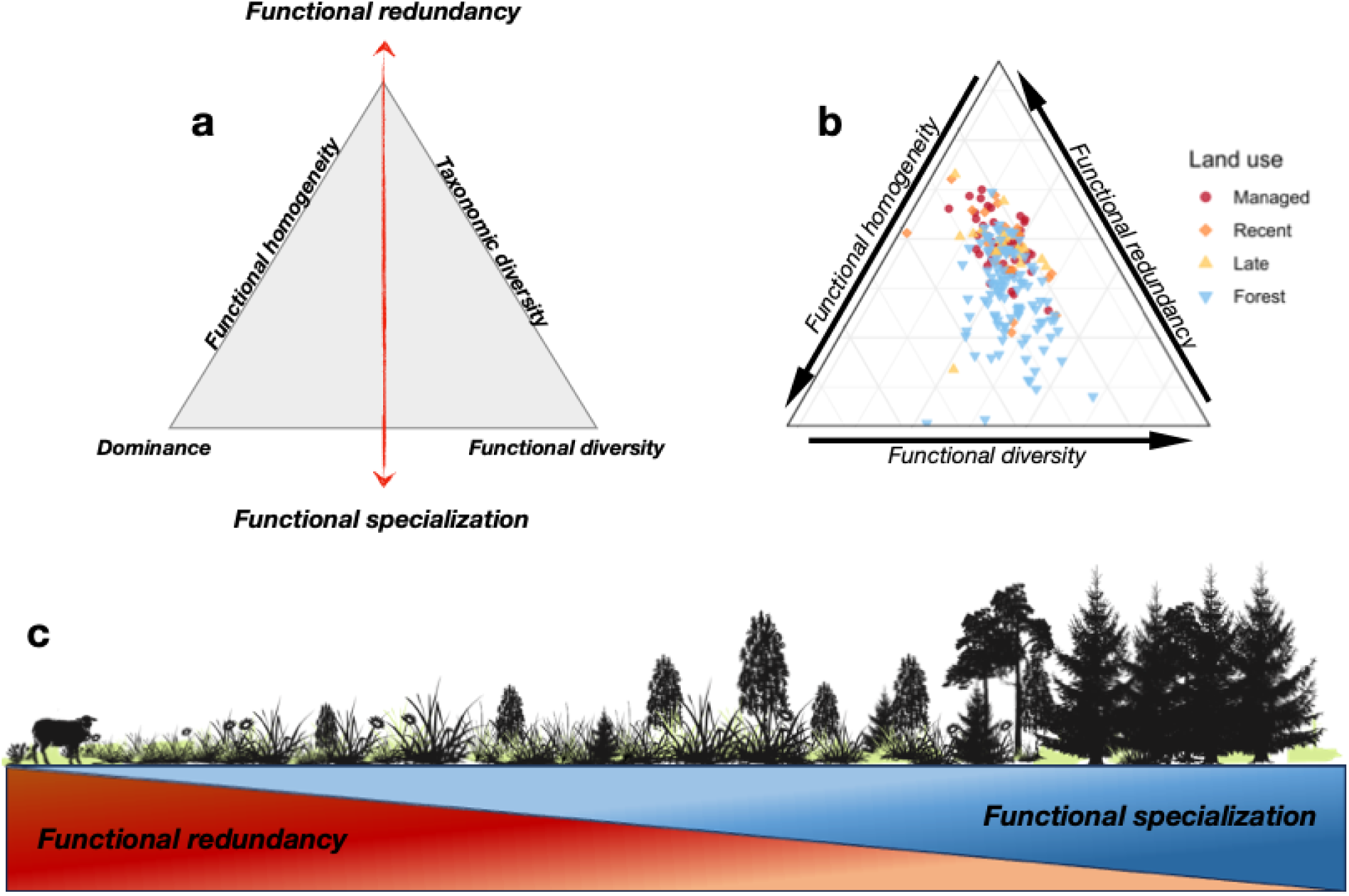
A general framework of shifting microbial redundancy-diversity during succession. Conceptual ternary plots showing **a**) interrelated components of taxonomic and functional diversity and how these can be arranged conceptually to produce an axis of functional redundancy-functional specialization. Ternary plot **b**) showing the placement of grassland and forest sites according to the diversity metrics shown in panel (a) of bacterial communities inferred from predicted metagenomes. Corresponding ternary plots for P- and N-cycling can be found in the Supplementary Data Fig. 7. See refs^30–31^ for a mathematical and conceptual elucidation of the interrelated diversity components shown in panel **a**. Conceptual gradient **b**) depicting the redundancy-specialization axis during succession after land abandonment.

### Functional diversity a key driver of microbial carbon cycling capacity

Lastly, we tested whether increasing functional diversity and specialization during succession would affect carbon cycling rates across a gradient of increasing substrate complexity. For this, we incubated a subset of paired grassland-forest samples (*n* = 154) and measured substrate-induced respiration (SIR) across six substrate types ranging from easily degradable oligosaccharides (glucose) to complex recalcitrant substrates like lignin and chitin. C-cycling rates were generally higher in forest than in grassland soils but increased with abandonment and succession (Fig. 6a). Interestingly, the least and the most recalcitrant substrates (glucose, chitin) were exceptions to this, with no differences between any of the grassland and forest soils (Table S18). When combining the full range of respiration rates into an aggregate measure of carbon cycling capacity (MSIR, see methods) and using this as a response variable with microbial taxonomic and functional diversity as predictors, we found that fungal C-cycling genetic diversity was key to the overall community’s carbon cycling capacity (Fig. 6b). Notably, this relationship did not differ between grasslands and forests (Anova: F_1,70_ = 0.015, *p* = 0.903), suggesting high fungal control over decomposition in across the full range of ecosystems undergoing land abandonment. In contrast, we found no corresponding relationship for bacterial C-cycling genetic diversity (Fig. 6c). MSIR moreover correlated negatively with bacterial taxonomic diversity (*r* = −0.37, df = 143, *p* < 0.001) and showed no significant relationship with fungal diversity (*r* = −0.06, df = 142, *p* = 0.49).

**Fig. 6:**
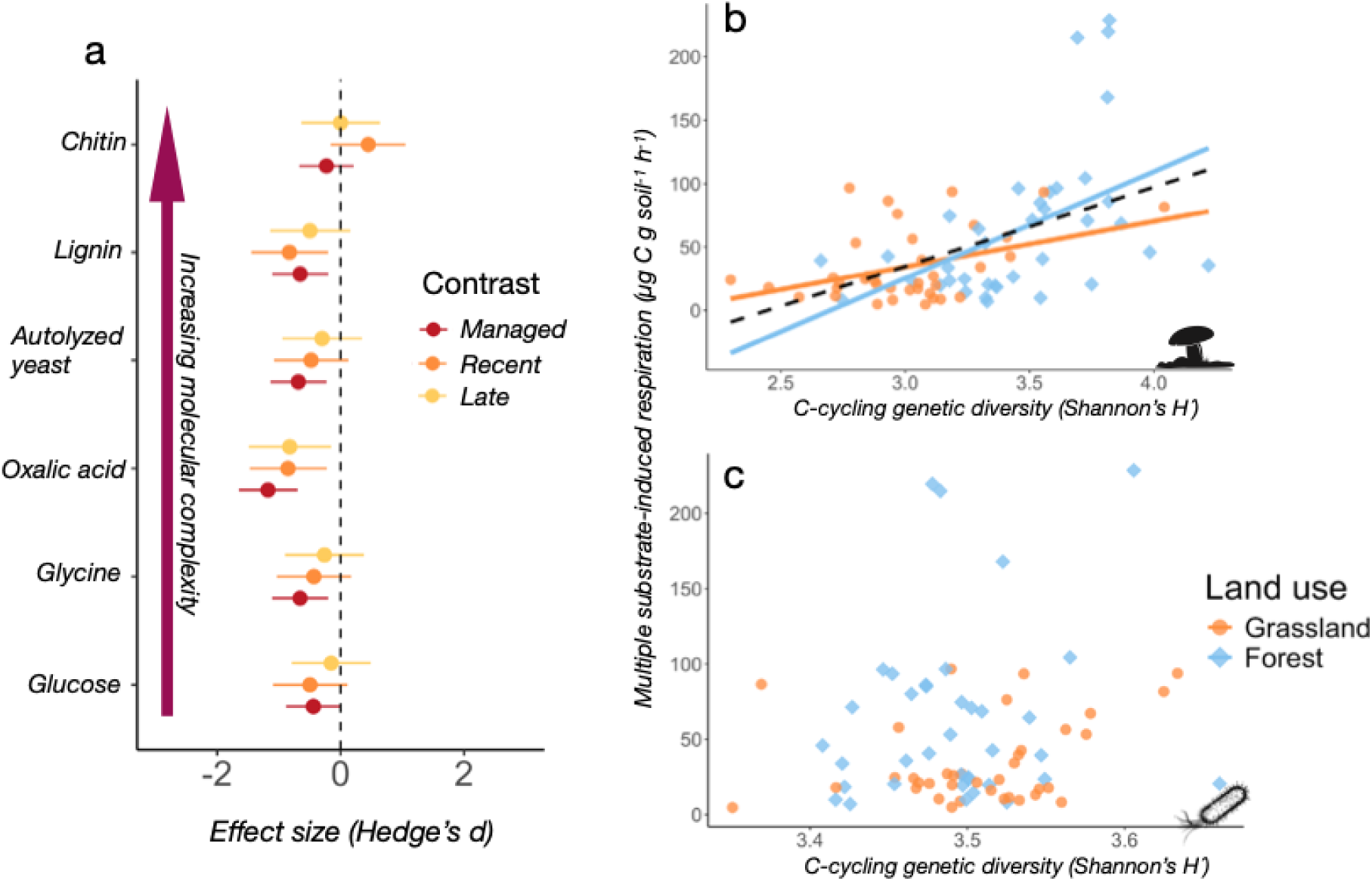
Fungal functional diversity drives carbon cycling capacity in grassland and forest soils. **a)** Effect sizes of substrate-induced respiration (SIR) rates between paired grassland and forest sites for managed, recently abandoned and late successional grasslands. Points to the left of the dashed line indicate higher respiration rates in forests than in grasslands, whereas points to the right of the dashed line indicate higher grassland respiration rates than forest respiration rates. Lines denote 95 % confidence intervals. Scatter plots showing the relationship between **b**) fungal, and **c**) bacterial genetic C-cycling diversity and the aggregate respiration rate of all substrates (MSIR). Solid lines indicate significant (*p* < 0.05) fit of the regression line for each land-use type (grassland, forest). The dashed line in panel **b**) shows the overall fit of all points independent of land-use type.

## DISCUSSION

Elucidating the patterns and processes occurring after land abandonment is a notorious challenge in ecology because the gradients used to infer ecosystem development are spatially confined and fail to match the extent of land abandonment. By establishing a nationwide successional gradient of paired grassland and forest sites (Fig. 1), we could examine how microbial communities respond to land abandonment at an unprecedented scale. We found that these responses predominantly occurred as thresholds between the late-stage successional grasslands and the fully afforested sites, contrasting with foundational successional theory, where succession is thought to occur as a gradual and largely predictable series of changes^16,33^. Our results moreover suggest that these threshold dynamics dominate multiple microbial community properties such as taxonomic diversity (Fig. 2), genetic diversity (Fig. 3), and, by extension, also functional specialization (Figs. 4-5) and functional redundancy (Fig. 3; Fig. 5). While it was not possible to locate the exact threshold location based on our study design, we note that it coincided with a sharp decline in soil pH, increasing soil C:N, and higher levels of leaf dry matter content (Table S3), all well-known changes in soil conditions after land abandonment in boreal biomes^14^. This aligns well with soil pH, C:N, and leaf properties (LDMC) consistently singled out as main drivers of both taxonomic and functional diversity in our variable selection analyses (Figs. S1–2), and with previously highlighted mechanisms of microbial change such as rapid depletion of base cations through tree growth leading to soil acidification^34^ and a build-up of recalcitrant leaf litter requiring complex enzymes to decompose^35^. Plant-microbial interactions are seen as a fundamental driver of succession and ecosystem change^13,36,37^, but our findings paint a contrasting picture of high microbial community resilience and functional redundancy to significant levels of afforestation and plant community change. Similar results from biodiversity-rich meadows in Central Europe also show substantial plant-microbial decoupling during abandonment^38^, suggesting that microbial responses to abandonment may be unified across even larger spatial scales than the ones examined here. This would also align with recent findings showing unified soil microbiome responses to climatic extreme events across continental scales^39^. Yet, predicting how individual groups of microbes respond to abandonment would require substantial knowledge about local site conditions, including more detailed knowledge about the thresholds and tipping points in which also microbial communities undergo abrupt and potentially irreversible changes^19^.

The absence of a direct positive or negative relationship between taxonomic and functional diversity is indicative of substantial functional redundancy^27^. We found differences in this relationship between grasslands and forests, with notably higher functional diversity relative to taxonomic diversity in forest soils (Fig. 2-3). This suggests that fewer taxa contribute to the overall pool of genetic diversity in forest soils and is corroborated by the generally strong correlations between taxonomic and functional diversity in these sites but not in grassland (Table S11). Importantly, these relationships were independent of the communities’ average genome size (AGS; Table S10), suggesting that a shift toward taxa with larger genome size is not the main driver behind the loss of bacterial redundancy in forest soils^40^. By contrast, fungal genome sizes increase in nutrient-poor soils^41^, and it is therefore plausible that the compositional turnover of fungi during succession is driven by selection for taxa with larger genomes. Although this hypothesis could not be tested explicitly due to the lack of tools to reliably infer AGS for fungal communities, it could explain the negative relationship observed here between fungal taxonomic and C-cycling genetic diversity (Fig. 3i). We also note the generally negligible influence of climate (mean annual temperature and precipitation) on our results (Fig. S1), indicating consistent microbial successional dynamics across differing climatic zones. The generally decreasing overlaps in the genetic repertoires of both fungal and bacterial communities with succession moreover indicate that specialization occurred in all measured aspects of nutrient cycling (Fig. 4a-d; Fig. S3) and across all pathways (Fig. 4e-f, Figs. S3–4). This is in line with recent evidence showing that microbial specialization along one axis of environmental variation is generally paralleled by specialization across other niche axes^42^, leading to widespread multispecialization. Complex substrates enriched in recalcitrant lignin and chitin typically dominate the litter-derived pool of organic matter in forest soils^43^, and the enzymatic degradation of these into progressively simpler molecules selects for a highly specialized web of microbial cross-feeding^44^. By contrast, grassland soils are characterized by simpler and more easily degradable carbon inputs, in which metabolic generalists with broad enzymatic repertoires dominate the decomposer community^45^. Crucially, the use of substrate-induced respiration to measure the soils’ carbon-cycling capacity suggests that this was primarily driven by the functional diversity of the fungal communities (Fig. 6), highlighting the presumably strong causal link between functional specialization and nutrient cycling efficiency observed recently in the context of N-cycling^46^.

Functional diversity and functional redundancy are both crucial components supporting ecosystem functioning and stability^29,47,48^. Yet, our results indicate that these two diversity components may be inversely interrelated and hence trade-off during succession (Fig. 5c). This idea has been previously proposed in the context of global bird, mammal, and plant distributions^31,49^, but to our knowledge, it has not been evaluated and tested for soil microbes. Using computationally predicted bacterial metagenomes, we found that grassland and forest sites clustered regularly along the redundancy-specialization axis for all nutrient cycling processes (Fig. 5a; Fig. S7), and that this trade-off was likely driven by changing soil C:N, pH, and LDMC (Fig. S6). This corroborates our hypothesis that changes in resource complexity and soil acidity trigger shifts from taxonomically diverse but functionally redundant (i.e. more similar with respect to functioning) to functionally rich communities with more specialized communities. An intriguing outcome of this trade-off is that ecosystems undergoing afforestation after land abandonment may become more efficient when it comes to carbon- and nutrient cycling due to higher functional diversity and specialization, but less resilient to disturbances due to lower functional redundancy^29^. We note similar findings from other environmental gradients showing negative relations between taxonomic and functional diversity^40,50,51^.

Lastly, we note that the designation of conventionally managed forest sites as the successional endpoints in this study reflects a deliberate choice to study the outcomes of land abandonment processes in Sweden and Europe more generally. An estimated 75 % of all forests in Sweden and Europe are moreover in even-aged stands^52^, which strongly suggests that abandoned land, once approaching full afforestation, are incorporated into conventional forestry regimes comprising monoculture plantations^7^ and clear-cut felling^53^. Using conventionally managed but adjacent forest sites represents the best possible reference for the ultimate fate of abandoned grasslands, as opposed to any putatively undisturbed “climax” forest (sensu Clemens^54^), which is considerably more scarce.

## CONCLUSION

Our results reveal high microbial community resilience and functional redundancy during land abandonment and gradual afforestation, followed by increased functional diversity and specialization in fully afforested reference sites. These dynamics imply threshold responses and a ubiquitous trade-off between two critical components of ecosystems, functional redundancy and functional diversity, as ecosystems develop after abandonment. Importantly, these trade-offs occurred in all aspects of nutrient cycling and may affect ecosystem functioning and resilience. Further studies are needed to investigate the generality of these trade-offs and their implications for long-term ecosystem stability.

## ONLINE METHODS

### Study design

The study comprised 102 grassland and 105 forest sites (Table S1), of which 190 (92 %) were paired based on a geographical proximity criterion (< 6.5 km distance between paired sites, median distance = 3.63 km). Grasslands were initially categorized into differing successional stages (Managed, recently abandoned, Late successional) based on site descriptions, and this was corroborated by permutational distance-based multivariate analyses (db-MANOVA, Table S2) for differences in plant community composition coupled with indicator species analysis^55^.

All forest sites were classified as production forests (i.e. > 1 m^3^ ha^-1^ year^-1^ biomass production) and managed with conventional Swedish forestry practices but differed in stand age and density depending on location. Although local differences in climate, parent material, and topography exist at this spatial scale, we assumed that grasslands, if and when abandoned and fully afforested, would share approximately the same vegetation and soil microbiome characteristics as their adjacent forest sites^20^. Climate data comprising mean annual temperature (MAT) and precipitation (MAP) for each site was collected from the Swedish Meteorological and Hydrological Institute’s (SMHI) long-term local weather stations. Information about the parent material for each site was gathered from the Swedish Geological Survey (SGU).

### Soil sampling

The sites were sampled between July and September 2020 following the standardized protocol established in^56^. Briefly, at each site, 15 samples were collected across a circular area of ∼ 200 m^2^ using a soil corer (diameter = 3 cm, depth = 10 cm), and pooled into a bulk sample. Surface layers constituting litter and fibric material was removed prior to sampling but otherwise no distinction was made between the organic and mineral layers. A subset of the pooled sample was air dried (<40 °C) within 24 hours of collection and stored in a zip-lock plastic bag with silica gel to minimize humidity and prevent development of molds during transit and stored frozen (−20 °C) until molecular analyses. All equipment was sterilized with 95% ethanol between sites.

### Abiotic drivers of soil diversity

Approximately 15 g of soil was used to analyze pH (1:5 soil:water suspension). Available phosphorus (P-AL) and potassium (K-AL) were extracted using ammonium lactate and acetic acid at pH 3.75^57^ and analyzed using the stannous chloride-molybdate procedure (P-AL) and inductively coupled plasma atomic emission spectroscopy (ICP-AES), respectively. Exchangeable calcium and magnesium concentrations were measured in ammonium acetate extract (pH = 7.0) using ICP-AES at Agrilab Uppsala, Sweden. Total C and N contents was determined on aliquots (1-20 mg) of air-dried soil using an Elemental Analyzer (Euroa EA, Eurovector, Milano, Italy). We used leaf dry matter content (LDMC) as a proxy for leaf quality, as this has been shown to correlate well with litter quality and decomposition in successional systems in our study area^14^ LDMC values were derived from the TRY database^58^ (Ver. 6.0) for 120 out of the 235 plant species (51.1 %), and converted to community-weighted means for each site. While we acknowledge that traits derived from plant databases disregard intraspecific variation that could affect the results, it has been shown that interspecific trait variation is of considerably greater importance than intraspecific variation^59^.

### Substrate-induced respiration

We performed catabolic profiling on a subset (*n* = 156, or *n* = 77 paired grassland-forest sites, Table S1) of air-dried soils using substrate-induced respiration (SIR) for a range of carbon substrates of differing complexity. Briefly, 8 mL solutions of glucose, glycine, oxalic acid, autolyzed yeast, lignin, and chitin was added to 4 g dry-weight equivalent of fresh soil (1 analytical replicate per solution). After 1 h of pre-incubation, soils were incubated for 4 h at 20 °C except for lignin and chitin which were incubated for 24 h. After incubation, respiration for each amendment was determined using Gas Chromotography (Trace CG Ultra Gas Chromatograph (Thermo Fisher Scientific, Milan, Italy)). Basal respiration rates, measured on 8 mL ddH_2_O samples, were subtracted from each substrate-specific respiration rate to account for the added effect of water flushing (Birch effect)^60^. We used Multiple substrate-induced respiration (MSIR) as a proxy for the communities’ carbon cycling capacity^61^, which was calculated by first subtracting basal respiration rates from each substrate-amended soil, and subsequently summing the individual substrate respiration rates for each sample.

## MOLECULAR ANALYSES

### DNA extraction and sequencing

DNA was extracted from 200 mg of dried and milled soil samples using the PowerMax Soil DNA Isolation Mini kit (Qiagen GmbH, Hilden, Germany) following the manufacturer’s instructions. The extracted DNA was quality-checked with 260/280 and 260/230 nm wavelength ratios using a NanoDrop™ (Thermo Scientific, Massachusetts, USA) and stored at −20 °C until sequencing. For bacterial amplicons, the universal prokaryote primers 515F and 926R were used to amplify the 16S V4 subregion of rRNA gene^62^. DNA samples were amplified using the following conditions in three replicate runs: 95°C for 15 min, followed by 26 cycles of 95°C for 30 s, 50°C for 30 s and 72°C for 1 min with a final extension step at 72°C for 10 min. The 25 μl PCR mix consisted of 18 μl sterilized H_2_O, 5 μl 5 × HOT FIREPol Blend MasterMix 0.5 μl of each primer (20 μl) and 1 μl template DNA (final concentration of 400 nM). For amplicons of eukayotes/fungal ITS, we used the universal eukaryote PCR primers ITS9mun and ITS4ngsuni^63^. PCR amplification followed the protocol described in^64^. Briefly, 0.5 µl of each forward and reverse primer (20 mM), 1 µl of DNA extract and 18 µl ddH_2_O were used in combination with 5 µl of 5L×LHOT FIREPol Blend Master Mix (Solis Biodyne, Tartu, Estonia). Thermal cycling followed an initial denaturation at 95 °C for 15 min; 25–30 cycles of denaturation for 30 s at 95 °C, annealing for 30 s at 57 °C, elongation for 1 min at 72 °C; final elongation at 72 °C for 10 min; and storage at 4 °C. Duplicate PCR products were pooled and quality checked on a 1 % agarose gel. All primers were tagged with a 10-base pair barcode for sample identification. Blanks containing ddH_2_O instead of DNA template were used as negative controls in the library preparation. The amplicons from the replicates were pooled, purified using a purification kit containing agarose gel (FavourPrep Gel/PCR Purification mini Kit-300 Preps; Favourgen) and shipped for library preparation in the sequencing service facility of University of Tartu (the Estonian Biocenter). Sequencing of ITS was performed on the PacBio Sequel System at Novogene (UK), and 16S libraries were sequenced on two runs using an Illumina MiSeq platform (2 × 250 bp paired-end chemistry).

Shotgun metagenomic sequencing was performed on pooled equimolar amounts of DNA from a subset (*n* = 94, or *n* = 47 pairs) of grassland and forest sites (Table S1). Library preparation and sequencing were performed at the service provider facilities (Novogene Europe) using the Novogene NGS DNA Library Prep Set kit and sequenced on Illumina NovaSeq with 2L×L150Lbp paired end reads.

## BIOINFORMATICS

### Metabarcoding

We used the LotuS2 version 2.22^65^ pipeline to quality-filter, demultiplex, and process the filtered reads into OTUs. Chimera detection and removal was done using Uchime^66^ with all singletons and sequences shorter than 100 bp discarded. Clustering of sequences was done using a *de-novo* clustering algorithm in UPARSE^67^ based on a 97% similarity threshold. Taxonomy was assigned against the SILVA (ver. 138.1) and UNITE (ver. 8.1) databases for prokaryotic and fungal sequences respectively. All datasets were manually curated to remove contaminant sequences based on negative controls. OTUs representing archaea, chloroplasts, eukaryotes, and mitochondria were omitted from the bacterial dataset, and OTUs unassigned at class level were omitted from the ITS dataset after OTU clustering. To reduce biases due to low sequence coverage, we discarded samples with < 100 reads (*n* = 8 samples) and < 3000 reads (*n* = 3 samples) from the fungal and bacterial dataset, respectively. This resulted in a total of 204 samples comprising 8 532 758 reads (sample mean ± SD = 41 8127 ± 33 514) across 8937 OTUs for bacteria, and 199 samples with a total of 437 006 reads (sample mean ± SD = 2196 ± 2750) covering 2818 OTUs for fungi.

### Shotgun metagenomics

Analysis of metagenomic reads was done using MATAFILER pipeline^68^ using a workflow optimized for complex environmental metagenomes^69^. Briefly, reads obtained from the shotgun metagenomic sequencing of soil samples were quality-filtered by removing reads shorter than 70% of the maximum expected read length (150 bp), with an observed accumulated error > 2 or an estimated accumulated error >2.5 with a probability of ≥ 0.01, > 1 ambiguous position or if base quality dropped below 20, using sdm (version 1.46)^65^. All 95 samples produced sufficient quantity of reads (average 37 186 296 ± 7 365 270 reads per sample) and were retained for statistical analyses. To estimate the functional composition of each sample, similarity search using DIAMOND (version 2.0.5; options −k 5 −e 1e-4 –sensitive) in blastx mode^70^ was employed. Prior to that, the quality-filtered read pairs were merged using FLASH (version 1.2.10)^71^. The mapping scores of two unmerged query reads that mapped to the same target were combined to avoid double counting. In these cases, the hit scores were combined by averaging the percent identity of both hits. The best hit for a given query was based on the highest bit score and highest percent identity to the subject sequence. Using this method, we calculated the relative abundance of (clusters of) orthologous gene groups (OG) by mapping quality-filtered reads against the eggnog database (version 5)^72^. We used the MicrobiomeCensus pipeline to estimate average genome size (AGS) of bacterial communities^73^.

### Inferring pathways related to C-N-P cycling

Quality-filtered reads were blasted against the *CAZyme* database^74^ to derive OGs related to C-cycling. These were further mapped against *CAZymes* with known substrate affinities based on a custom database provided in ref^75^. For N- and P-cycling genes, we annotated genes from the Kyoto Encyclopedia of Genes and Genomes (KEGG) Orthology database^76^ and mapped these against custom databases containing N- and P-cycling genes provided in^77,78^. The total number of metagenomic reads, including reads and OG for each nutrient cycling process are shown in Table S4.

### Predicted metagenomes

Testing for trade-offs between functional redundancy and specialization require joint taxa-function matrices to assess functional distances between taxa^31^. We used *Picrust2* (ver. 2.5.2)^32^ to predict functional metagenomes (PFM) from quality filtered 16S rRNA sequences. The weighted nearest sequenced taxon index (NSTI) of the PFM ranged between 0.129 and 0.312, indicating generally good predictions. From the PFMs, we next extracted the functional pathways related to C-N-P cycling based on OG from the KEGG database for subsequent analyses.

## STATISTICAL ANALYSES

All statistical analyses were performed using R (ver. 4.4.1) using the following packages *effectsize*^79^*, vegan*^80^*, adiv*^81^*, lme4*^82^*, microniche*^83^*, rfpermute*^84^*, SRS*^85^. For all analyses involving multiple comparisons, *p*-values were adjusted with Benjamin-Hochberg corrections.

### Diversity analyses

We used Shannon’s *H’* to estimate taxonomic (i.e. OTU) and functional (C-N-P-cycling genes) alpha diversity as it considers both richness and evenness. We used linear mixed models (LMM) with sample pairs, the spatial distance between them, and parent material as random factors to account for the matched structure of the dataset and any differences in geological substrate. As forests are the putative successional endpoint of the abandoned grasslands, we set these as the reference level in the LMMs, with subsequent pairwise comparisons in a separate model without set reference level to examine differences between grassland stages. Differences in community composition (beta diversity) were tested using perMANOVA (10^^4^ permutations) on Bray-Curtis distances for taxonomic and functional matrices. As grasslands across differing stages of succession were overall similar in terms of alpha and beta diversity, we pooled these into a single grasslands category and used effect sizes (partial ω) to estimate the magnitude of the diversity effects (alpha, beta) between grasslands and forests. Prior to all diversity analyses, OTU matrices were normalized using scaling with ranked subsampling (SRS)^85^, with a minimum sequencing depth (*cmin* = 200 and 3000) for fungi and bacteria, respectively. Functional matrices were normalized by rarefying to minimum sequencing depth.

### Assessing functional redundancy and specialization, and their links to carbon-cycling capacity

We used a set of approaches to examine the genetic redundancy related to nutrient cycling during succession. Firstly, taxonomic diversity (Shannon’s *H’* of OTU tables) was used as a predictor of functional diversity (Shannon’s *H’* of C-N-P-genes from metagenomes) in ordinary least-square regression (OLS) models. For these analyses, all grassland stages were pooled into a single grassland category to match the sample size of forest sites.

Significant fits between taxonomic and functional diversity was deemed indicative of low functional redundancy^27^.

We next used Levin’s niche overlap^86^ (implemented in the *Microniche* package^83^ to quantify the metabolic overlap of C-N-P genes across land uses, using rarefied metagenome matrices. Only genes above the default limit of quantification (LOQ = 1.65) were kept for subsequent analyses. We assumed that genes with high overlaps are occur widely within a given land use, akin to generalist taxa in classical niche theory, whereas low overlaps indicate specialization^86^. The analyses were repeated using specific nutrient cycling pathways. We used OLS and second-order polynomial regressions to assess changes in average genetic overlap across successional stages, with pairwise Wilcoxon tests were used to test for differences in average overlap of substrates (C-cycling) and nutrient cycling pathways (N-P-cycling) between land use-stages. Differences in SIR between paired grassland and forest sites were tested through effect size measurements (Hedge’s *d*) for each assessed substrate. We then examined the relationship between the communities’ total carbon cycling capacity and diversity by regressing MSIR as response variable against taxonomic and functional diversity as predictor variables in OLS models.

### Assessing trade-offs between functional redundancy and diversity

Trade-offs between community-level functional redundancy and diversity, as well as drivers of both measures, were examined with predicted bacterial metagenomes based on the framework developed in refs.^30,31^. We considered KO pathways related to C-N-P-cycling as traits and related these to the relative abundances of bacterial OTUs at each site. Specifically, we first scaled all pathways (range 0-1) by their minimum and maximum values and calculated the functional Euclidean distances between all pairs of OTUs. This step was done for each land-use stage separately to avoid the inclusion of OTUs not found within the meta-community. Functional distances were then scaled by division with their land-use specific maximum values. From this, functional diversity (Rao’s quadratic diversity *Q*), functional redundancy *R*, and the Simpson dominance index *D* were calculated using the *adiv* package^81^. The position of each site along the axes of these three measures were assessed and visualized using ternary plots and perMANOVA tests (Bray-Curtis distances, 9999 permutations). Note that the limited availability of fungal genomes precludes similar analyses for the fungal dataset.

### Data availability

The raw amplicon sequences have been deposited at NCBI under accession PRJNA994701. Shotgun metagenome sequence data are deposited at the Sequence Read Archive (SRA) under the project ID PRJEB56463.

## ACKNOWLEDGEMENTS

We wish to thank Anders Glimskär Remiil and Bertil Westerlund Riksskogstaxeringen for contribution with finding sites and providing metadata.

Grants:

**TRS:** C.F. Lundströms Stiftelse, CF2023-0019, Lars Hiertas Minne FO2021-0302

**MB:** Swedish University of Agricultural Sciences (early career grant), the Swedish Research Councils Formas (Grant 2020–00807) and the Swedish Research Council (VR; Grant 2021–03724).

**FH** & **JF**: were supported by the UKRI Biotechnology and Biological Sciences Research Council (BBSRC) Institute Strategic Programme Food Microbiome and Health BB/X011054/1 and its constituent project BBS/E/F/000PR13631, Decoding Biodiversity BBX011089/1 and its constituent work packages BBS/E/ER/230002A and BBS/E/ER/230002B, FH as well as by European Research Council H2020 StG (erc-stg-948219, EPYC).

**JF** was supported by the BBSRC Norwich Research Park Biosciences Doctoral Training Partnership, BB/T008717/1.

---

**Extended Data Fig. 1:**
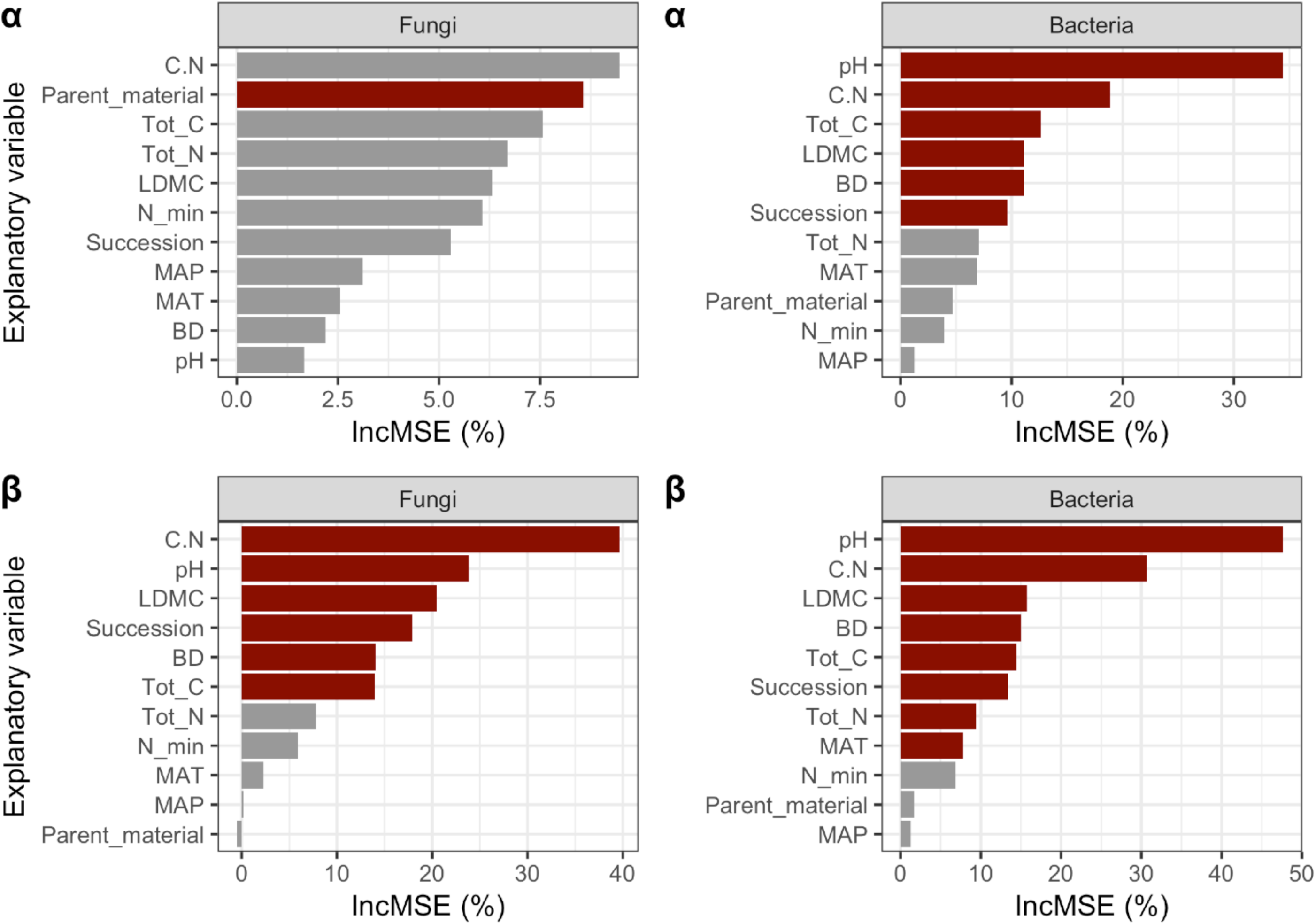
Drivers of microbial diversity. Barplot showing results from random-forest based variable selection for drivers of fungal and bacterial alpha- and beta diversity. IncMSE (%) describes the relative importance of the explanatory variable in the predicted response variable. Red color denotes that the explanatory variable is significantly (*p* < 0.05) linked to the response variable.

**Extended Data Fig. 2:**
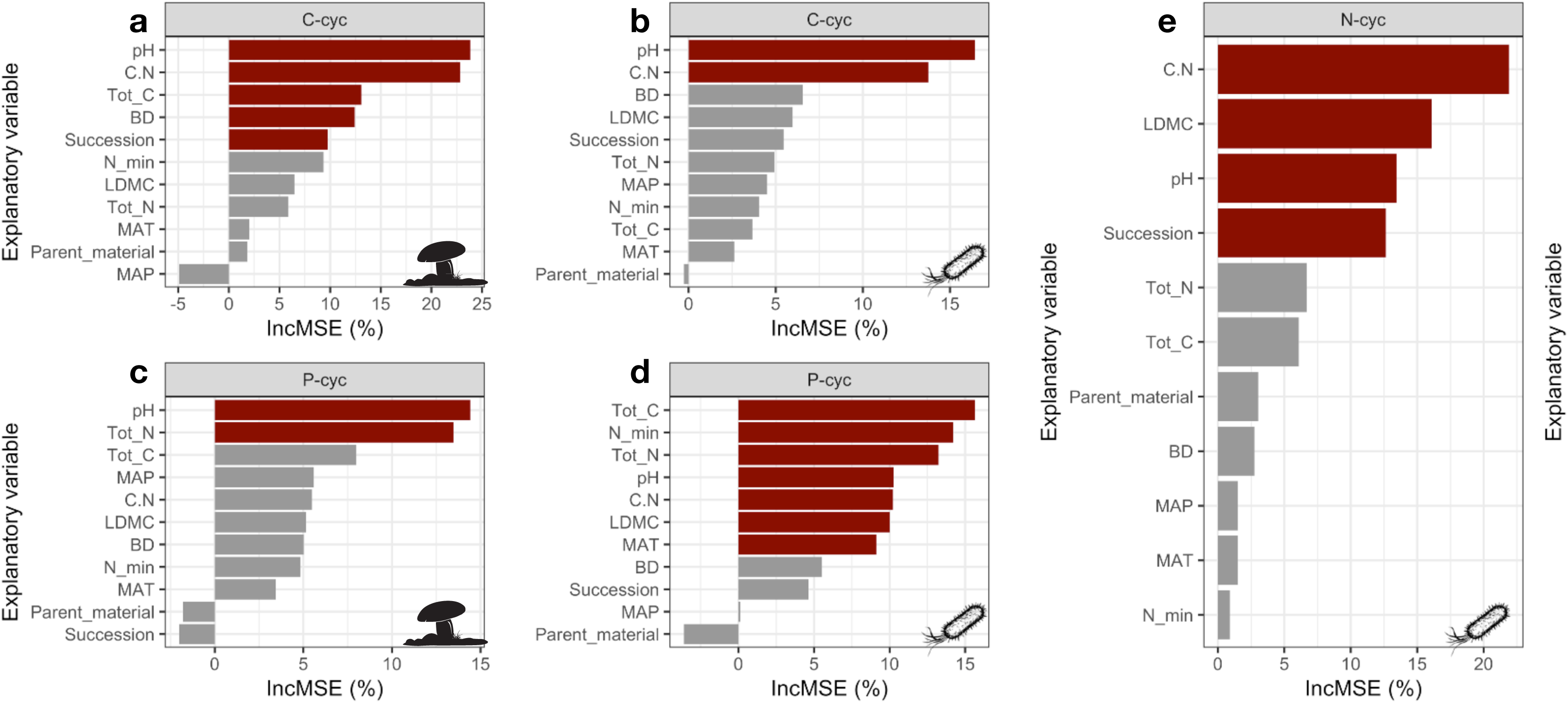
Drivers of functional diversity. Barplots showing results from random-forest based variable selection for drivers of fungal and bacterial functional (i.e. genetic) diversity across differing nutrient cycling processes. IncMSE (%) describes the relative importance of the explanatory variable in the predicted response variable. Red color denotes that the explanatory variable is significantly (*p* < 0.05) linked to the response variable.

**Extended Data Fig. 3:**
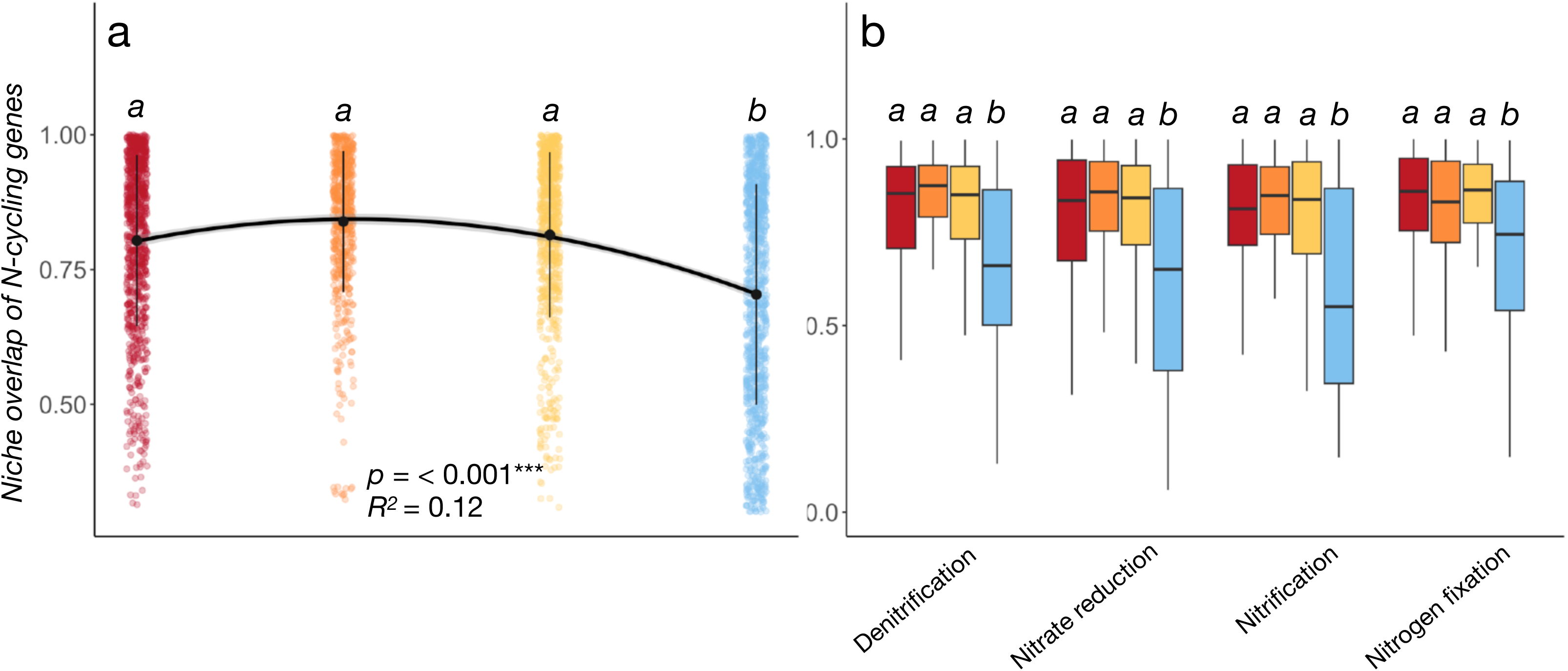
Genetic overlap of N-cycling genes and pathways. Genetic overlap of **a**) bacterial N-cycling genes across the land abandonment gradient. Black points show mean genetic overlap ± s.d. (vertical lines) for each land-use stage. Horizontal lines connecting land-use stages indicate significant (*p* < 0.05) ordinary least-square or second-order polynomial regression fits (Table S12). In panel **b**) the overlaps are partitioned across key N-cycling pathways. Letters indicate significant differences (*p* < 0.05) between land use-stages based on pairwise Wilcoxon Rank-Sum Tests, with full test result details founc in Table S13 & Table S16. DN = Denitrification; NR = Nitrate reduction; Nitr = Nitrification; NF = Nitrogen fixation.

**Extended Data Fig. 4:**
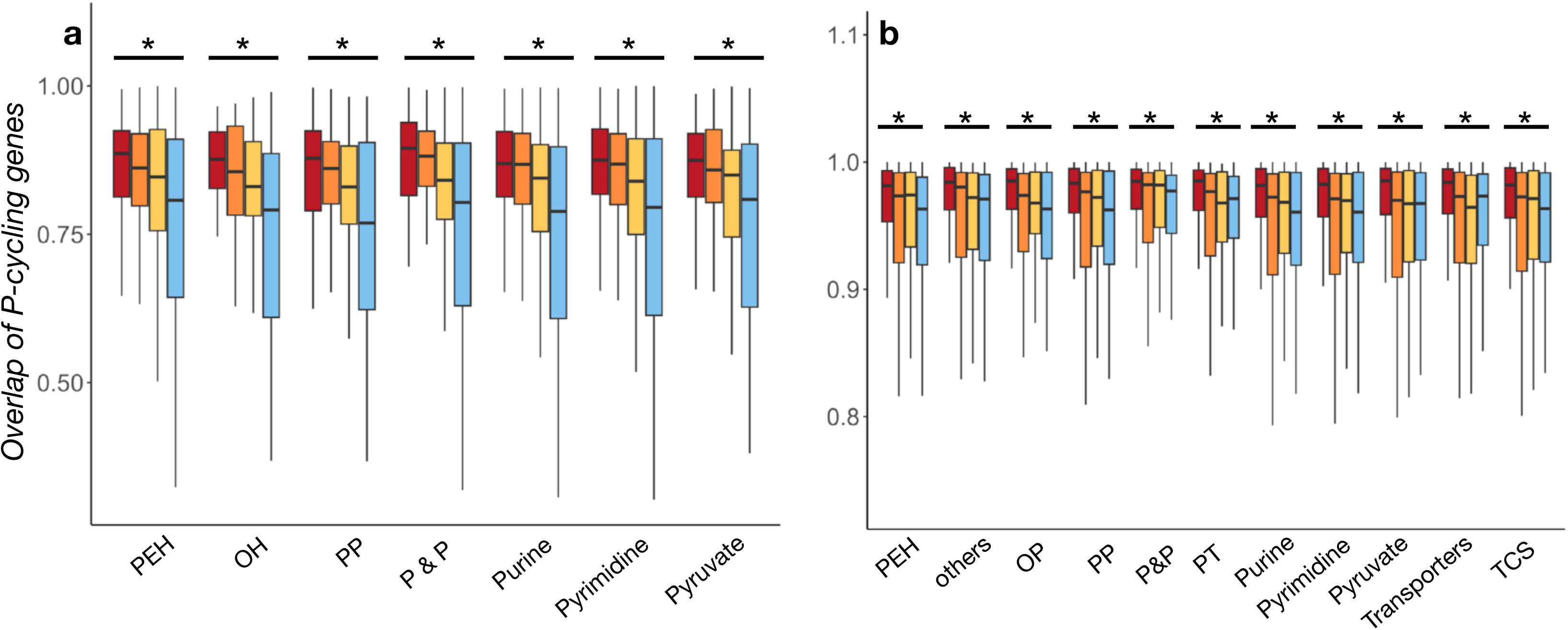
Genetic overlap of P-cycling pathways. Genetic overlap of **a**) fungal and **b**) bacterial P-cycling genes partitioned across pathways. Stars (*) above these pathways indicate significant (*p* < 0.05) differences between one or more land-use stages based on pairwise Wilcoxon tests, with full test result details found in Table S15. PEH = Phosphoester hydrolysis; OH = Organic phosphoester hydrolysis; PP = Pentose phosphate; P&P = Phosphonate and Phospinate metabolism; OP = Oxidative phosphorylation; PT = Phosphotransferase; TCS = Two component system.

**Extended Data Fig. 5:**
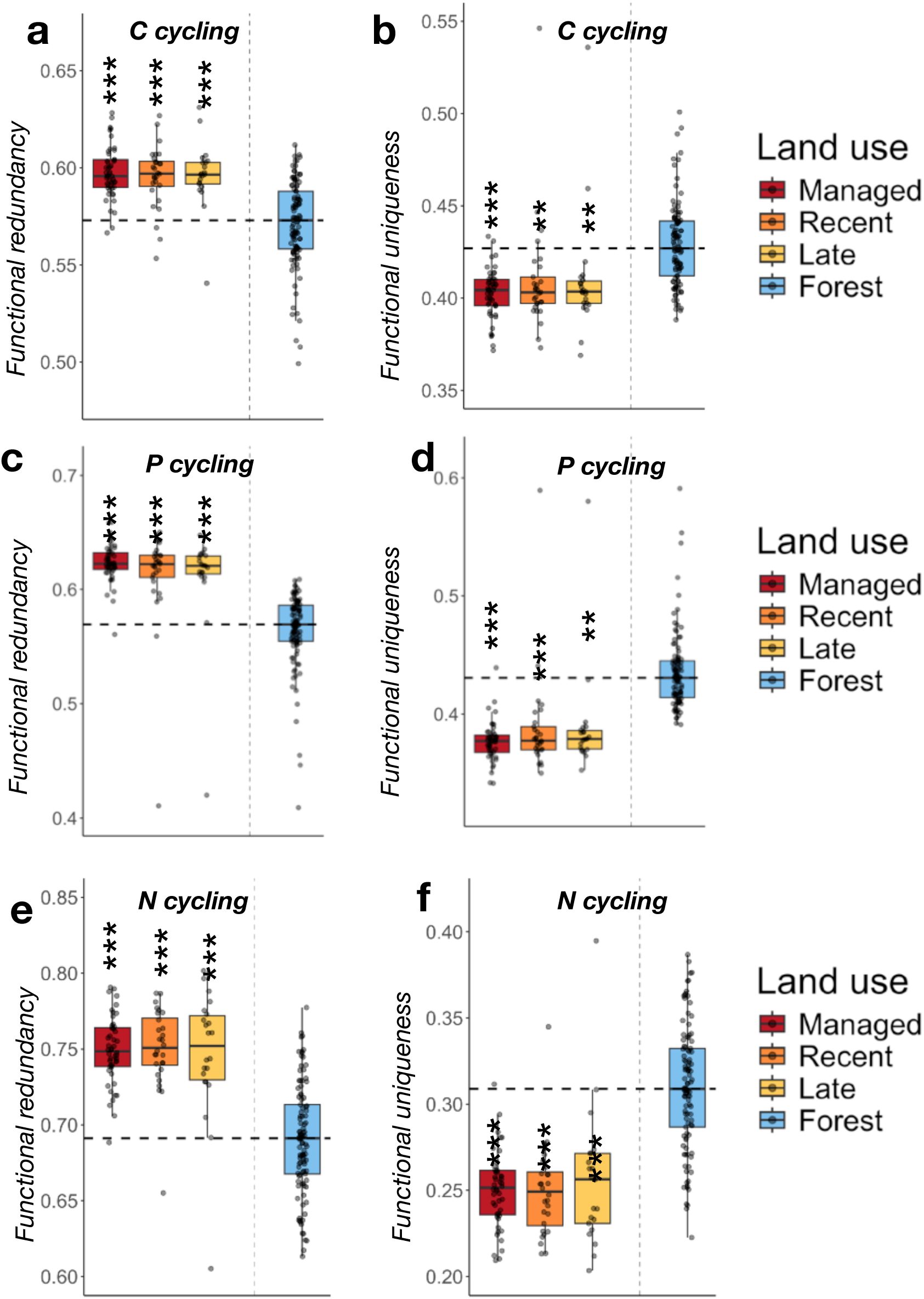
Functional redundancy and specialization. Boxplots showing levels of functional redundancy and functional uniqueness (specialization) across **a**) C-cycling, **b**) P-cycling, and **c**) N-cycling genes for bacterial communities based on predicted metagenomes. The lower and upper hinges of the boxplots represent the 25^th^ and 75^th^ percentiles, respectively, and the middle line is the median. The whiskers extend from the median by 1.5x the interquartile range. The dashed horizontal line indicates the median of the forest sites, with asterisks (*) denoting significant differences between paired grassland and forest sites based on mixed-effect linear models accounting for paired structure and spatial distance with the following significance levels: \**p* < 0.05, \*\**p* < 0.01, \*\*\**p* < 0.001.

**Extended Data Fig. 6:**
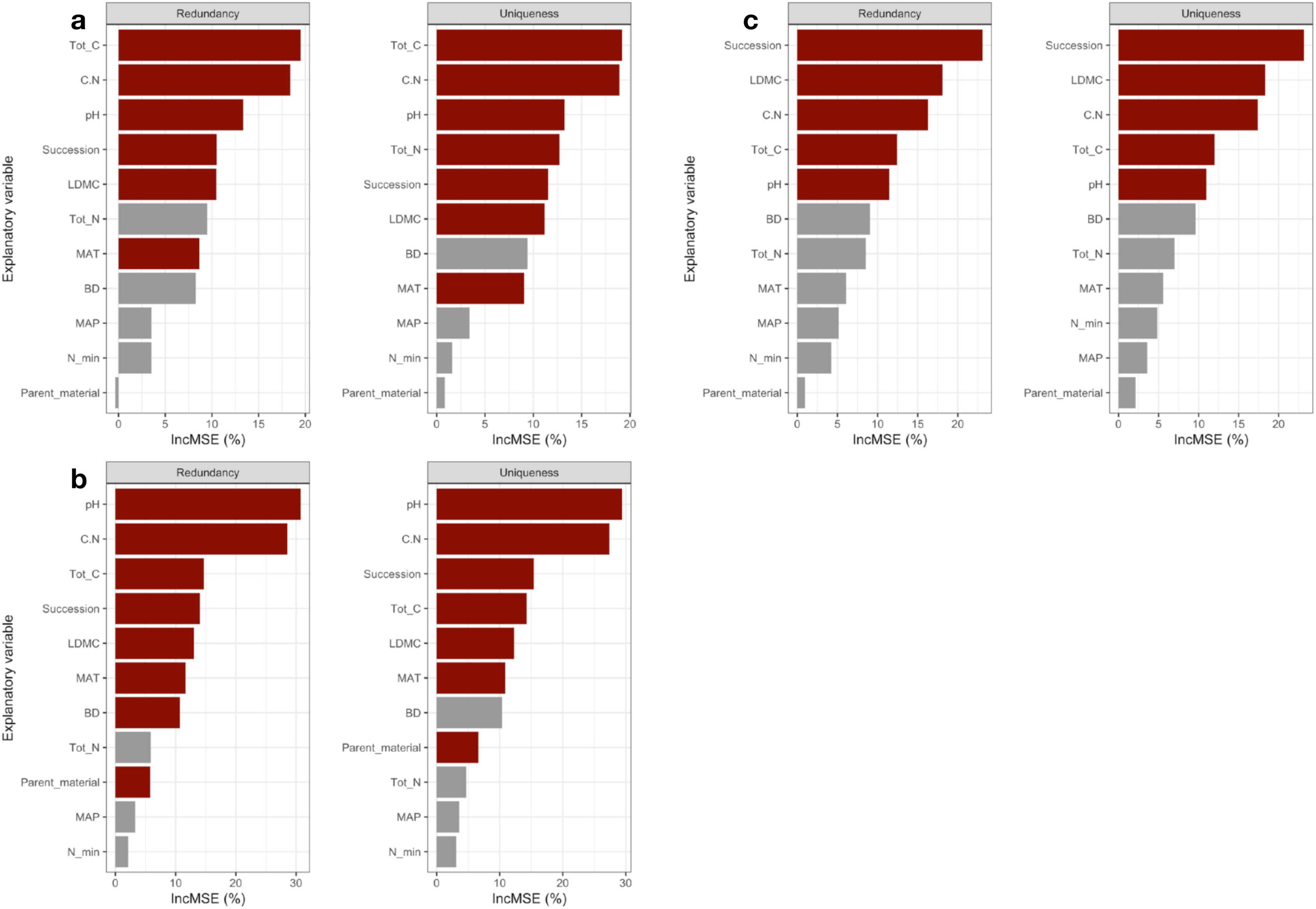
Drivers of functional redundancy and specialization. Barplots showing results from random-forest based variable selection for drivers of functional redundancy and functional uniqueness (i.e. specialization) across **a**) C-cycling, **b**) P-cycling, and **c**) N-cycling genes for bacterial communities based on predicted metagenomes. IncMSE (%) describes the relative importance of the explanatory variable in the predicted response variable. Red color denotes that the explanatory variable is significantly (*p* < 0.05) linked to the response variable.

**Extended Data Fig. 7:**
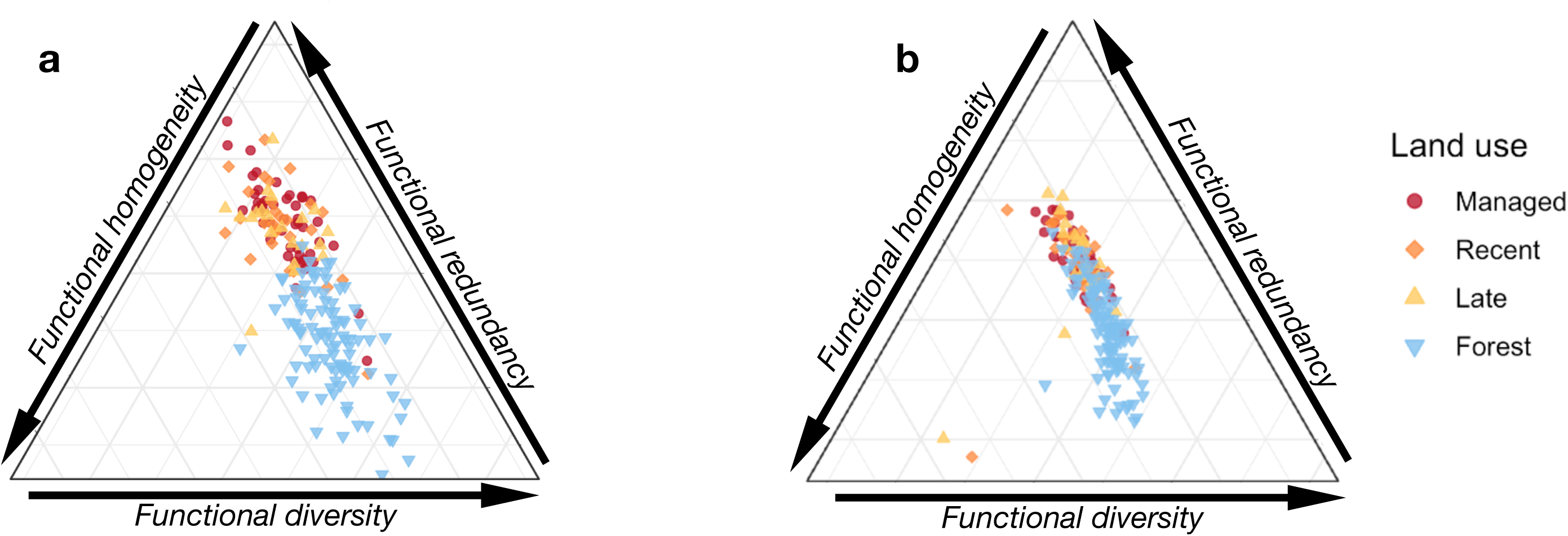
Ternary plots of N- and P-cycling across the redundancy-specialization axis. Ternary plots showing the clustering of sites according to their placement along the successional gradient for **a**) C-cycling, **b**) P-cycling, and **c**) N-cycling genes.

**Table S1:**
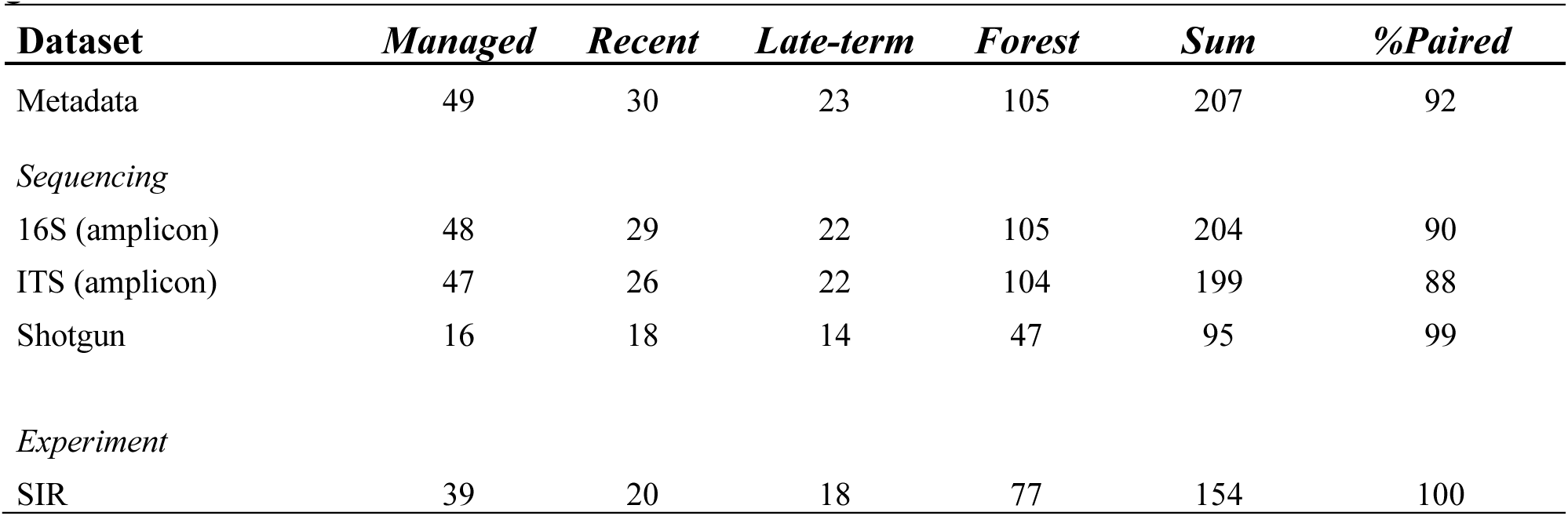
Site distribution for each dataset and land-use category across the successional gradient.

**Table S2:**
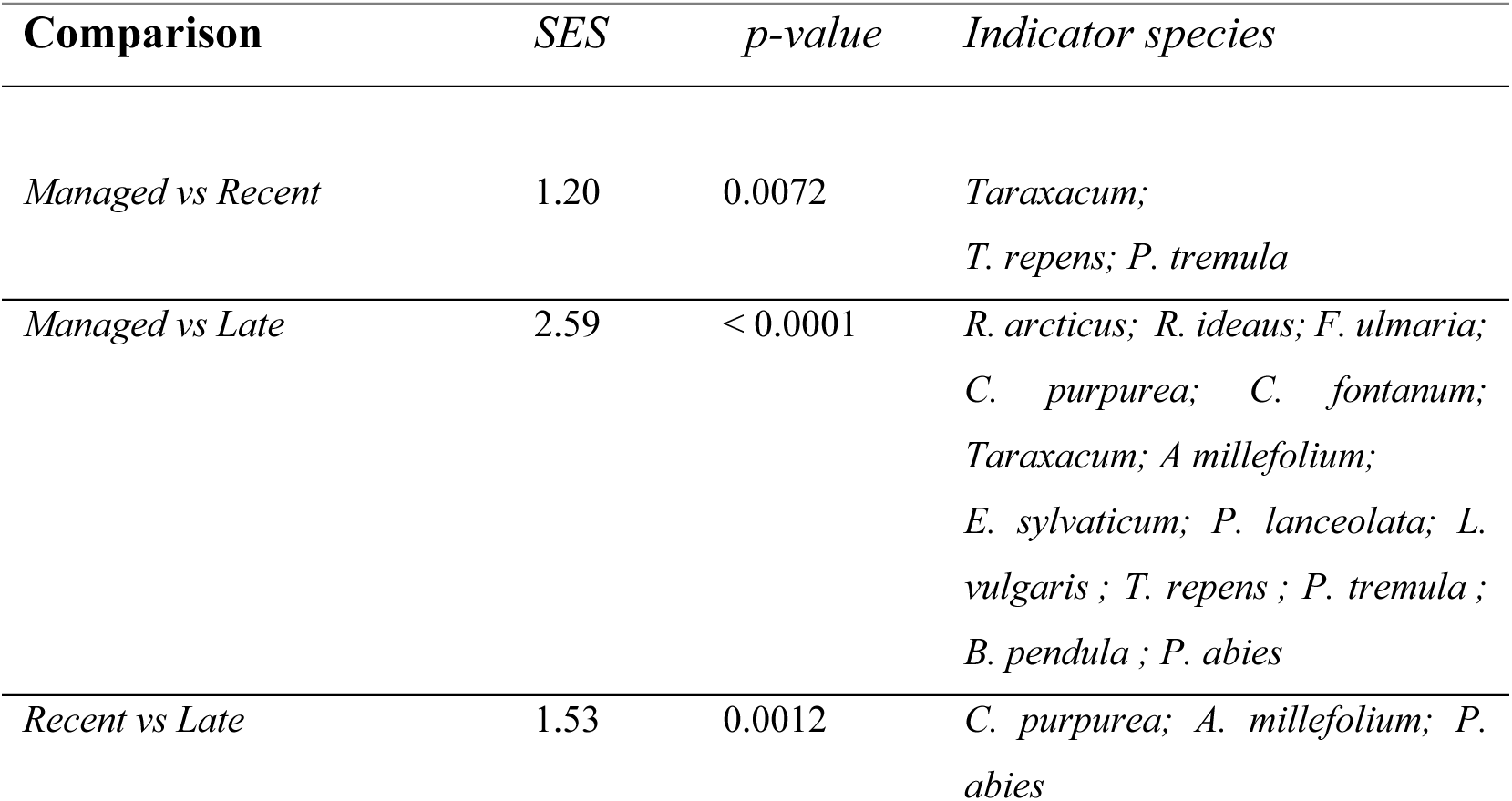
Combined tests for differences in composition and indicator species contributing to observed differences between pairs of plant communities. All indicator species significant (*p* < 0.05), and all *p*-values adjusted for multiple comparisons (Benjamin-Hochberg)

**Table S3:**
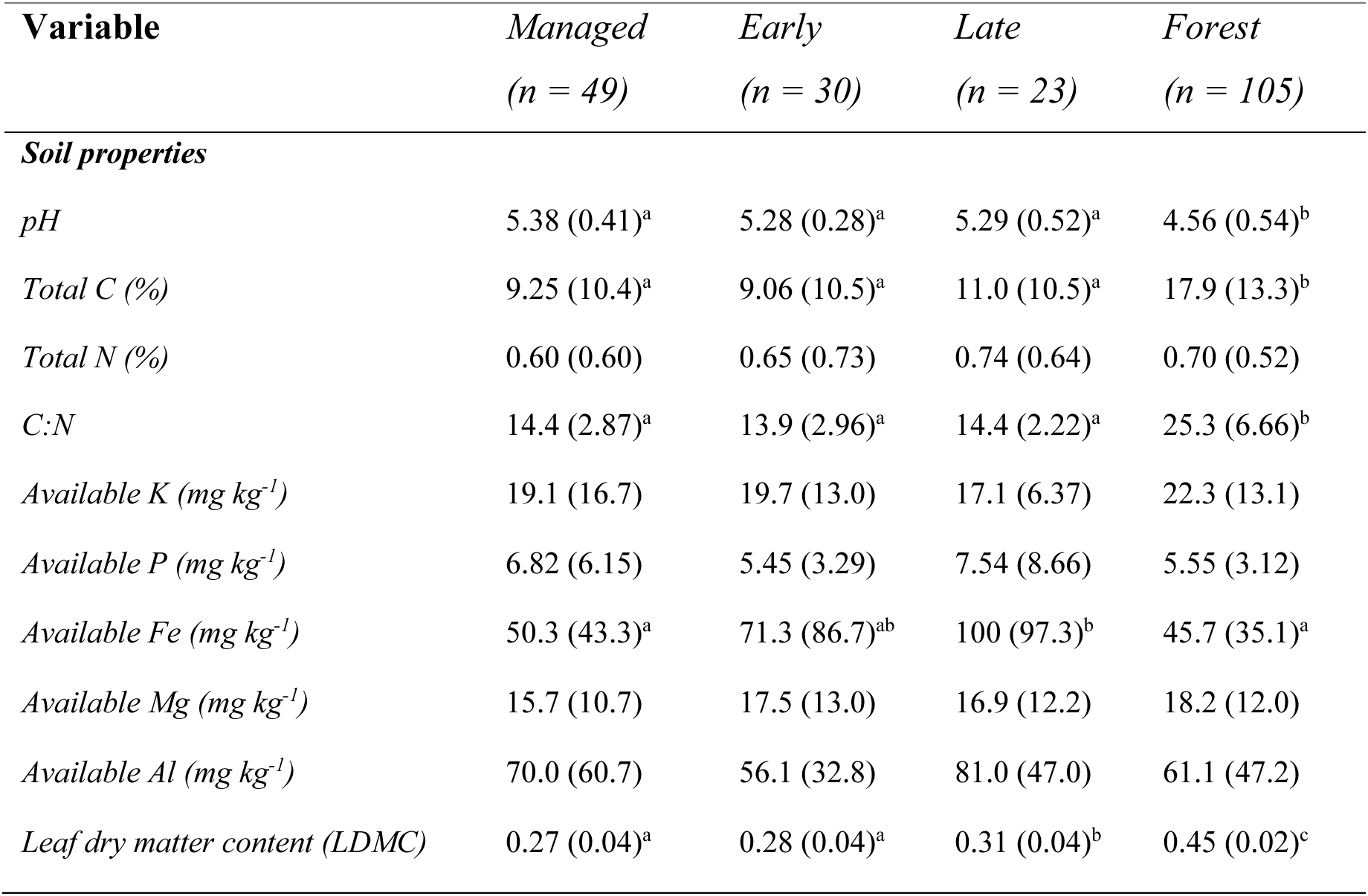
Soil and leaf properties (mean ± SD) across the successional gradient. Letters indicate significant differences (*p* < 0.05) based on pairwise Wilcoxon tests.

**Table S4:**
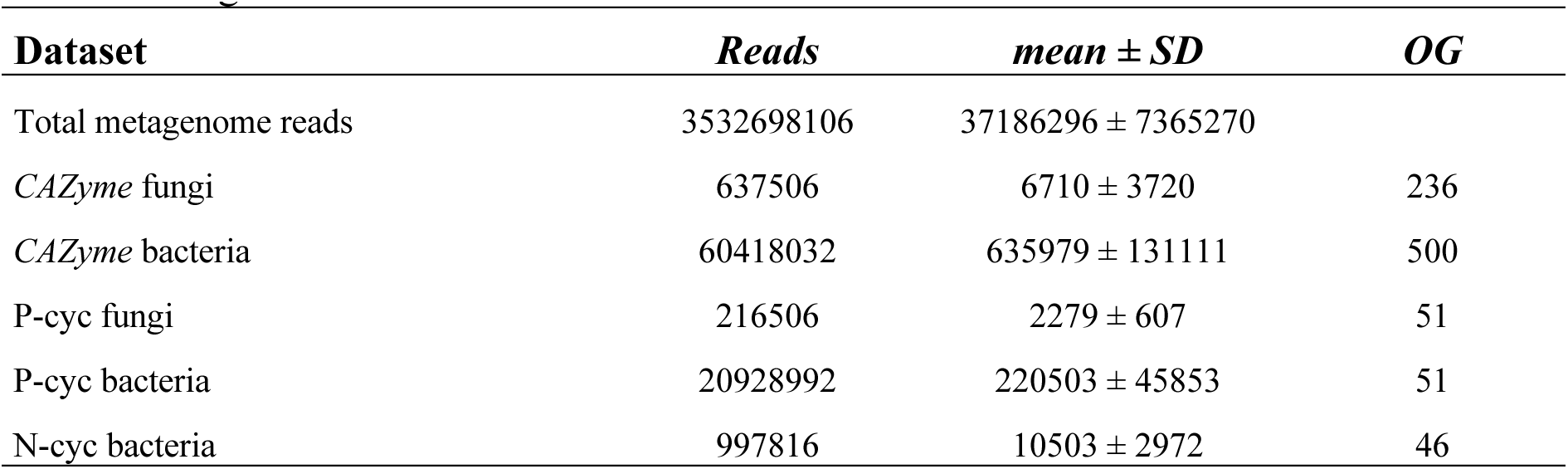
Number of reads and annotated orthologous groups (OG) related to nutrient cycling pathways obtained from the shotgun metagenomic sequencing of *n* = 95 samples across the successional gradient.

**Table S5:**
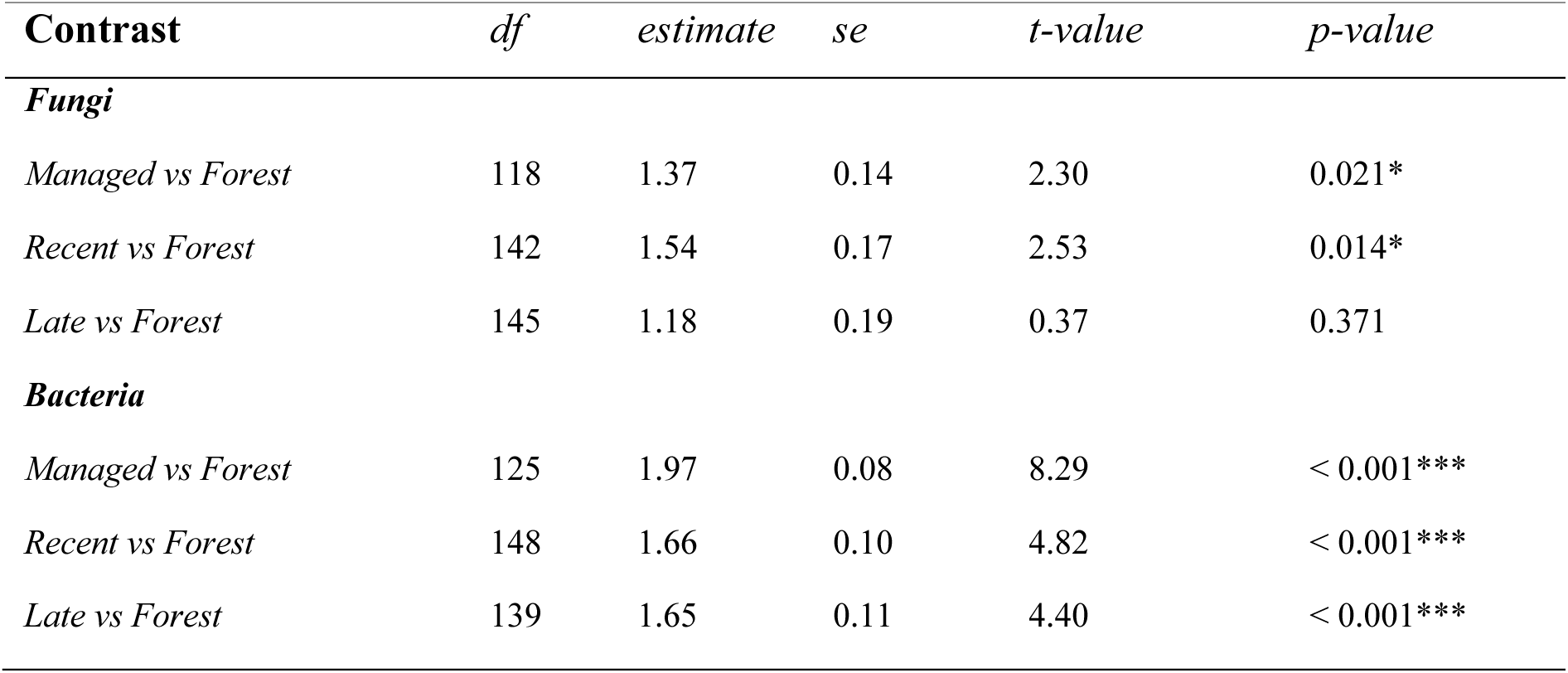
Differences in microbial taxonomic diversity (Shannon’s *H’*) between grassland and forest sites based on linear mixed models incorporating site pairs, spatial distance, and geological parent material as random factors.

**Table S6:**
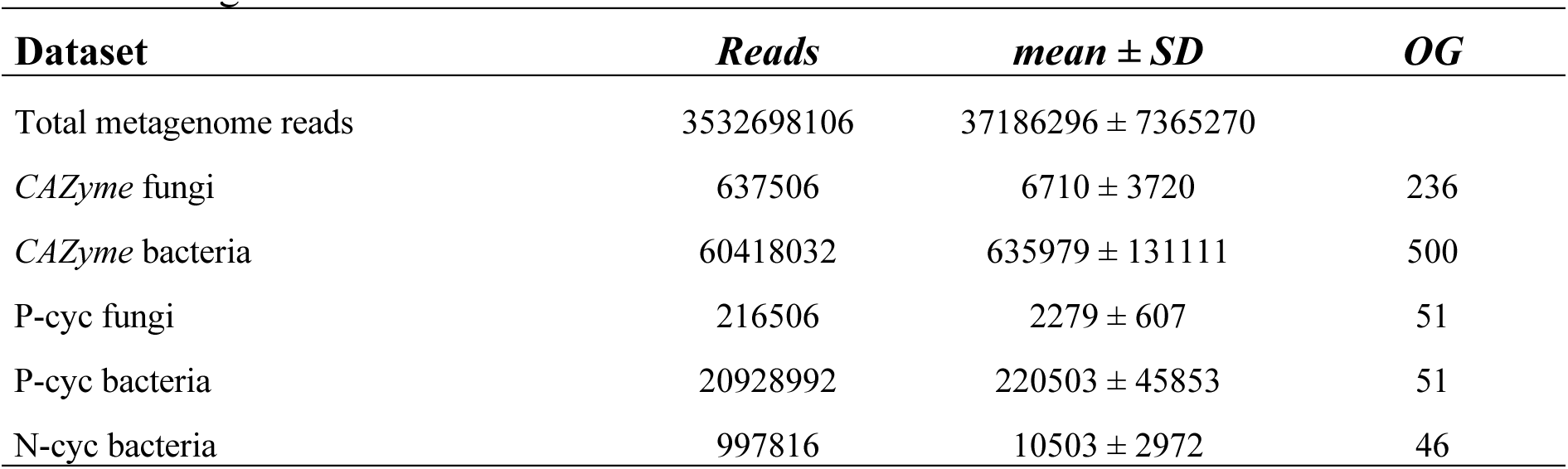
Differences in microbial taxonomic diversity (Shannon’s *H’*) between grasslands in differing successional stages based on linear mixed models incorporating geological parent material as random factor.

**Table S7:**
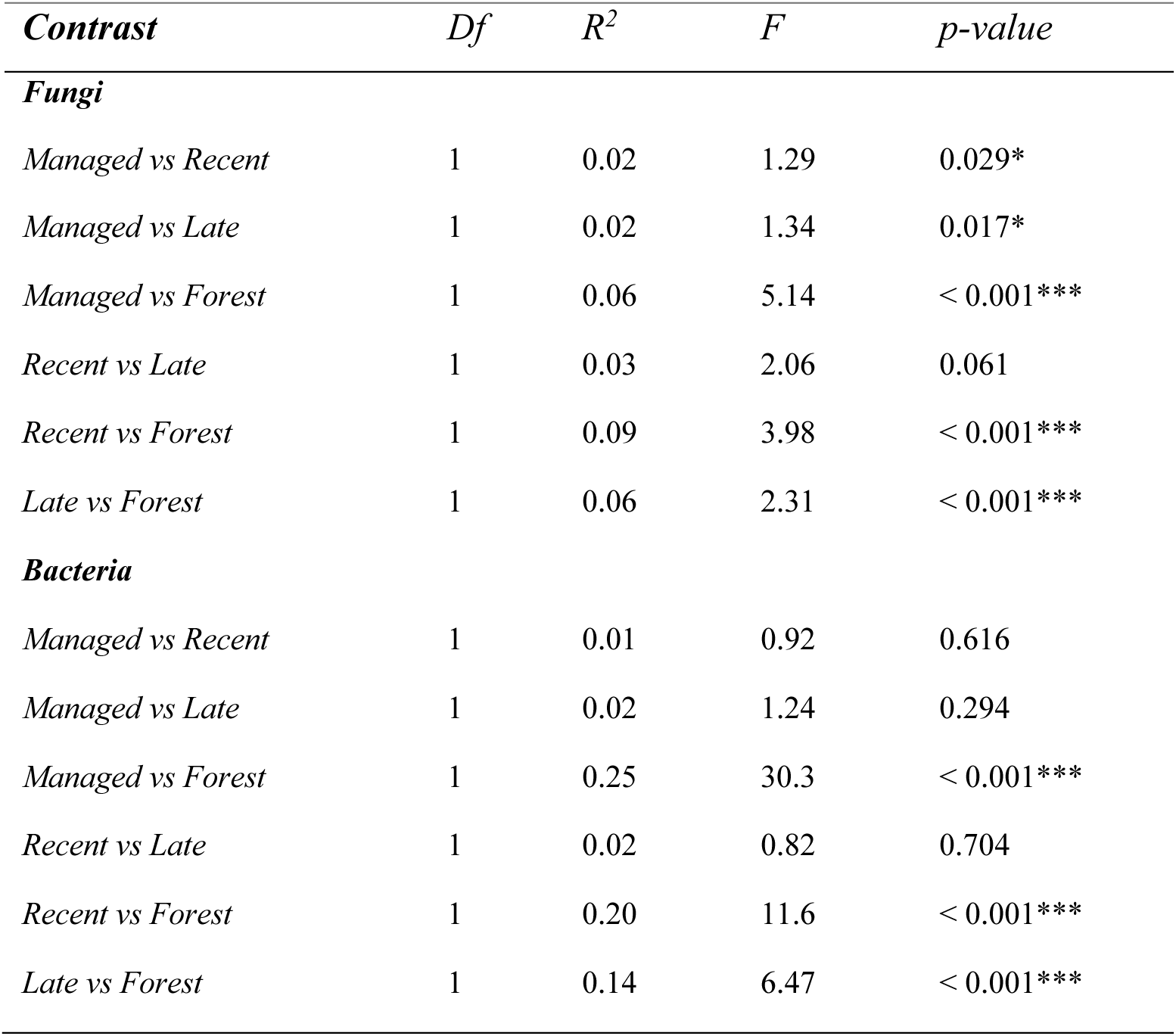
Results from permutational multivariate tests (perMANOVA) for differences in community composition between fungal and bacterial OTUs.

**Table S8:**
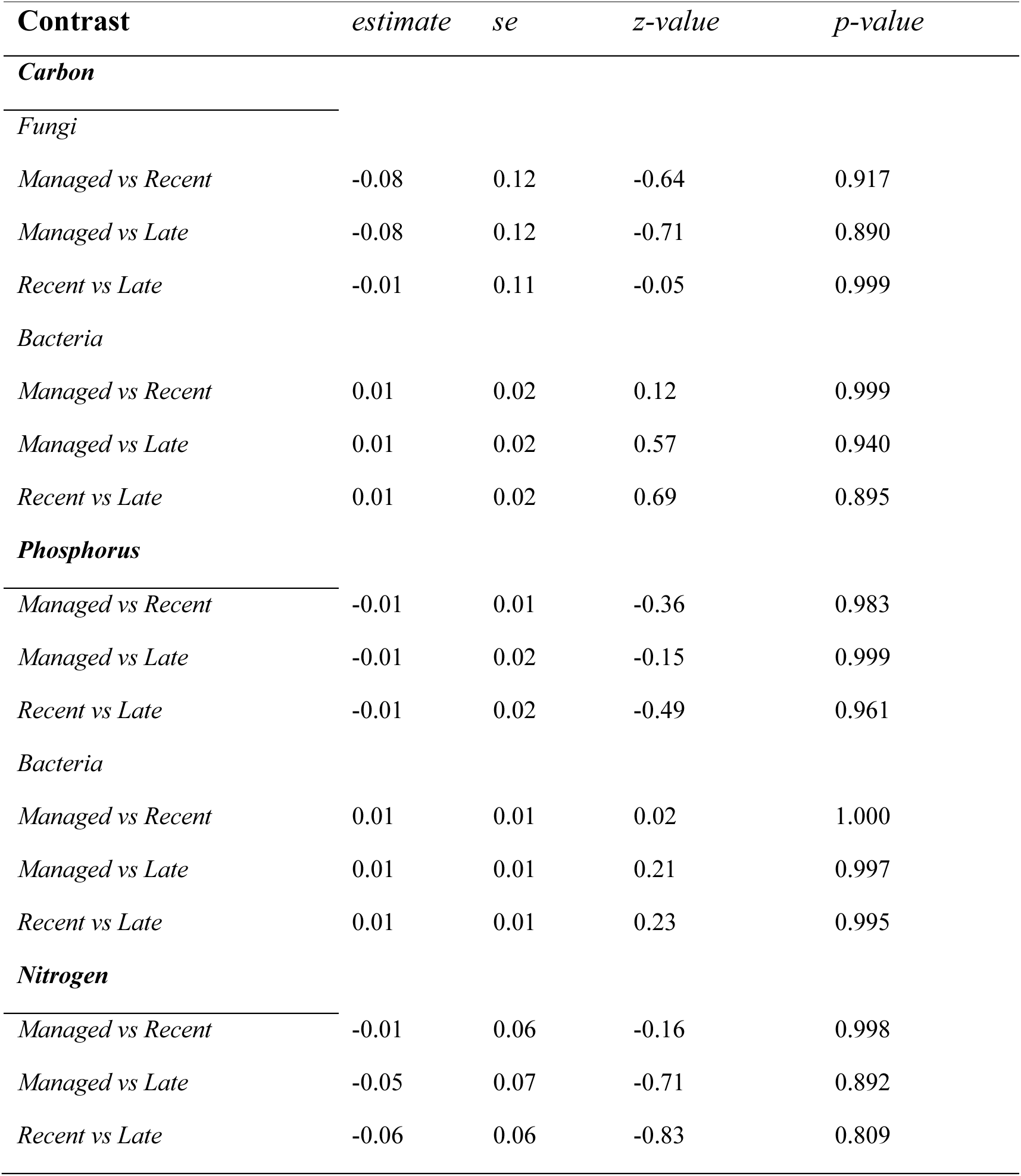
Differences in microbial biogeochemical genetic diversity (Shannon’s *H’*) between grasslands in differing successional stages based on linear mixed models incorporating geological parent material as random factor.

**Table S9:**
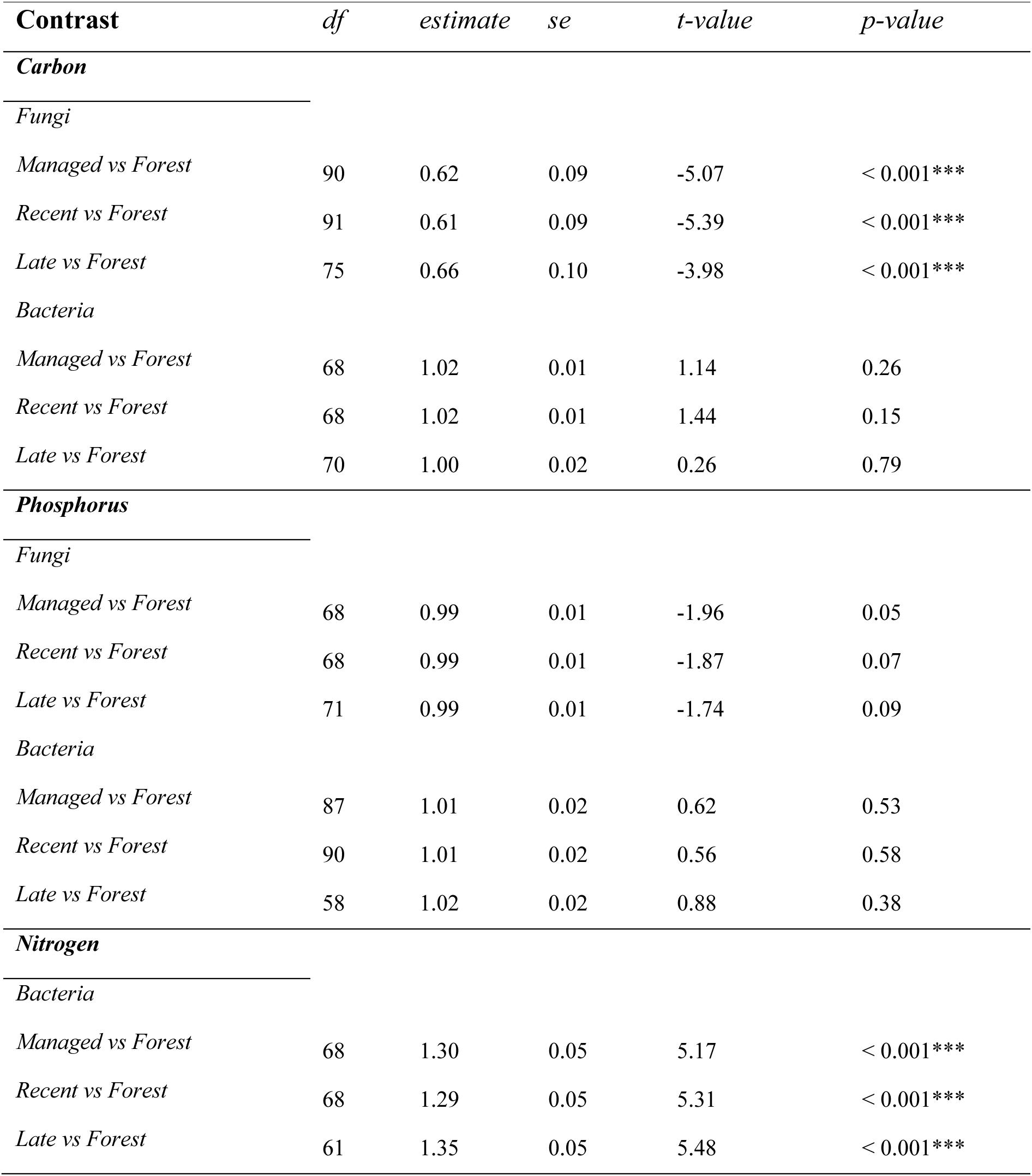
Differences in microbial functional diversity (Shannon’s *H’*) between grassland and forest sites based on linear mixed models incorporating site pairs, spatial distance, and geological parent material as random factors.

**Table S10:**
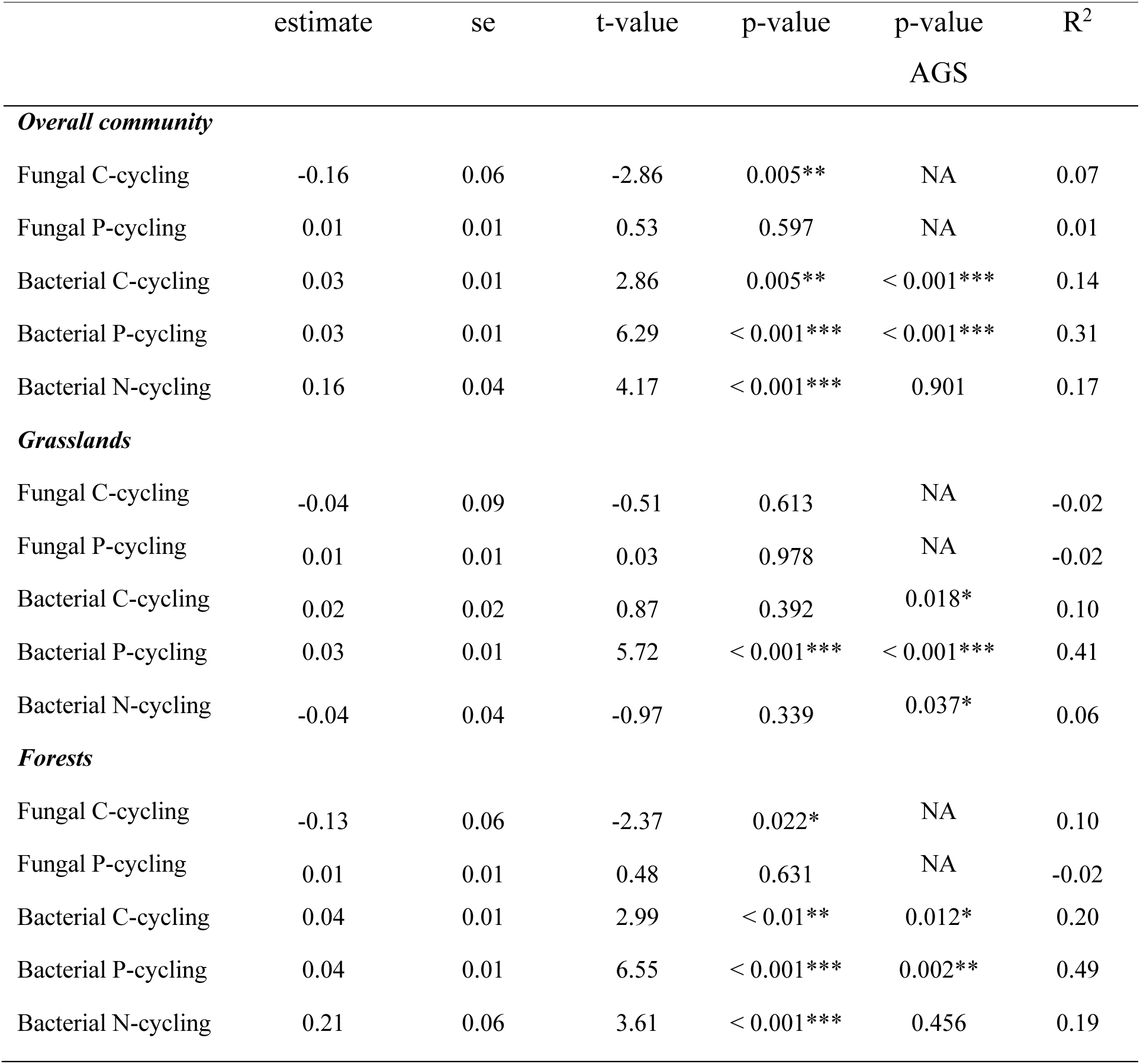
Relationship between taxonomic (Shannon’s *H’* of OTU matrices) and functional (Shannon’s *H’* of functional matrices) diversity for microbial communities between grassland successional stages, including when divided by ecosystem type (grassland, forest). Results based on ordinary least-square (OLS) regression. For bacterial communities, the average genome size (AGS) was included as a covariate in the regressions.

**Table S11:**
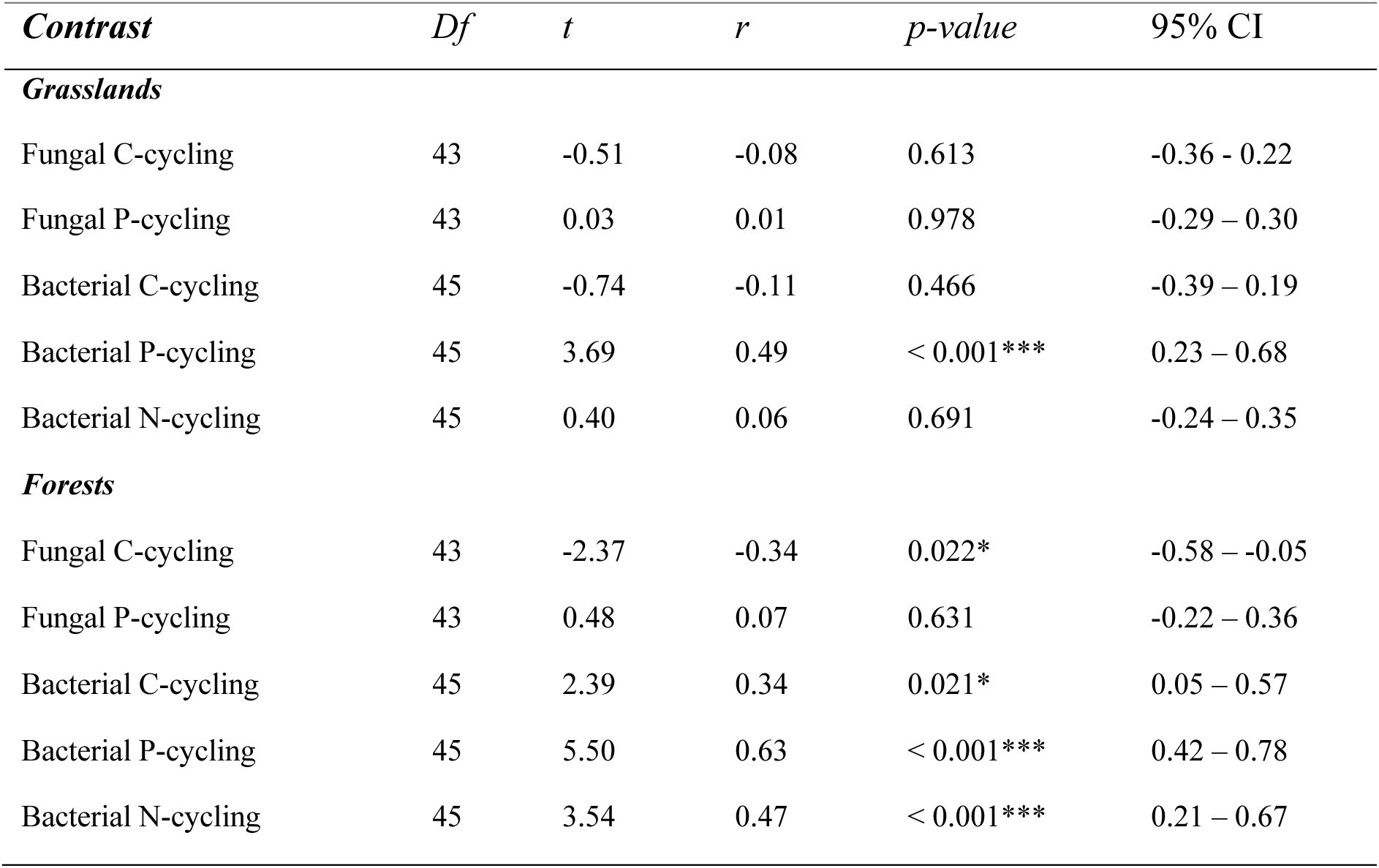
Pearson correlation tests between taxonomic (Shannon’s *H’* of OUT matrices) and functional (Shannon’s *H’* of functional matrices) diversity for microbial communities divided by ecosystem type (grassland, forest).

**Table S12:**
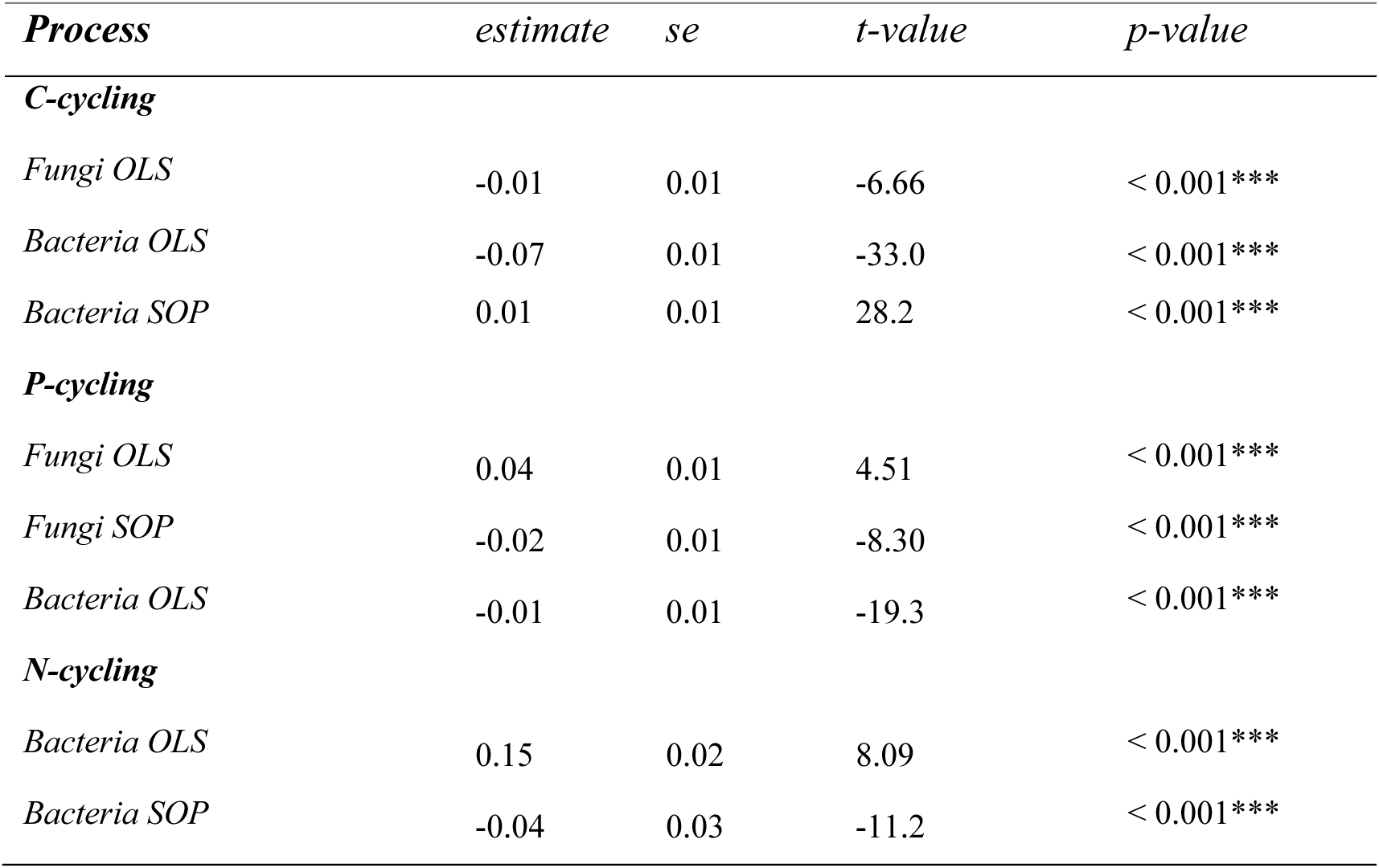
Results from ordinary least-square regression (OLS) and second-order polynomial (SOP) regression between average niche overlap of C-N-P-cycling genes across the successional gradient.

**Table S13:**
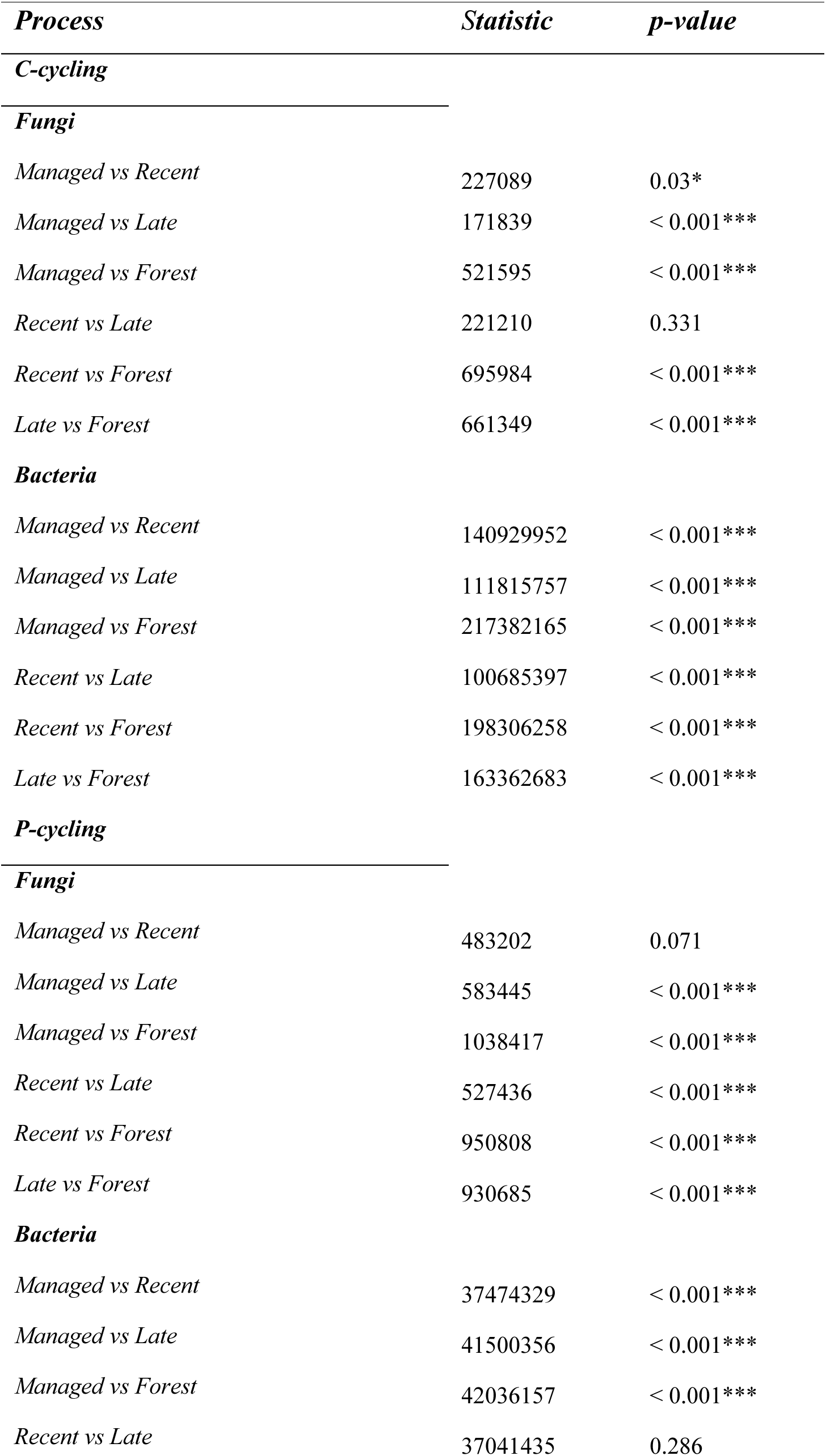
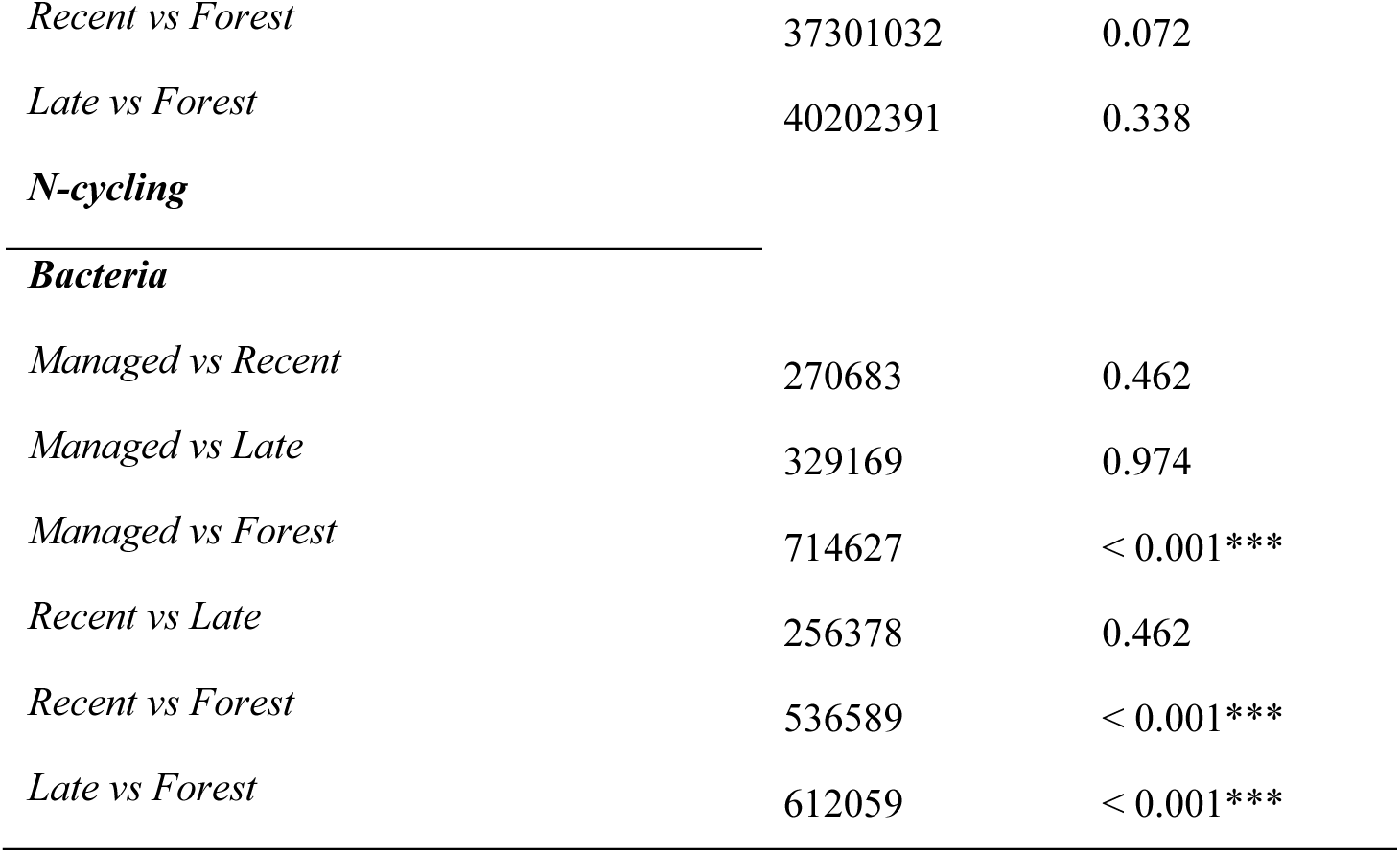
Results from pairwise Wilcoxon Rank-Sum tests for differences in average niche overlap of genes related to nutrient cycling processes between land uses across the successional gradient. *P-*values have been adjusted using Benjamin-Hochberg correction for multiple testing.

**Table S14:**
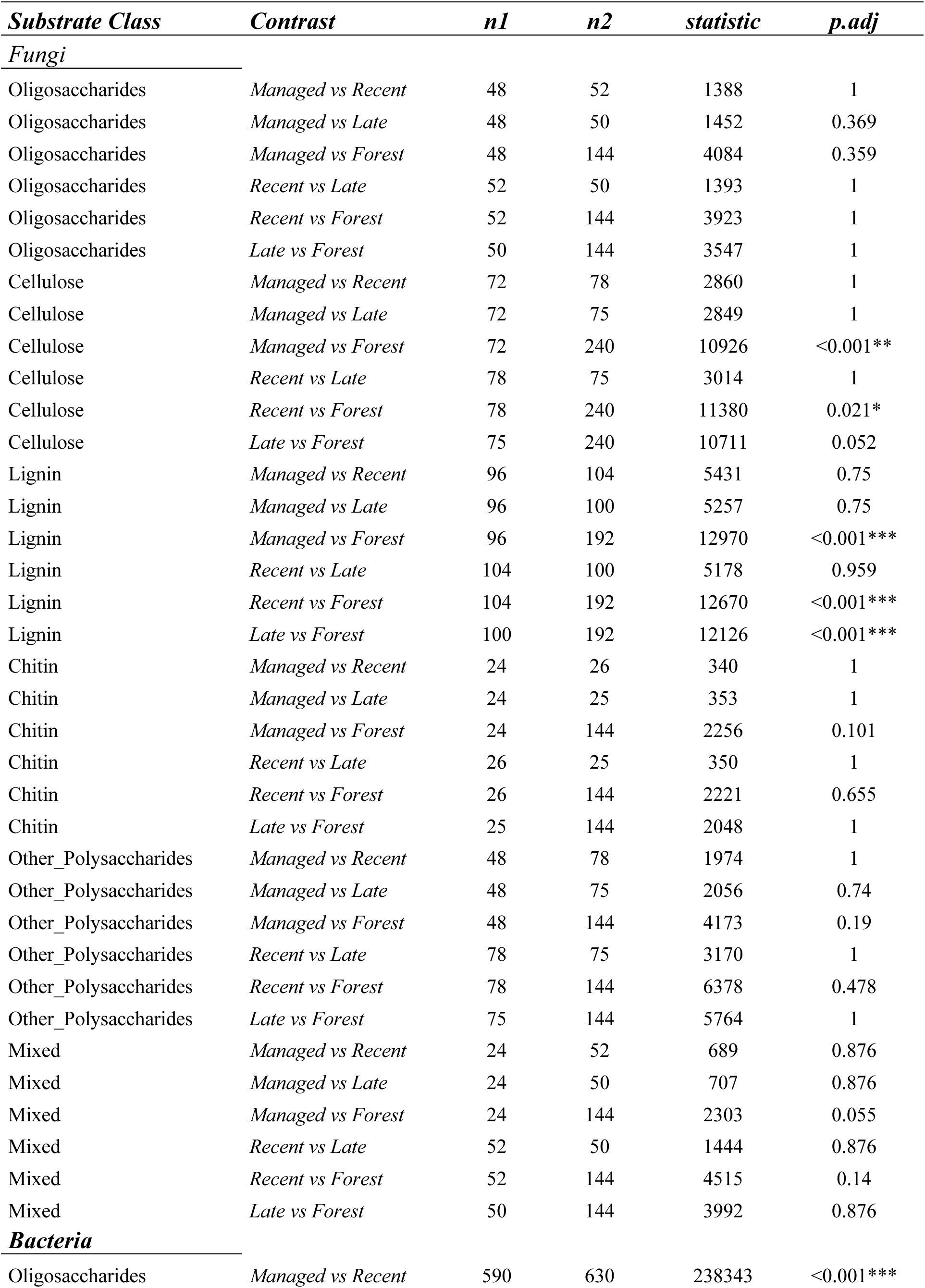
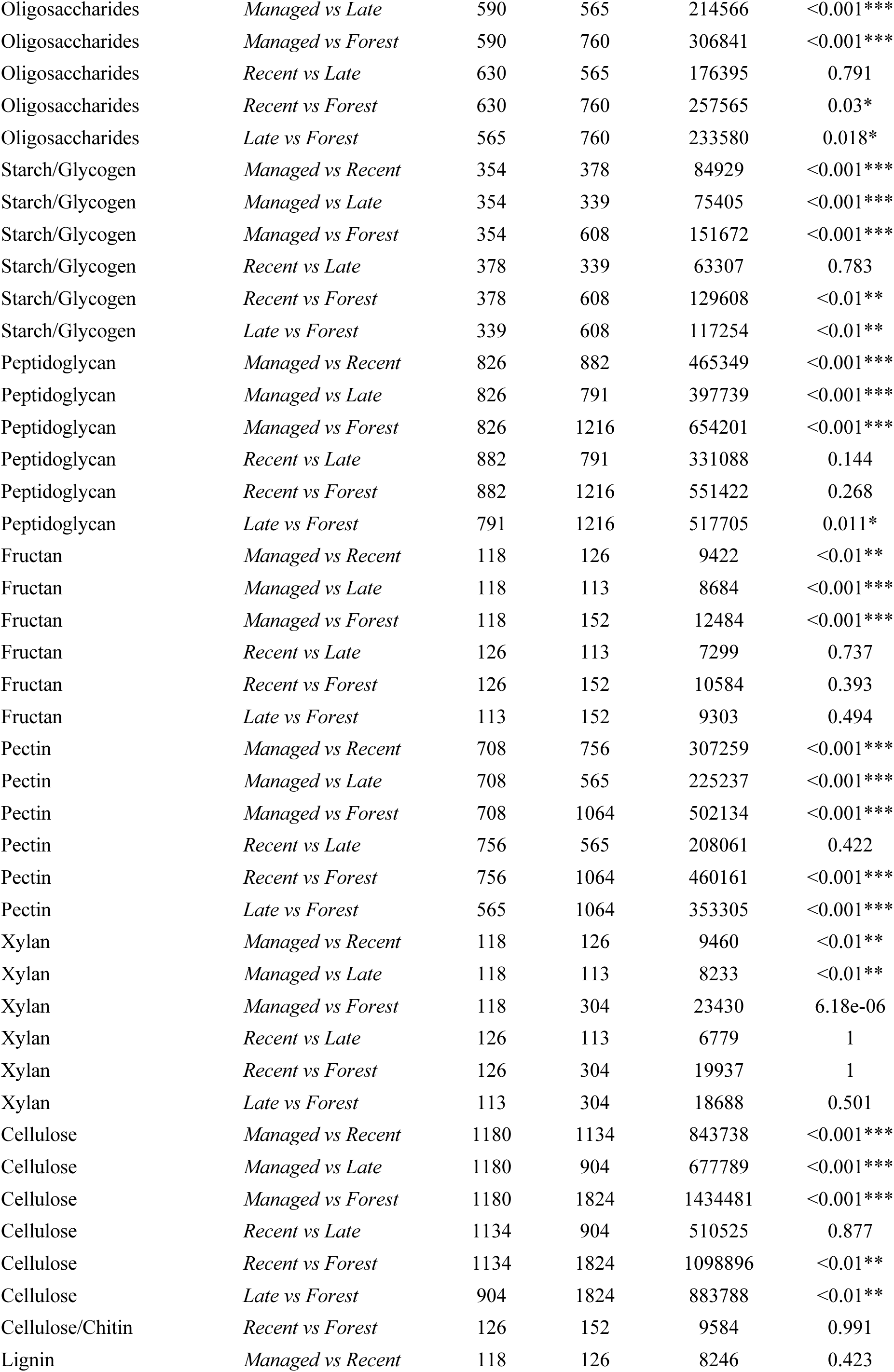
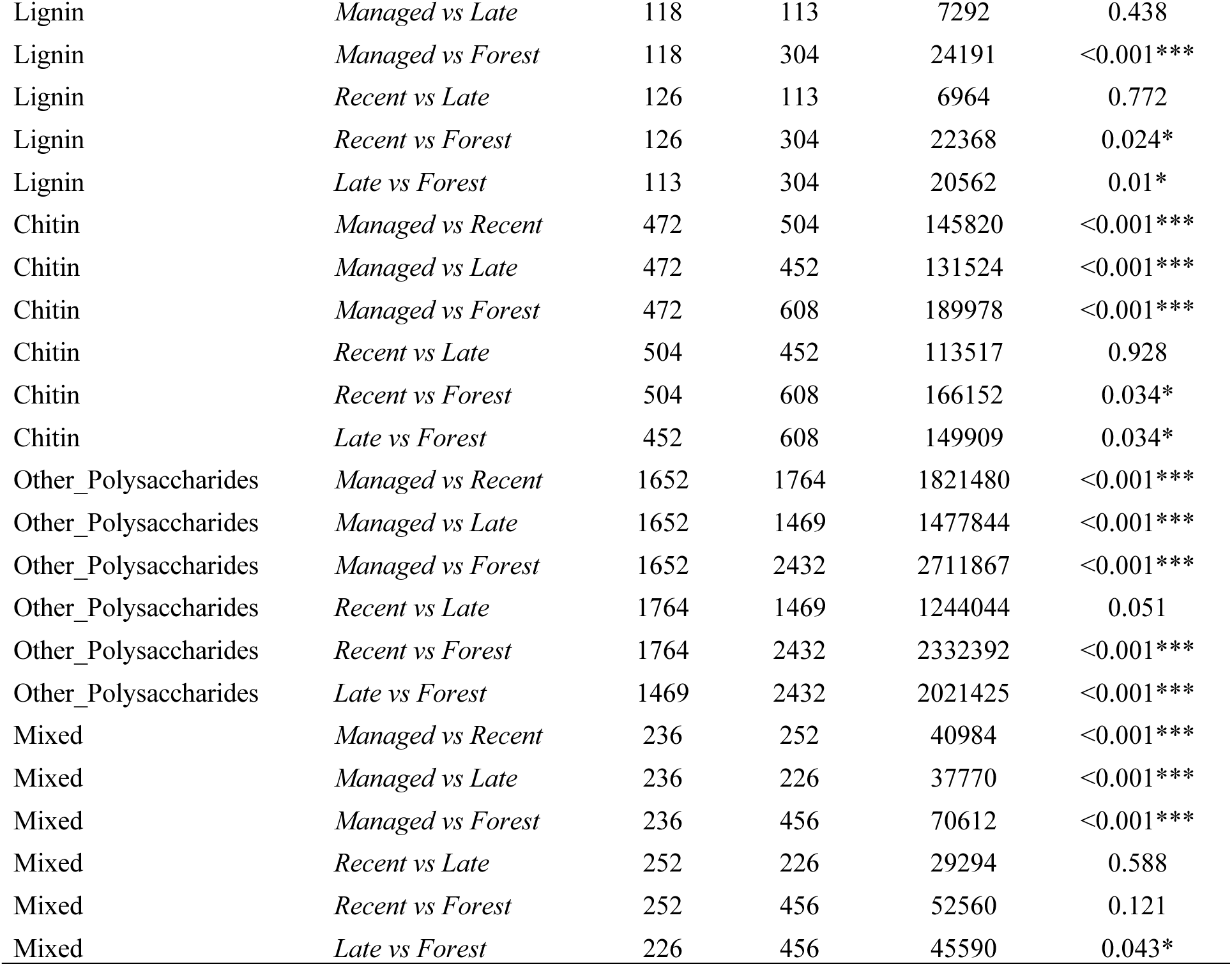
Results from pairwise Wilcoxon Rank-Sum tests for differences in average niche overlap of C-cycling genes partitioned across substrate classes. *P-*values adjusted using Benjamin-Hochberg correction for multiple testing.

**Table S15:**
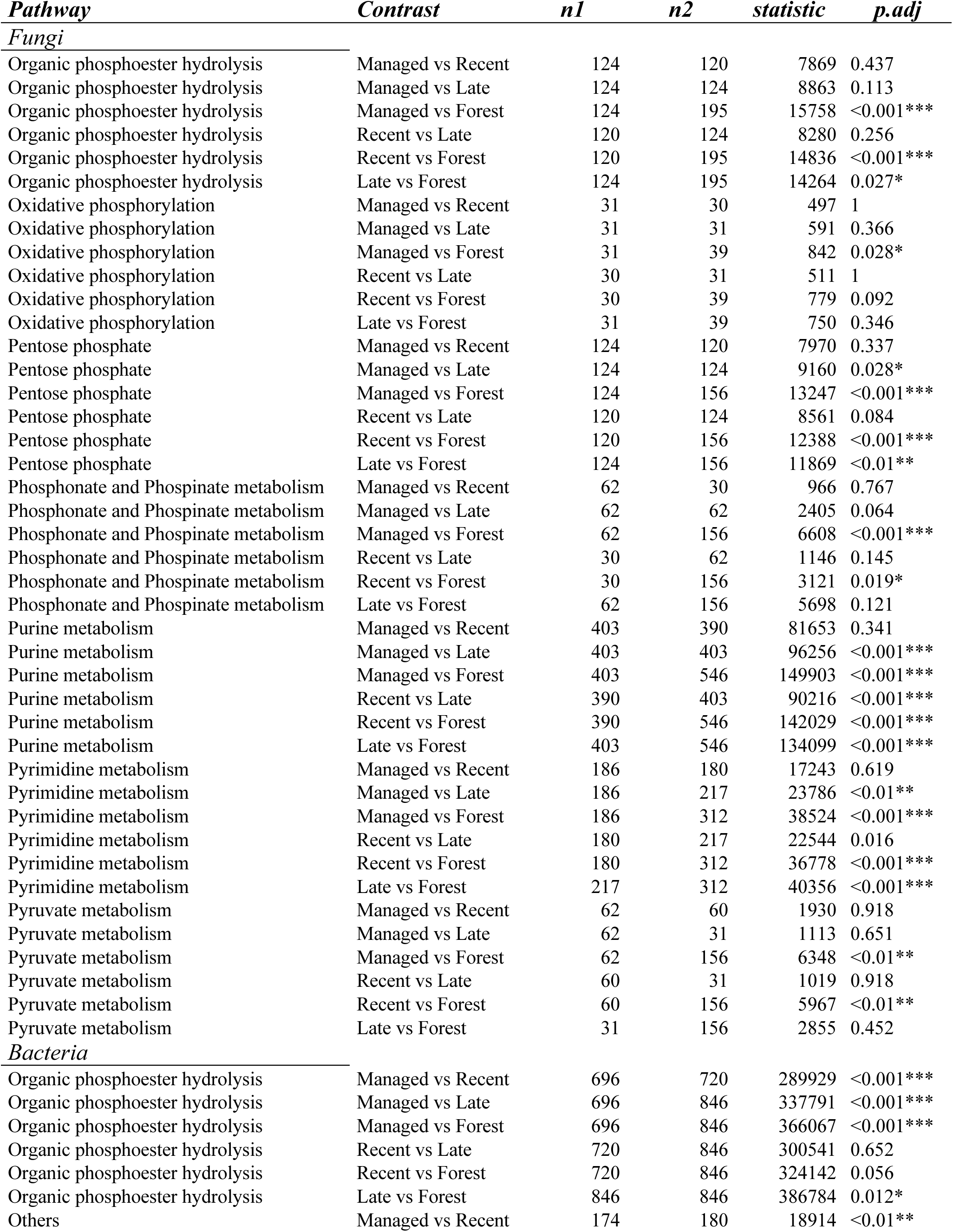
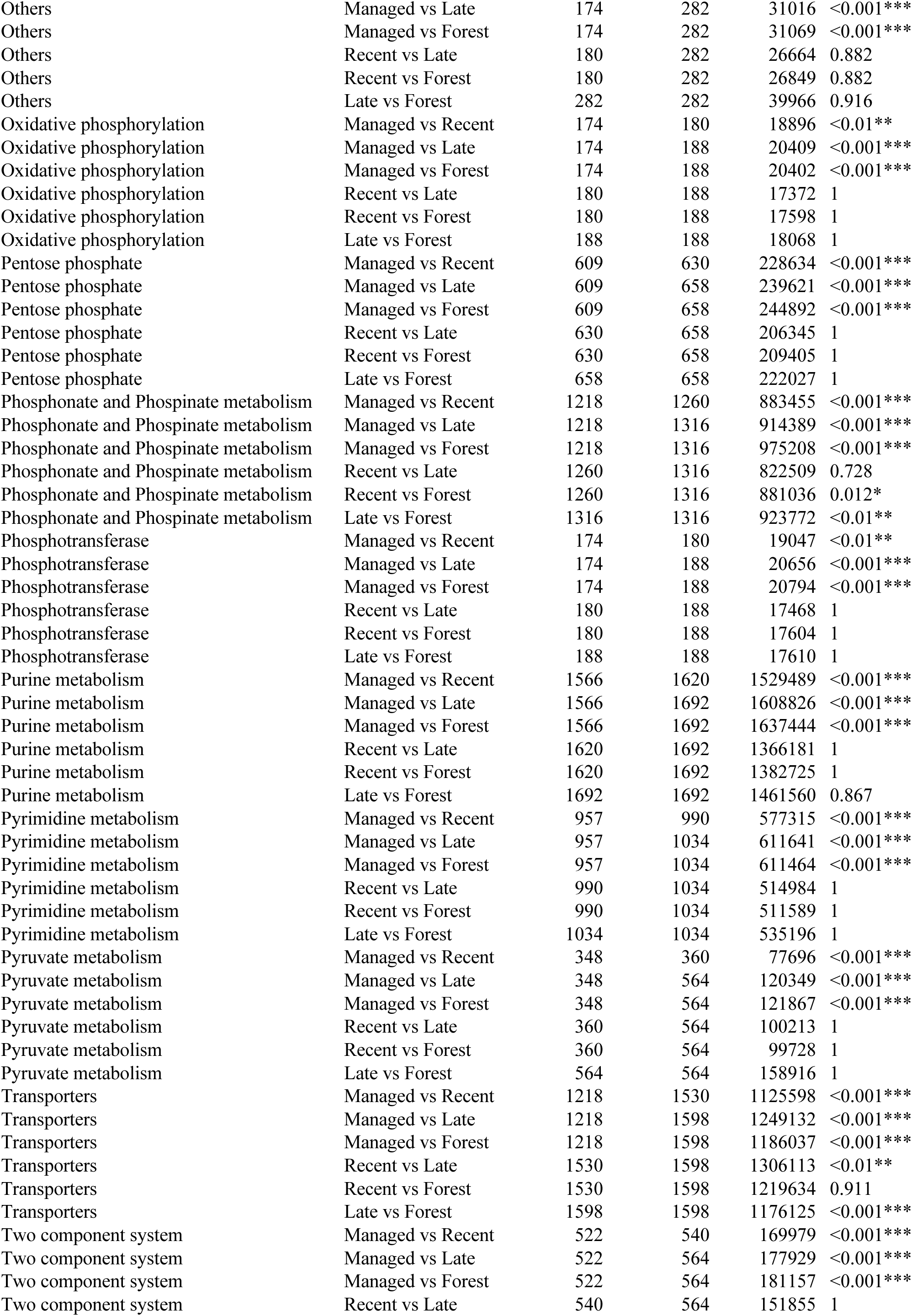
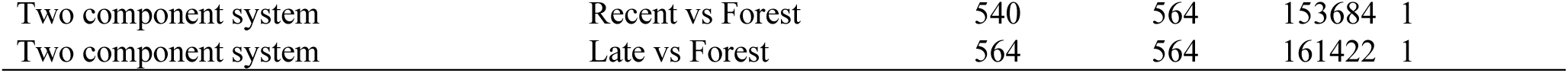
Results from pairwise Wilcoxon Rank-Sum tests for differences in average niche overlap of P-cycling genes partitioned across pathways. *P-*values adjusted using Benjamin-Hochberg correction for multiple testing.

**Table S16:**
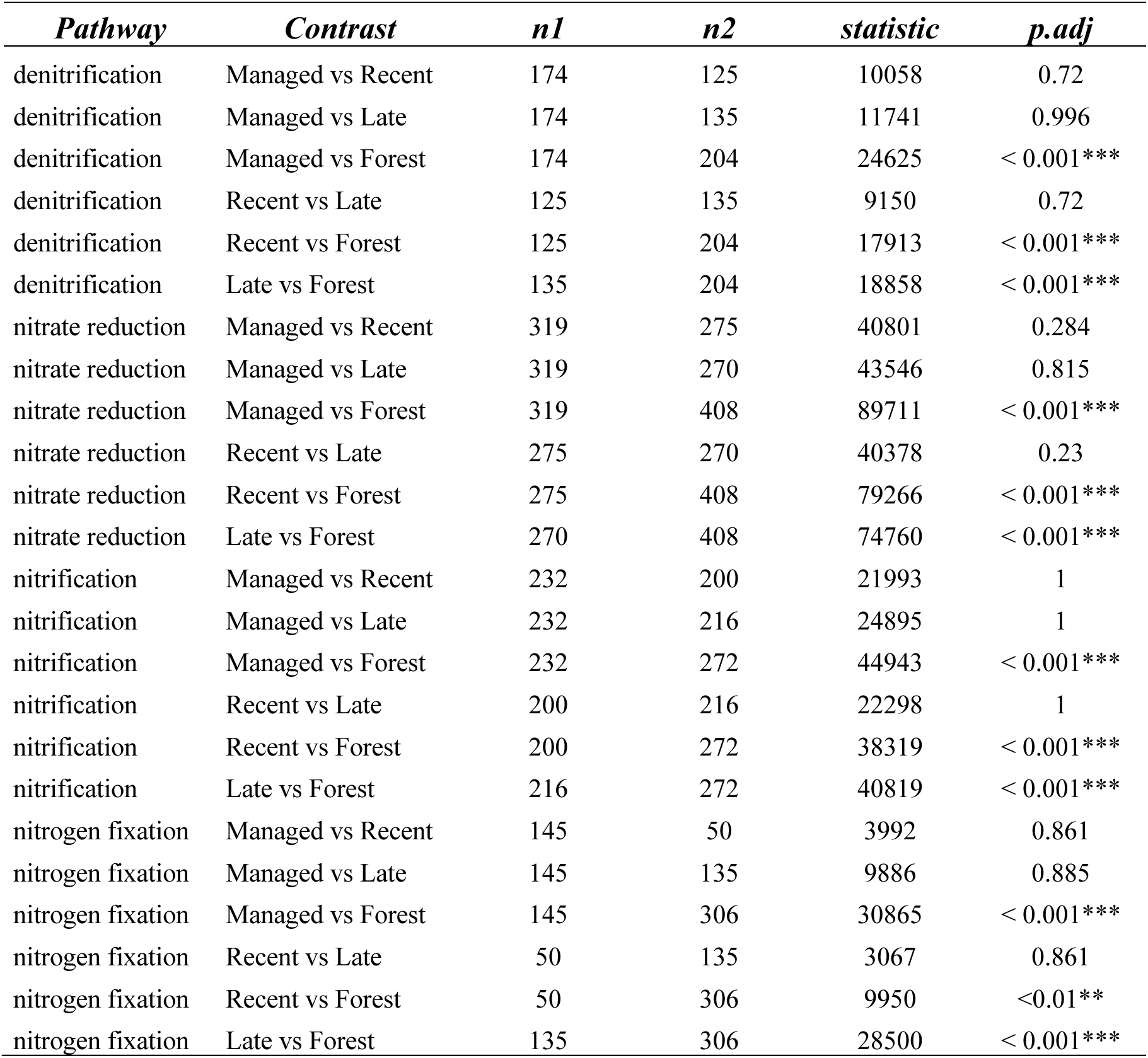
Results from pairwise Wilcoxon Rank-Sum tests for differences in average niche overlap of N-cycling genes partitioned across pathways. *P-*values adjusted using Benjamin-Hochberg correction for multiple testing.

**Table S17:**
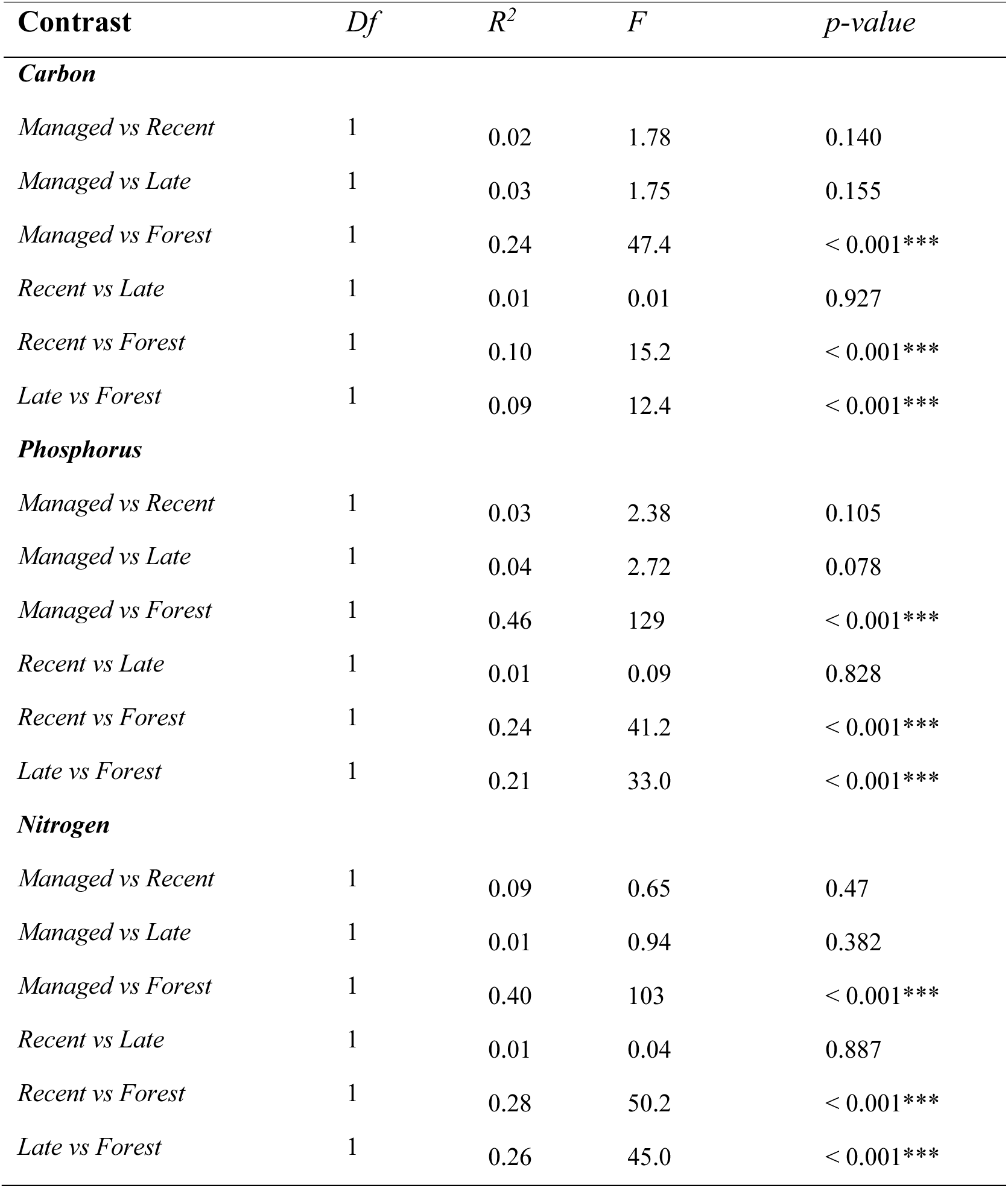
Differences in the composition of functional diversity, redundancy, and Simpson’s dominance of predicted bacterial metagenomes between successional stages. Results based on permutational multivariate tests (perMANOVA) with Bray-Curtis distances and 10^^4^ permutations

**Table S18:**
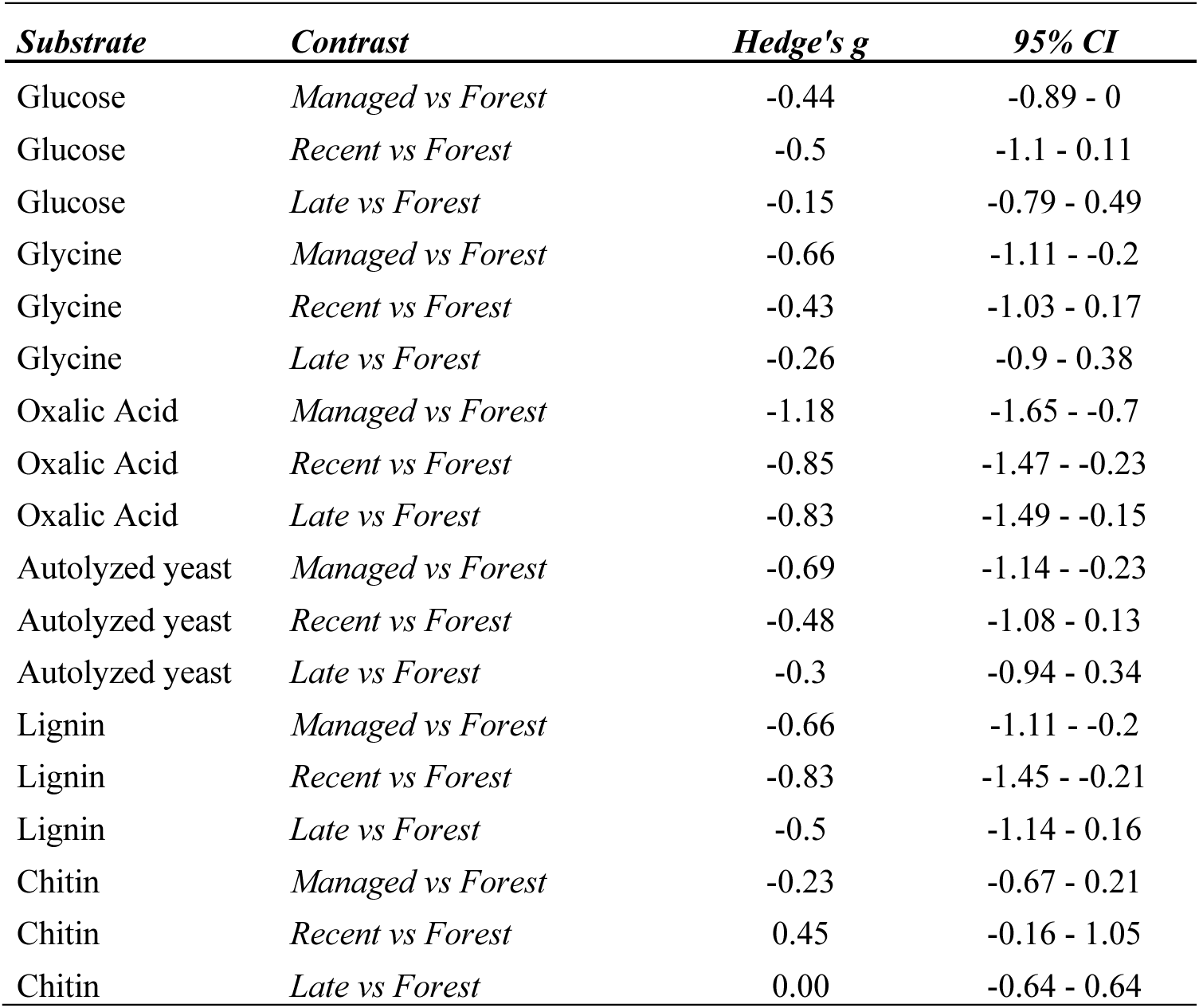
Effect sizes (*Hedge’s g*) of substrate-induced respiration rates between grassland sites at differing stages of succession compared to their respective paired forest sites. Negative effect sizes indicate higher respiration rates in forest sites.

