## Supplementary Materials for "Functional diversity of soil microbial communities increases with ecosystem development"

**SUPPLEMENTARY TABLES**

**Table S1**: Site distribution for each dataset and land-use category across the successional gradient

| **Dataset** | ***Managed*** | ***Recent*** | ***Late-term*** | ***Forest*** | ***Sum*** | ***%Paired*** |
| --- | --- | --- | --- | --- | --- | --- |
| Metadata | 49 | 30 | 23 | 105 | 207 | 92 |
| *Sequencing* |  |  |  |  |  |  |
| 16S (amplicon) | 48 | 29 | 22 | 105 | 204 | 90 |
| ITS (amplicon) | 47 | 26 | 22 | 104 | 199 | 88 |
| Shotgun | 16 | 18 | 14 | 47 | 95 | 99 |
| *Experiment* |  |  |  |  |  |  |
| SIR | 39 | 20 | 18 | 77 | 154 | 100 |

**Table S2:** Combined tests for differences in composition and indicator species contributing to observed differences between pairs of plant communities. All indicator species significant (*p* < 0.05), and all *p*-values adjusted for multiple comparisons (Benjamin-Hochberg)

| Comparison | *SES* | *p-value* | *Indicator species* |
| --- | --- | --- | --- |
| *Managed vs Recent* | 1.20 | 0.0072 | *Taraxacum;*  *T. repens; P. tremula* |
| *Managed vs Late* | 2.59 | < 0.0001 | *R. arcticus; R. ideaus; F. ulmaria; C. purpurea; C. fontanum; Taraxacum; A millefolium;*  *E. sylvaticum; P. lanceolata; L. vulgaris ; T. repens ; P. tremula ; B. pendula ; P. abies* |
| *Recent vs Late* | 1.53 | 0.0012 | *C. purpurea; A. millefolium; P. abies* |

**Table S3:** Soil and leaf properties (mean ± SD) across the successional gradient.

Letters indicate significant differences (*p* < 0.05) based on pairwise Wilcoxon tests.

| Variable | *Managed*  *(n = 49)* | *Early*  *(n = 30)* | *Late*  *(n = 23)* | *Forest*  *(n = 105)* |
| --- | --- | --- | --- | --- |
| *Soil properties* |  |  |  |  |
| *pH* | 5.38 (0.41)^a^ | 5.28 (0.28)^a^ | 5.29 (0.52)^a^ | 4.56 (0.54)^b^ |
| *Total C (%)* | 9.25 (10.4)^a^ | 9.06 (10.5)^a^ | 11.0 (10.5)^a^ | 17.9 (13.3)^b^ |
| *Total N (%)* | 0.60 (0.60) | 0.65 (0.73) | 0.74 (0.64) | 0.70 (0.52) |
| *C:N* | 14.4 (2.87)^a^ | 13.9 (2.96)^a^ | 14.4 (2.22)^a^ | 25.3 (6.66)^b^ |
| *Available K (mg kg^-1^)* | 19.1 (16.7) | 19.7 (13.0) | 17.1 (6.37) | 22.3 (13.1) |
| *Available P (mg kg^-1^)* | 6.82 (6.15) | 5.45 (3.29) | 7.54 (8.66) | 5.55 (3.12) |
| *Available Fe (mg kg^-1^)* | 50.3 (43.3)^a^ | 71.3 (86.7)^ab^ | 100 (97.3)^b^ | 45.7 (35.1)^a^ |
| *Available Mg (mg kg^-1^)* | 15.7 (10.7) | 17.5 (13.0) | 16.9 (12.2) | 18.2 (12.0) |
| *Available Al (mg kg^-1^)* | 70.0 (60.7) | 56.1 (32.8) | 81.0 (47.0) | 61.1 (47.2) |
| *Leaf dry matter content (LDMC)* | 0.27 (0.04)^a^ | 0.28 (0.04)^a^ | 0.31 (0.04)^b^ | 0.45 (0.02)^c^ |

**Table S4**: Number of reads and annotated orthologous groups (OG) related to nutrient cycling pathways obtained from the shotgun metagenomic sequencing of *n* = 95 samples across the successional gradient.

| **Dataset** | ***Reads*** | ***mean ± SD*** | ***OG*** |
| --- | --- | --- | --- |
| Total metagenome reads | 3532698106 | 37186296 ± 7365270 |  |
| *CAZyme* fungi | 637506 | 6710 ± 3720 | 236 |
| *CAZyme* bacteria | 60418032 | 635979 ± 131111 | 500 |
| P-cyc fungi | 216506 | 2279 ± 607 | 51 |
| P-cyc bacteria | 20928992 | 220503 ± 45853 | 51 |
| N-cyc bacteria | 997816 | 10503 ± 2972 | 46 |

**Table S5**: Differences in microbial taxonomic diversity (Shannon’s *H´*) between grassland and forest sites based on linear mixed models incorporating site pairs, spatial distance, and geological parent material as random factors.

| Contrast | *df* | *estimate* | *se* | *t-value* | *p-value* |
| --- | --- | --- | --- | --- | --- |
| *Fungi* |  |  |  |  |  |
| *Managed vs Forest* | 118 | 1.37 | 0.14 | 2.30 | 0.021* |
| *Recent vs Forest* | 142 | 1.54 | 0.17 | 2.53 | 0.014* |
| *Late vs Forest* | 145 | 1.18 | 0.19 | 0.37 | 0.371 |
| *Bacteria* |  |  |  |  |  |
| *Managed vs Forest* | 125 | 1.97 | 0.08 | 8.29 | < 0.001*** |
| *Recent vs Forest* | 148 | 1.66 | 0.10 | 4.82 | < 0.001*** |
| *Late vs Forest* | 139 | 1.65 | 0.11 | 4.40 | < 0.001*** |

**Table S6**: Differences in microbial taxonomic diversity (Shannon’s *H´*) between grasslands in differing successional stages based on linear mixed models incorporating geological parent material as random factor.

| Contrast | *df* | *estimate* | *se* | *t-value* | *p-value* |
| --- | --- | --- | --- | --- | --- |
| *Fungi* |  |  |  |  |  |
| *Managed vs Forest* | 118 | 1.37 | 0.14 | 2.30 | 0.021* |
| *Recent vs Forest* | 142 | 1.54 | 0.17 | 2.53 | 0.014* |
| *Late vs Forest* | 145 | 1.18 | 0.19 | 0.37 | 0.371 |
| *Bacteria* |  |  |  |  |  |
| *Managed vs Forest* | 125 | 1.97 | 0.08 | 8.29 | < 0.001*** |
| *Recent vs Forest* | 148 | 1.66 | 0.10 | 4.82 | < 0.001*** |
| *Late vs Forest* | 139 | 1.65 | 0.11 | 4.40 | < 0.001*** |

| Contrast | *estimate* | *se* | *z-value* | *p-value* |
| --- | --- | --- | --- | --- |
| *Fungi* |  |  |  |  |
| *Managed vs Recent* | 0.12 | 0.19 | 0.61 | 0.926 |
| *Managed vs Late* | 0.15 | 0.21 | 0.69 | 0.897 |
| *Recent vs Late* | 0.26 | 0.23 | 1.13 | 0.665 |
| *Bacteria* |  |  |  |  |
| *Managed vs Recent* | -0.17 | 0.12 | -1.47 | 0.445 |
| *Managed vs Late* | 0.18 | 0.13 | 1.15 | 0.478 |
| *Recent vs Late* | 0.01 | 0.14 | 0.03 | 1.000 |

**Table S7**: Results from permutational multivariate tests (perMANOVA) for differences in community composition between fungal and bacterial OTUs.

| *Contrast* | *Df* | *R^2^* | *F* | *p-value* |
| --- | --- | --- | --- | --- |
| *Fungi* |  |  |  |  |
| *Managed vs Recent* | 1 | 0.02 | 1.29 | 0.029* |
| *Managed vs Late* | 1 | 0.02 | 1.34 | 0.017* |
| *Managed vs Forest* | 1 | 0.06 | 5.14 | < 0.001*** |
| *Recent vs Late* | 1 | 0.03 | 2.06 | 0.061 |
| *Recent vs Forest* | 1 | 0.09 | 3.98 | < 0.001*** |
| *Late vs Forest* | 1 | 0.06 | 2.31 | < 0.001*** |
| *Bacteria* |  |  |  |  |
| *Managed vs Recent* | 1 | 0.01 | 0.92 | 0.616 |
| *Managed vs Late* | 1 | 0.02 | 1.24 | 0.294 |
| *Managed vs Forest* | 1 | 0.25 | 30.3 | < 0.001*** |
| *Recent vs Late* | 1 | 0.02 | 0.82 | 0.704 |
| *Recent vs Forest* | 1 | 0.20 | 11.6 | < 0.001*** |
| *Late vs Forest* | 1 | 0.14 | 6.47 | < 0.001*** |

**Table S8**: Differences in microbial biogeochemical genetic diversity (Shannon’s *H´*) between grasslands in differing successional stages based on linear mixed models incorporating geological parent material as random factor.

| Contrast | *estimate* | *se* | *z-value* | *p-value* |
| --- | --- | --- | --- | --- |
| *Carbon* |  |  |  |  |
| *Fungi* |  |  |  |  |
| *Managed vs Recent* | -0.08 | 0.12 | -0.64 | 0.917 |
| *Managed vs Late* | -0.08 | 0.12 | -0.71 | 0.890 |
| *Recent vs Late* | -0.01 | 0.11 | -0.05 | 0.999 |
| *Bacteria* |  |  |  |  |
| *Managed vs Recent* | 0.01 | 0.02 | 0.12 | 0.999 |
| *Managed vs Late* | 0.01 | 0.02 | 0.57 | 0.940 |
| *Recent vs Late* | 0.01 | 0.02 | 0.69 | 0.895 |
| *Phosphorus* |  |  |  |  |
| *Managed vs Recent* | -0.01 | 0.01 | -0.36 | 0.983 |
| *Managed vs Late* | -0.01 | 0.02 | -0.15 | 0.999 |
| *Recent vs Late* | -0.01 | 0.02 | -0.49 | 0.961 |
| *Bacteria* |  |  |  |  |
| *Managed vs Recent* | 0.01 | 0.01 | 0.02 | 1.000 |
| *Managed vs Late* | 0.01 | 0.01 | 0.21 | 0.997 |
| *Recent vs Late* | 0.01 | 0.01 | 0.23 | 0.995 |
| *Nitrogen* |  |  |  |  |
| *Managed vs Recent* | -0.01 | 0.06 | -0.16 | 0.998 |
| *Managed vs Late* | -0.05 | 0.07 | -0.71 | 0.892 |
| *Recent vs Late* | -0.06 | 0.06 | -0.83 | 0.809 |

**Table S9**: Differences in microbial functional diversity (Shannon’s *H´*) between grassland and forest sites based on linear mixed models incorporating site pairs, spatial distance, and geological parent material as random factors.

| Contrast | *df* | *estimate* | *se* | *t-value* | *p-value* |
| --- | --- | --- | --- | --- | --- |
| *Carbon* |  |  |  |  |  |
| *Fungi* |  |  |  |  |  |
| *Managed vs Forest* | 90 | 0.62 | 0.09 | -5.07 | < 0.001*** |
| *Recent vs Forest* | 91 | 0.61 | 0.09 | -5.39 | < 0.001*** |
| *Late vs Forest* | 75 | 0.66 | 0.10 | -3.98 | < 0.001*** |
| *Bacteria* |  |  |  |  |  |
| *Managed vs Forest* | 68 | 1.02 | 0.01 | 1.14 | 0.26 |
| *Recent vs Forest* | 68 | 1.02 | 0.01 | 1.44 | 0.15 |
| *Late vs Forest* | 70 | 1.00 | 0.02 | 0.26 | 0.79 |
| *Phosphorus* |  |  |  |  |  |
| *Fungi* |  |  |  |  |  |
| *Managed vs Forest* | 68 | 0.99 | 0.01 | -1.96 | 0.05 |
| *Recent vs Forest* | 68 | 0.99 | 0.01 | -1.87 | 0.07 |
| *Late vs Forest* | 71 | 0.99 | 0.01 | -1.74 | 0.09 |
| *Bacteria* |  |  |  |  |  |
| *Managed vs Forest* | 87 | 1.01 | 0.02 | 0.62 | 0.53 |
| *Recent vs Forest* | 90 | 1.01 | 0.02 | 0.56 | 0.58 |
| *Late vs Forest* | 58 | 1.02 | 0.02 | 0.88 | 0.38 |
| *Nitrogen* |  |  |  |  |  |
| *Bacteria* |  |  |  |  |  |
| *Managed vs Forest* | 68 | 1.30 | 0.05 | 5.17 | < 0.001*** |
| *Recent vs Forest* | 68 | 1.29 | 0.05 | 5.31 | < 0.001*** |
| *Late vs Forest* | 61 | 1.35 | 0.05 | 5.48 | < 0.001*** |

**Table S10**: Relationship between taxonomic (Shannon’s *H´* of OTU matrices) and functional (Shannon’s *H´* of functional matrices) diversity for microbial communities between grassland successional stages, including when divided by ecosystem type (grassland, forest). Results based on ordinary least-square (OLS) regression. For bacterial communities, the average genome size (AGS) was included as a covariate in the regressions.

|  | estimate | se | t-value | p-value | p-value AGS | R^2^ |
| --- | --- | --- | --- | --- | --- | --- |
| *Overall community* |  |  |  |  |  |  |
| Fungal C-cycling | -0.16 | 0.06 | -2.86 | 0.005** | NA | 0.07 |
| Fungal P-cycling | 0.01 | 0.01 | 0.53 | 0.597 | NA | 0.01 |
| Bacterial C-cycling | 0.03 | 0.01 | 2.86 | 0.005** | < 0.001*** | 0.14 |
| Bacterial P-cycling | 0.03 | 0.01 | 6.29 | < 0.001*** | < 0.001*** | 0.31 |
| Bacterial N-cycling | 0.16 | 0.04 | 4.17 | < 0.001*** | 0.901 | 0.17 |
| *Grasslands* |  |  |  |  |  |  |
| Fungal C-cycling | -0.04 | 0.09 | -0.51 | 0.613 | NA | -0.02 |
| Fungal P-cycling | 0.01 | 0.01 | 0.03 | 0.978 | NA | -0.02 |
| Bacterial C-cycling | 0.02 | 0.02 | 0.87 | 0.392 | 0.018* | 0.10 |
| Bacterial P-cycling | 0.03 | 0.01 | 5.72 | < 0.001*** | < 0.001*** | 0.41 |
| Bacterial N-cycling | -0.04 | 0.04 | -0.97 | 0.339 | 0.037* | 0.06 |
| *Forests* |  |  |  |  |  |  |
| Fungal C-cycling | -0.13 | 0.06 | -2.37 | 0.022* | NA | 0.10 |
| Fungal P-cycling | 0.01 | 0.01 | 0.48 | 0.631 | NA | -0.02 |
| Bacterial C-cycling | 0.04 | 0.01 | 2.99 | < 0.01** | 0.012* | 0.20 |
| Bacterial P-cycling | 0.04 | 0.01 | 6.55 | < 0.001*** | 0.002** | 0.49 |
| Bacterial N-cycling | 0.21 | 0.06 | 3.61 | < 0.001*** | 0.456 | 0.19 |

**Table S11**: Pearson correlation tests between taxonomic (Shannon’s *H´* of OUT matrices) and functional (Shannon’s *H´* of functional matrices) diversity for microbial communities divided by ecosystem type (grassland, forest).

| *Contrast* | *Df* | *t* | *r* | *p-value* | 95% CI |
| --- | --- | --- | --- | --- | --- |
| *Grasslands* |  |  |  |  |  |
| Fungal C-cycling | 43 | -0.51 | -0.08 | 0.613 | -0.36 - 0.22 |
| Fungal P-cycling | 43 | 0.03 | 0.01 | 0.978 | -0.29 – 0.30 |
| Bacterial C-cycling | 45 | -0.74 | -0.11 | 0.466 | -0.39 – 0.19 |
| Bacterial P-cycling | 45 | 3.69 | 0.49 | < 0.001*** | 0.23 – 0.68 |
| Bacterial N-cycling | 45 | 0.40 | 0.06 | 0.691 | -0.24 – 0.35 |
| *Forests* |  |  |  |  |  |
| Fungal C-cycling | 43 | -2.37 | -0.34 | 0.022* | -0.58 – -0.05 |
| Fungal P-cycling | 43 | 0.48 | 0.07 | 0.631 | -0.22 – 0.36 |
| Bacterial C-cycling | 45 | 2.39 | 0.34 | 0.021* | 0.05 – 0.57 |
| Bacterial P-cycling | 45 | 5.50 | 0.63 | < 0.001*** | 0.42 – 0.78 |
| Bacterial N-cycling | 45 | 3.54 | 0.47 | < 0.001*** | 0.21 – 0.67 |

**Table S12**: Results from ordinary least-square regression (OLS) and second-order polynomial (SOP) regression between average niche overlap of C-N-P-cycling genes across the successional gradient

| *Process* | *estimate* | *se* | *t-value* | *p-value* |
| --- | --- | --- | --- | --- |
| *C-cycling* |  |  |  |  |
| *Fungi OLS* | -0.01 | 0.01 | -6.66 | < 0.001*** |
| *Bacteria OLS* | -0.07 | 0.01 | -33.0 | < 0.001*** |
| *Bacteria SOP* | 0.01 | 0.01 | 28.2 | < 0.001*** |
| *P-cycling* |  |  |  |  |
| *Fungi OLS* | 0.04 | 0.01 | 4.51 | < 0.001*** |
| *Fungi SOP* | -0.02 | 0.01 | -8.30 | < 0.001*** |
| *Bacteria OLS* | -0.01 | 0.01 | -19.3 | < 0.001*** |
| *N-cycling* |  |  |  |  |
| *Bacteria OLS* | 0.15 | 0.02 | 8.09 | < 0.001*** |
| *Bacteria SOP* | -0.04 | 0.03 | -11.2 | < 0.001*** |

**Table S13**: Results from pairwise Wilcoxon Rank-Sum tests for differences in average niche overlap of genes related to nutrient cycling processes between land uses across the successional gradient. *P-*values have been adjusted using Benjamin-Hochberg correction for multiple testing.

| *Process* | *Statistic* | *p-value* |
| --- | --- | --- |
| *C-cycling* |  |  |
| *Fungi* |  |  |
| *Managed vs Recent* | 227089 | 0.03* |
| *Managed vs Late* | 171839 | < 0.001*** |
| *Managed vs Forest* | 521595 | < 0.001*** |
| *Recent vs Late* | 221210 | 0.331 |
| *Recent vs Forest* | 695984 | < 0.001*** |
| *Late vs Forest* | 661349 | < 0.001*** |
| *Bacteria* |  |  |
| *Managed vs Recent* | 140929952 | < 0.001*** |
| *Managed vs Late* | 111815757 | < 0.001*** |
| *Managed vs Forest* | 217382165 | < 0.001*** |
| *Recent vs Late* | 100685397 | < 0.001*** |
| *Recent vs Forest* | 198306258 | < 0.001*** |
| *Late vs Forest* | 163362683 | < 0.001*** |
| *P-cycling* |  |  |
| *Fungi* |  |  |
| *Managed vs Recent* | 483202 | 0.071 |
| *Managed vs Late* | 583445 | < 0.001*** |
| *Managed vs Forest* | 1038417 | < 0.001*** |
| *Recent vs Late* | 527436 | < 0.001*** |
| *Recent vs Forest* | 950808 | < 0.001*** |
| *Late vs Forest* | 930685 | < 0.001*** |
| *Bacteria* |  |  |
| *Managed vs Recent* | 37474329 | < 0.001*** |
| *Managed vs Late* | 41500356 | < 0.001*** |
| *Managed vs Forest* | 42036157 | < 0.001*** |
| *Recent vs Late* | 37041435 | 0.286 |
| *Recent vs Forest* | 37301032 | 0.072 |
| *Late vs Forest* | 40202391 | 0.338 |
| *N-cycling* |  |  |
| *Bacteria* |  |  |
| *Managed vs Recent* | 270683 | 0.462 |
| *Managed vs Late* | 329169 | 0.974 |
| *Managed vs Forest* | 714627 | < 0.001*** |
| *Recent vs Late* | 256378 | 0.462 |
| *Recent vs Forest* | 536589 | < 0.001*** |
| *Late vs Forest* | 612059 | < 0.001*** |

**Table S14**: Results from pairwise Wilcoxon Rank-Sum tests for differences in average niche overlap of C-cycling genes partitioned across substrate classes. *P-*values adjusted using Benjamin-Hochberg correction for multiple testing.

| ***Substrate Class*** | ***Contrast*** | ***n1*** | ***n2*** | ***statistic*** | ***p.adj*** |
| --- | --- | --- | --- | --- | --- |
| *Fungi* |  |  |  |  |  |
| Oligosaccharides | *Managed vs Recent* | 48 | 52 | 1388 | 1 |
| Oligosaccharides | *Managed vs Late* | 48 | 50 | 1452 | 0.369 |
| Oligosaccharides | *Managed vs Forest* | 48 | 144 | 4084 | 0.359 |
| Oligosaccharides | *Recent vs Late* | 52 | 50 | 1393 | 1 |
| Oligosaccharides | *Recent vs Forest* | 52 | 144 | 3923 | 1 |
| Oligosaccharides | *Late vs Forest* | 50 | 144 | 3547 | 1 |
| Cellulose | *Managed vs Recent* | 72 | 78 | 2860 | 1 |
| Cellulose | *Managed vs Late* | 72 | 75 | 2849 | 1 |
| Cellulose | *Managed vs Forest* | 72 | 240 | 10926 | <0.001** |
| Cellulose | *Recent vs Late* | 78 | 75 | 3014 | 1 |
| Cellulose | *Recent vs Forest* | 78 | 240 | 11380 | 0.021* |
| Cellulose | *Late vs Forest* | 75 | 240 | 10711 | 0.052 |
| Lignin | *Managed vs Recent* | 96 | 104 | 5431 | 0.75 |
| Lignin | *Managed vs Late* | 96 | 100 | 5257 | 0.75 |
| Lignin | *Managed vs Forest* | 96 | 192 | 12970 | <0.001*** |
| Lignin | *Recent vs Late* | 104 | 100 | 5178 | 0.959 |
| Lignin | *Recent vs Forest* | 104 | 192 | 12670 | <0.001*** |
| Lignin | *Late vs Forest* | 100 | 192 | 12126 | <0.001*** |
| Chitin | *Managed vs Recent* | 24 | 26 | 340 | 1 |
| Chitin | *Managed vs Late* | 24 | 25 | 353 | 1 |
| Chitin | *Managed vs Forest* | 24 | 144 | 2256 | 0.101 |
| Chitin | *Recent vs Late* | 26 | 25 | 350 | 1 |
| Chitin | *Recent vs Forest* | 26 | 144 | 2221 | 0.655 |
| Chitin | *Late vs Forest* | 25 | 144 | 2048 | 1 |
| Other_Polysaccharides | *Managed vs Recent* | 48 | 78 | 1974 | 1 |
| Other_Polysaccharides | *Managed vs Late* | 48 | 75 | 2056 | 0.74 |
| Other_Polysaccharides | *Managed vs Forest* | 48 | 144 | 4173 | 0.19 |
| Other_Polysaccharides | *Recent vs Late* | 78 | 75 | 3170 | 1 |
| Other_Polysaccharides | *Recent vs Forest* | 78 | 144 | 6378 | 0.478 |
| Other_Polysaccharides | *Late vs Forest* | 75 | 144 | 5764 | 1 |
| Mixed | *Managed vs Recent* | 24 | 52 | 689 | 0.876 |
| Mixed | *Managed vs Late* | 24 | 50 | 707 | 0.876 |
| Mixed | *Managed vs Forest* | 24 | 144 | 2303 | 0.055 |
| Mixed | *Recent vs Late* | 52 | 50 | 1444 | 0.876 |
| Mixed | *Recent vs Forest* | 52 | 144 | 4515 | 0.14 |
| Mixed | *Late vs Forest* | 50 | 144 | 3992 | 0.876 |
| ***Bacteria*** |  |  |  |  |  |
| Oligosaccharides | *Managed vs Recent* | 590 | 630 | 238343 | <0.001*** |
| Oligosaccharides | *Managed vs Late* | 590 | 565 | 214566 | <0.001*** |
| Oligosaccharides | *Managed vs Forest* | 590 | 760 | 306841 | <0.001*** |
| Oligosaccharides | *Recent vs Late* | 630 | 565 | 176395 | 0.791 |
| Oligosaccharides | *Recent vs Forest* | 630 | 760 | 257565 | 0.03* |
| Oligosaccharides | *Late vs Forest* | 565 | 760 | 233580 | 0.018* |
| Starch/Glycogen | *Managed vs Recent* | 354 | 378 | 84929 | <0.001*** |
| Starch/Glycogen | *Managed vs Late* | 354 | 339 | 75405 | <0.001*** |
| Starch/Glycogen | *Managed vs Forest* | 354 | 608 | 151672 | <0.001*** |
| Starch/Glycogen | *Recent vs Late* | 378 | 339 | 63307 | 0.783 |
| Starch/Glycogen | *Recent vs Forest* | 378 | 608 | 129608 | <0.01** |
| Starch/Glycogen | *Late vs Forest* | 339 | 608 | 117254 | <0.01** |
| Peptidoglycan | *Managed vs Recent* | 826 | 882 | 465349 | <0.001*** |
| Peptidoglycan | *Managed vs Late* | 826 | 791 | 397739 | <0.001*** |
| Peptidoglycan | *Managed vs Forest* | 826 | 1216 | 654201 | <0.001*** |
| Peptidoglycan | *Recent vs Late* | 882 | 791 | 331088 | 0.144 |
| Peptidoglycan | *Recent vs Forest* | 882 | 1216 | 551422 | 0.268 |
| Peptidoglycan | *Late vs Forest* | 791 | 1216 | 517705 | 0.011* |
| Fructan | *Managed vs Recent* | 118 | 126 | 9422 | <0.01** |
| Fructan | *Managed vs Late* | 118 | 113 | 8684 | <0.001*** |
| Fructan | *Managed vs Forest* | 118 | 152 | 12484 | <0.001*** |
| Fructan | *Recent vs Late* | 126 | 113 | 7299 | 0.737 |
| Fructan | *Recent vs Forest* | 126 | 152 | 10584 | 0.393 |
| Fructan | *Late vs Forest* | 113 | 152 | 9303 | 0.494 |
| Pectin | *Managed vs Recent* | 708 | 756 | 307259 | <0.001*** |
| Pectin | *Managed vs Late* | 708 | 565 | 225237 | <0.001*** |
| Pectin | *Managed vs Forest* | 708 | 1064 | 502134 | <0.001*** |
| Pectin | *Recent vs Late* | 756 | 565 | 208061 | 0.422 |
| Pectin | *Recent vs Forest* | 756 | 1064 | 460161 | <0.001*** |
| Pectin | *Late vs Forest* | 565 | 1064 | 353305 | <0.001*** |
| Xylan | *Managed vs Recent* | 118 | 126 | 9460 | <0.01** |
| Xylan | *Managed vs Late* | 118 | 113 | 8233 | <0.01** |
| Xylan | *Managed vs Forest* | 118 | 304 | 23430 | 6.18e-06 |
| Xylan | *Recent vs Late* | 126 | 113 | 6779 | 1 |
| Xylan | *Recent vs Forest* | 126 | 304 | 19937 | 1 |
| Xylan | *Late vs Forest* | 113 | 304 | 18688 | 0.501 |
| Cellulose | *Managed vs Recent* | 1180 | 1134 | 843738 | <0.001*** |
| Cellulose | *Managed vs Late* | 1180 | 904 | 677789 | <0.001*** |
| Cellulose | *Managed vs Forest* | 1180 | 1824 | 1434481 | <0.001*** |
| Cellulose | *Recent vs Late* | 1134 | 904 | 510525 | 0.877 |
| Cellulose | *Recent vs Forest* | 1134 | 1824 | 1098896 | <0.01** |
| Cellulose | *Late vs Forest* | 904 | 1824 | 883788 | <0.01** |
| Cellulose/Chitin | *Recent vs Forest* | 126 | 152 | 9584 | 0.991 |
| Lignin | *Managed vs Recent* | 118 | 126 | 8246 | 0.423 |
| Lignin | *Managed vs Late* | 118 | 113 | 7292 | 0.438 |
| Lignin | *Managed vs Forest* | 118 | 304 | 24191 | <0.001*** |
| Lignin | *Recent vs Late* | 126 | 113 | 6964 | 0.772 |
| Lignin | *Recent vs Forest* | 126 | 304 | 22368 | 0.024* |
| Lignin | *Late vs Forest* | 113 | 304 | 20562 | 0.01* |
| Chitin | *Managed vs Recent* | 472 | 504 | 145820 | <0.001*** |
| Chitin | *Managed vs Late* | 472 | 452 | 131524 | <0.001*** |
| Chitin | *Managed vs Forest* | 472 | 608 | 189978 | <0.001*** |
| Chitin | *Recent vs Late* | 504 | 452 | 113517 | 0.928 |
| Chitin | *Recent vs Forest* | 504 | 608 | 166152 | 0.034* |
| Chitin | *Late vs Forest* | 452 | 608 | 149909 | 0.034* |
| Other_Polysaccharides | *Managed vs Recent* | 1652 | 1764 | 1821480 | <0.001*** |
| Other_Polysaccharides | *Managed vs Late* | 1652 | 1469 | 1477844 | <0.001*** |
| Other_Polysaccharides | *Managed vs Forest* | 1652 | 2432 | 2711867 | <0.001*** |
| Other_Polysaccharides | *Recent vs Late* | 1764 | 1469 | 1244044 | 0.051 |
| Other_Polysaccharides | *Recent vs Forest* | 1764 | 2432 | 2332392 | <0.001*** |
| Other_Polysaccharides | *Late vs Forest* | 1469 | 2432 | 2021425 | <0.001*** |
| Mixed | *Managed vs Recent* | 236 | 252 | 40984 | <0.001*** |
| Mixed | *Managed vs Late* | 236 | 226 | 37770 | <0.001*** |
| Mixed | *Managed vs Forest* | 236 | 456 | 70612 | <0.001*** |
| Mixed | *Recent vs Late* | 252 | 226 | 29294 | 0.588 |
| Mixed | *Recent vs Forest* | 252 | 456 | 52560 | 0.121 |
| Mixed | *Late vs Forest* | 226 | 456 | 45590 | 0.043* |

**Table S15**: Results from pairwise Wilcoxon Rank-Sum tests for differences in average niche overlap of P-cycling genes partitioned across pathways. *P-*values adjusted using Benjamin-Hochberg correction for multiple testing.

| ***Pathway*** | ***Contrast*** | ***n1*** | ***n2*** | ***statistic*** | ***p.adj*** |
| --- | --- | --- | --- | --- | --- |
| *Fungi* |  |  |  |  |  |
| Organic phosphoester hydrolysis | Managed vs Recent | 124 | 120 | 7869 | 0.437 |
| Organic phosphoester hydrolysis | Managed vs Late | 124 | 124 | 8863 | 0.113 |
| Organic phosphoester hydrolysis | Managed vs Forest | 124 | 195 | 15758 | <0.001*** |
| Organic phosphoester hydrolysis | Recent vs Late | 120 | 124 | 8280 | 0.256 |
| Organic phosphoester hydrolysis | Recent vs Forest | 120 | 195 | 14836 | <0.001*** |
| Organic phosphoester hydrolysis | Late vs Forest | 124 | 195 | 14264 | 0.027* |
| Oxidative phosphorylation | Managed vs Recent | 31 | 30 | 497 | 1 |
| Oxidative phosphorylation | Managed vs Late | 31 | 31 | 591 | 0.366 |
| Oxidative phosphorylation | Managed vs Forest | 31 | 39 | 842 | 0.028* |
| Oxidative phosphorylation | Recent vs Late | 30 | 31 | 511 | 1 |
| Oxidative phosphorylation | Recent vs Forest | 30 | 39 | 779 | 0.092 |
| Oxidative phosphorylation | Late vs Forest | 31 | 39 | 750 | 0.346 |
| Pentose phosphate | Managed vs Recent | 124 | 120 | 7970 | 0.337 |
| Pentose phosphate | Managed vs Late | 124 | 124 | 9160 | 0.028* |
| Pentose phosphate | Managed vs Forest | 124 | 156 | 13247 | <0.001*** |
| Pentose phosphate | Recent vs Late | 120 | 124 | 8561 | 0.084 |
| Pentose phosphate | Recent vs Forest | 120 | 156 | 12388 | <0.001*** |
| Pentose phosphate | Late vs Forest | 124 | 156 | 11869 | <0.01** |
| Phosphonate and Phospinate metabolism | Managed vs Recent | 62 | 30 | 966 | 0.767 |
| Phosphonate and Phospinate metabolism | Managed vs Late | 62 | 62 | 2405 | 0.064 |
| Phosphonate and Phospinate metabolism | Managed vs Forest | 62 | 156 | 6608 | <0.001*** |
| Phosphonate and Phospinate metabolism | Recent vs Late | 30 | 62 | 1146 | 0.145 |
| Phosphonate and Phospinate metabolism | Recent vs Forest | 30 | 156 | 3121 | 0.019* |
| Phosphonate and Phospinate metabolism | Late vs Forest | 62 | 156 | 5698 | 0.121 |
| Purine metabolism | Managed vs Recent | 403 | 390 | 81653 | 0.341 |
| Purine metabolism | Managed vs Late | 403 | 403 | 96256 | <0.001*** |
| Purine metabolism | Managed vs Forest | 403 | 546 | 149903 | <0.001*** |
| Purine metabolism | Recent vs Late | 390 | 403 | 90216 | <0.001*** |
| Purine metabolism | Recent vs Forest | 390 | 546 | 142029 | <0.001*** |
| Purine metabolism | Late vs Forest | 403 | 546 | 134099 | <0.001*** |
| Pyrimidine metabolism | Managed vs Recent | 186 | 180 | 17243 | 0.619 |
| Pyrimidine metabolism | Managed vs Late | 186 | 217 | 23786 | <0.01** |
| Pyrimidine metabolism | Managed vs Forest | 186 | 312 | 38524 | <0.001*** |
| Pyrimidine metabolism | Recent vs Late | 180 | 217 | 22544 | 0.016 |
| Pyrimidine metabolism | Recent vs Forest | 180 | 312 | 36778 | <0.001*** |
| Pyrimidine metabolism | Late vs Forest | 217 | 312 | 40356 | <0.001*** |
| Pyruvate metabolism | Managed vs Recent | 62 | 60 | 1930 | 0.918 |
| Pyruvate metabolism | Managed vs Late | 62 | 31 | 1113 | 0.651 |
| Pyruvate metabolism | Managed vs Forest | 62 | 156 | 6348 | <0.01** |
| Pyruvate metabolism | Recent vs Late | 60 | 31 | 1019 | 0.918 |
| Pyruvate metabolism | Recent vs Forest | 60 | 156 | 5967 | <0.01** |
| Pyruvate metabolism | Late vs Forest | 31 | 156 | 2855 | 0.452 |
| *Bacteria* |  |  |  |  |  |
| Organic phosphoester hydrolysis | Managed vs Recent | 696 | 720 | 289929 | <0.001*** |
| Organic phosphoester hydrolysis | Managed vs Late | 696 | 846 | 337791 | <0.001*** |
| Organic phosphoester hydrolysis | Managed vs Forest | 696 | 846 | 366067 | <0.001*** |
| Organic phosphoester hydrolysis | Recent vs Late | 720 | 846 | 300541 | 0.652 |
| Organic phosphoester hydrolysis | Recent vs Forest | 720 | 846 | 324142 | 0.056 |
| Organic phosphoester hydrolysis | Late vs Forest | 846 | 846 | 386784 | 0.012* |
| Others | Managed vs Recent | 174 | 180 | 18914 | <0.01** |
| Others | Managed vs Late | 174 | 282 | 31016 | <0.001*** |
| Others | Managed vs Forest | 174 | 282 | 31069 | <0.001*** |
| Others | Recent vs Late | 180 | 282 | 26664 | 0.882 |
| Others | Recent vs Forest | 180 | 282 | 26849 | 0.882 |
| Others | Late vs Forest | 282 | 282 | 39966 | 0.916 |
| Oxidative phosphorylation | Managed vs Recent | 174 | 180 | 18896 | <0.01** |
| Oxidative phosphorylation | Managed vs Late | 174 | 188 | 20409 | <0.001*** |
| Oxidative phosphorylation | Managed vs Forest | 174 | 188 | 20402 | <0.001*** |
| Oxidative phosphorylation | Recent vs Late | 180 | 188 | 17372 | 1 |
| Oxidative phosphorylation | Recent vs Forest | 180 | 188 | 17598 | 1 |
| Oxidative phosphorylation | Late vs Forest | 188 | 188 | 18068 | 1 |
| Pentose phosphate | Managed vs Recent | 609 | 630 | 228634 | <0.001*** |
| Pentose phosphate | Managed vs Late | 609 | 658 | 239621 | <0.001*** |
| Pentose phosphate | Managed vs Forest | 609 | 658 | 244892 | <0.001*** |
| Pentose phosphate | Recent vs Late | 630 | 658 | 206345 | 1 |
| Pentose phosphate | Recent vs Forest | 630 | 658 | 209405 | 1 |
| Pentose phosphate | Late vs Forest | 658 | 658 | 222027 | 1 |
| Phosphonate and Phospinate metabolism | Managed vs Recent | 1218 | 1260 | 883455 | <0.001*** |
| Phosphonate and Phospinate metabolism | Managed vs Late | 1218 | 1316 | 914389 | <0.001*** |
| Phosphonate and Phospinate metabolism | Managed vs Forest | 1218 | 1316 | 975208 | <0.001*** |
| Phosphonate and Phospinate metabolism | Recent vs Late | 1260 | 1316 | 822509 | 0.728 |
| Phosphonate and Phospinate metabolism | Recent vs Forest | 1260 | 1316 | 881036 | 0.012* |
| Phosphonate and Phospinate metabolism | Late vs Forest | 1316 | 1316 | 923772 | <0.01** |
| Phosphotransferase | Managed vs Recent | 174 | 180 | 19047 | <0.01** |
| Phosphotransferase | Managed vs Late | 174 | 188 | 20656 | <0.001*** |
| Phosphotransferase | Managed vs Forest | 174 | 188 | 20794 | <0.001*** |
| Phosphotransferase | Recent vs Late | 180 | 188 | 17468 | 1 |
| Phosphotransferase | Recent vs Forest | 180 | 188 | 17604 | 1 |
| Phosphotransferase | Late vs Forest | 188 | 188 | 17610 | 1 |
| Purine metabolism | Managed vs Recent | 1566 | 1620 | 1529489 | <0.001*** |
| Purine metabolism | Managed vs Late | 1566 | 1692 | 1608826 | <0.001*** |
| Purine metabolism | Managed vs Forest | 1566 | 1692 | 1637444 | <0.001*** |
| Purine metabolism | Recent vs Late | 1620 | 1692 | 1366181 | 1 |
| Purine metabolism | Recent vs Forest | 1620 | 1692 | 1382725 | 1 |
| Purine metabolism | Late vs Forest | 1692 | 1692 | 1461560 | 0.867 |
| Pyrimidine metabolism | Managed vs Recent | 957 | 990 | 577315 | <0.001*** |
| Pyrimidine metabolism | Managed vs Late | 957 | 1034 | 611641 | <0.001*** |
| Pyrimidine metabolism | Managed vs Forest | 957 | 1034 | 611464 | <0.001*** |
| Pyrimidine metabolism | Recent vs Late | 990 | 1034 | 514984 | 1 |
| Pyrimidine metabolism | Recent vs Forest | 990 | 1034 | 511589 | 1 |
| Pyrimidine metabolism | Late vs Forest | 1034 | 1034 | 535196 | 1 |
| Pyruvate metabolism | Managed vs Recent | 348 | 360 | 77696 | <0.001*** |
| Pyruvate metabolism | Managed vs Late | 348 | 564 | 120349 | <0.001*** |
| Pyruvate metabolism | Managed vs Forest | 348 | 564 | 121867 | <0.001*** |
| Pyruvate metabolism | Recent vs Late | 360 | 564 | 100213 | 1 |
| Pyruvate metabolism | Recent vs Forest | 360 | 564 | 99728 | 1 |
| Pyruvate metabolism | Late vs Forest | 564 | 564 | 158916 | 1 |
| Transporters | Managed vs Recent | 1218 | 1530 | 1125598 | <0.001*** |
| Transporters | Managed vs Late | 1218 | 1598 | 1249132 | <0.001*** |
| Transporters | Managed vs Forest | 1218 | 1598 | 1186037 | <0.001*** |
| Transporters | Recent vs Late | 1530 | 1598 | 1306113 | <0.01** |
| Transporters | Recent vs Forest | 1530 | 1598 | 1219634 | 0.911 |
| Transporters | Late vs Forest | 1598 | 1598 | 1176125 | <0.001*** |
| Two component system | Managed vs Recent | 522 | 540 | 169979 | <0.001*** |
| Two component system | Managed vs Late | 522 | 564 | 177929 | <0.001*** |
| Two component system | Managed vs Forest | 522 | 564 | 181157 | <0.001*** |
| Two component system | Recent vs Late | 540 | 564 | 151855 | 1 |
| Two component system | Recent vs Forest | 540 | 564 | 153684 | 1 |
| Two component system | Late vs Forest | 564 | 564 | 161422 | 1 |

**Table S16**: Results from pairwise Wilcoxon Rank-Sum tests for differences in average niche overlap of N-cycling genes partitioned across pathways. *P-*values adjusted using Benjamin-Hochberg correction for multiple testing.

| ***Pathway*** | ***Contrast*** | ***n1*** | ***n2*** | ***statistic*** | ***p.adj*** |
| --- | --- | --- | --- | --- | --- |
| denitrification | Managed vs Recent | 174 | 125 | 10058 | 0.72 |
| denitrification | Managed vs Late | 174 | 135 | 11741 | 0.996 |
| denitrification | Managed vs Forest | 174 | 204 | 24625 | < 0.001*** |
| denitrification | Recent vs Late | 125 | 135 | 9150 | 0.72 |
| denitrification | Recent vs Forest | 125 | 204 | 17913 | < 0.001*** |
| denitrification | Late vs Forest | 135 | 204 | 18858 | < 0.001*** |
| nitrate reduction | Managed vs Recent | 319 | 275 | 40801 | 0.284 |
| nitrate reduction | Managed vs Late | 319 | 270 | 43546 | 0.815 |
| nitrate reduction | Managed vs Forest | 319 | 408 | 89711 | < 0.001*** |
| nitrate reduction | Recent vs Late | 275 | 270 | 40378 | 0.23 |
| nitrate reduction | Recent vs Forest | 275 | 408 | 79266 | < 0.001*** |
| nitrate reduction | Late vs Forest | 270 | 408 | 74760 | < 0.001*** |
| nitrification | Managed vs Recent | 232 | 200 | 21993 | 1 |
| nitrification | Managed vs Late | 232 | 216 | 24895 | 1 |
| nitrification | Managed vs Forest | 232 | 272 | 44943 | < 0.001*** |
| nitrification | Recent vs Late | 200 | 216 | 22298 | 1 |
| nitrification | Recent vs Forest | 200 | 272 | 38319 | < 0.001*** |
| nitrification | Late vs Forest | 216 | 272 | 40819 | < 0.001*** |
| nitrogen fixation | Managed vs Recent | 145 | 50 | 3992 | 0.861 |
| nitrogen fixation | Managed vs Late | 145 | 135 | 9886 | 0.885 |
| nitrogen fixation | Managed vs Forest | 145 | 306 | 30865 | < 0.001*** |
| nitrogen fixation | Recent vs Late | 50 | 135 | 3067 | 0.861 |
| nitrogen fixation | Recent vs Forest | 50 | 306 | 9950 | <0.01** |
| nitrogen fixation | Late vs Forest | 135 | 306 | 28500 | < 0.001*** |

**Table S17**: Differences in the composition of functional diversity, redundancy, and Simpson’s dominance of predicted bacterial metagenomes between successional stages. Results based on permutational multivariate tests (perMANOVA) with Bray-Curtis distances and 10^^4^ permutations

| Contrast | *Df* | *R^2^* | *F* | *p-value* |
| --- | --- | --- | --- | --- |
| *Carbon* |  |  |  |  |
| *Managed vs Recent* | 1 | 0.02 | 1.78 | 0.140 |
| *Managed vs Late* | 1 | 0.03 | 1.75 | 0.155 |
| *Managed vs Forest* | 1 | 0.24 | 47.4 | < 0.001*** |
| *Recent vs Late* | 1 | 0.01 | 0.01 | 0.927 |
| *Recent vs Forest* | 1 | 0.10 | 15.2 | < 0.001*** |
| *Late vs Forest* | 1 | 0.09 | 12.4 | < 0.001*** |
| *Phosphorus* |  |  |  |  |
| *Managed vs Recent* | 1 | 0.03 | 2.38 | 0.105 |
| *Managed vs Late* | 1 | 0.04 | 2.72 | 0.078 |
| *Managed vs Forest* | 1 | 0.46 | 129 | < 0.001*** |
| *Recent vs Late* | 1 | 0.01 | 0.09 | 0.828 |
| *Recent vs Forest* | 1 | 0.24 | 41.2 | < 0.001*** |
| *Late vs Forest* | 1 | 0.21 | 33.0 | < 0.001*** |
| *Nitrogen* |  |  |  |  |
| *Managed vs Recent* | 1 | 0.09 | 0.65 | 0.47 |
| *Managed vs Late* | 1 | 0.01 | 0.94 | 0.382 |
| *Managed vs Forest* | 1 | 0.40 | 103 | < 0.001*** |
| *Recent vs Late* | 1 | 0.01 | 0.04 | 0.887 |
| *Recent vs Forest* | 1 | 0.28 | 50.2 | < 0.001*** |
| *Late vs Forest* | 1 | 0.26 | 45.0 | < 0.001*** |

**Table S18**: Effect sizes (*Hedge’s g*) of substrate-induced respiration rates between grassland sites at differing stages of succession compared to their respective paired forest sites. Negative effect sizes indicate higher respiration rates in forest sites.

| ***Substrate*** | ***Contrast*** | ***Hedge's g*** | ***95% CI*** |
| --- | --- | --- | --- |
| Glucose | *Managed vs Forest* | -0.44 | -0.89 - 0 |
| Glucose | *Recent vs Forest* | -0.5 | -1.1 - 0.11 |
| Glucose | *Late vs Forest* | -0.15 | -0.79 - 0.49 |
| Glycine | *Managed vs Forest* | -0.66 | -1.11 - -0.2 |
| Glycine | *Recent vs Forest* | -0.43 | -1.03 - 0.17 |
| Glycine | *Late vs Forest* | -0.26 | -0.9 - 0.38 |
| Oxalic Acid | *Managed vs Forest* | -1.18 | -1.65 - -0.7 |
| Oxalic Acid | *Recent vs Forest* | -0.85 | -1.47 - -0.23 |
| Oxalic Acid | *Late vs Forest* | -0.83 | -1.49 - -0.15 |
| Autolyzed yeast | *Managed vs Forest* | -0.69 | -1.14 - -0.23 |
| Autolyzed yeast | *Recent vs Forest* | -0.48 | -1.08 - 0.13 |
| Autolyzed yeast | *Late vs Forest* | -0.3 | -0.94 - 0.34 |
| Lignin | *Managed vs Forest* | -0.66 | -1.11 - -0.2 |
| Lignin | *Recent vs Forest* | -0.83 | -1.45 - -0.21 |
| Lignin | *Late vs Forest* | -0.5 | -1.14 - 0.16 |
| Chitin | *Managed vs Forest* | -0.23 | -0.67 - 0.21 |
| Chitin | *Recent vs Forest* | 0.45 | -0.16 - 1.05 |
| Chitin | *Late vs Forest* | 0.00 | -0.64 - 0.64 |
